# cellSTAAR: Incorporating single-cell-sequencing-based functional data to boost power in rare variant association testing of non-coding regions

**DOI:** 10.1101/2025.04.23.650307

**Authors:** Eric Van Buren, Yi Zhang, Xihao Li, Margaret Sunitha Selvaraj, Zilin Li, Hufeng Zhou, Nicholette D. Palmer, Donna K Arnett, John Blangero, Eric Boerwinkle, Brian E. Cade, Jenna C. Carlson, April P. Carson, Yii-Der Ida Chen, Joanne Curran, Ravindranath Duggirala, Myriam Fornage, Nora Franceschini, Misa Graff, Charles Gu, Xiuqing Guo, Jiang He, Nancy Heard-Cosa, Lifang Hou, Yi-Jen Hung, Rita R Kalyani, Sharon L.R. Kardia, Eimear Kenny, Charles Kooperberg, Brian G Kral, Leslie Lange, Dan Levy, Changwei Li, Simin Liu, Donald Lloyd-Jones, Ruth J.F. Loos, Ani W. Manichaikul, Lisa Warsinger Martin, Rasika Mathias, Ryan Minster, Braxton D. Mitchell, Josyf C. Mychaleckyj, Take Naseri, Kari North, Jeff O’Connell, James A. Perry, Patricia A. Peyser, Bruce M. Psaty, Laura M. Raffield, Vasan S. Ramachandran, Susan Redline, Alex Reiner, Stephen S. Rich, Jennifer A. Smith, Brian Spitzer, Hua Tang, Kent D. Taylor, Russell Tracy, Satupa‘itea Viali, Lisa Yanek, Wei Zhao, NHLBI Trans-Omics for Precision Medicine (TOPMed) Consortium, Jerome I. Rotter, Gina M. Peloso, Pradeep Natarajan, Xihong Lin

## Abstract

Whole genome sequencing (WGS) studies have identified hundreds of millions of rare variants (RVs) and have enabled RV association tests (RVATs) of these variants with complex traits and diseases. Analysis of non-coding variants is challenged by the considerable variability in regulatory function which candidate Cis-Regulatory Elements (cCREs) exhibit across cell types. We propose cellSTAAR, which integrates WGS data with single-cell ATAC-seq data to capture variability in chromatin accessibility across cell types via the construction of cell-type-specific functional annotations and variant sets. To reflect the uncertainty in cCRE-gene linking, cellSTAAR also links cCREs to their target genes using an omnibus framework which aggregates results from a variety of popular linking approaches. We applied cellSTAAR on Freeze 8 (N = 60,000) of the NHLBI Trans-Omics for Precision Medicine (TOPMed) consortium data to four lipids phenotypes: LDL cholesterol, a binary variable corresponding to high LDL cholesterol, HDL cholesterol, and triglycerides. We also provide replication results for all four phenotypes using UK Biobank (N = 190,000). Evidence from simulation studies and our real data analysis demonstrates that cellSTAAR boosts power and improves interpretation of RVATs of cCREs.

---

Large-scale multi-ancestry Whole-Genome sequencing (WGS) studies and biobanks, such as the Trans-Omics for Precision Medicine Program (TOPMed) of the National Heart, Lung, and Blood Institute [1], UK Biobank (UKB) [2], All of Us [3], and the Genome Sequencing Program (GSP) of the National Human Genome Research Institute (NHGRI), have been increasingly used to study the genetic underpinnings of complex human diseases. Collectively, these datasets include millions of individuals and billions of genetic variants, the vast majority of which are rare (Minor Allele Frequency (MAF) < 1%), non-coding, and many of which have unknown functional roles in human health and disease. It is of substantial interest to study the associations of non-coding rare variants with quantitative traits and diseases in multi-ethnic WGS studies and biobanks [4].

Because single variant tests have limited power for association tests of rare variants (RVs) [5–7], a number of set-based RV Association Tests (RVATs) have been developed, including SKAT [8], burden [9], their combinations [10–12], and STAAR [13]. A number of methods in statistical genetics, including PolyFun [14] and PAINTOR [15] for genetic finemapping, S-LDSC [16] for heritability partitioning, and eCAVIAR [17] for colocalization have shown that integrating functional annotations improves performance and biological interpretation. STAAR incorporates multi-faceted functional annotations to extend the advantages of integrating functional annotation data to RVATs by upweighting variants more likely to be functional to boost statistical power. STAAR also accounts for population structure and relatedness by utilizing a generalized linear mixed model (GLMM) based approach. This is especially important for reliable statistical inference because WGS data and biobanks often contain many related samples from ancestrally diverse populations. The STAAR framework using GLMMs is computationally scalable in analyses of hundreds of thousands of multi-ethnic related samples through the use of ancestry Principal Components (PCs) to account for population structure and sparse Genetic Relatedness Matrices (GRMs) to account for relatedness [18]. Extensive numerical results have shown that STAAR improves discovery in RVATs by incorporating multiple functional annotations, including unidimensional summary scores called annotation principal components (aPCs) [19].

Significant challenges remain, however, in RVATs of non-coding regions [4], particularly in candidate cis-regulatory elements (cCREs) [20, 21] such as enhancers and promoters. It is often a significant challenge to predict which genomic regions represent cCREs and which elements act as enhancers in any given cell type [22]. Additionally, previous work has shown considerable cell-type-specific heterogeneity in chromatin accessibility and regulatory activity [23, 24] in cCREs, particularly in enhancer regions. Such variability in chromatin accessibility by cell type is often missed in bulk assays [24–26], which average over multiple different cell types and can therefore distort meaningful biological variation across cell types.

Analyses integrating single-cell epigenetic data into existing methods in statistical genetics, including the previously mentioned S-LDSC and PolyFun, have demonstrated that incorporating this variability by cell type into analyses can improve the biological understanding of complex diseases [27, 28]. Including this variability into RVATs of cCREs therefore has the potential to provide a much-needed [4, 13] power improvement by prioritizing variants in active genomic regions in the most relevant cell types and improve result interpretation. This power improvement is especially critical for non-coding regions because variants in non-coding regions often exhibit weaker associations with phenotypes than variants in coding regions [4, 13, 29].

Considerable uncertainty exists in the linking between cCREs and their target genes, especially for enhancer regions. A number of different approaches to link pre-defined cCREs, particularly enhancers, to their target genes are commonly used. First, distance-based linking of candidate elements continues to show sufficiently good performance as compared with more sophisticated computational methods [30] in many situations. Second, two newer computational approaches, the Activity-By-Contact (ABC) model [21, 31] and EpiMap [32] (both described in more detail below), combine promoter-enhancer contact data and epigenetic and transcriptomic data, respectively, to make regulatory element-gene linking predictions. Third, evidence from eQTL studies and 3D conformation approaches, available in resources like SCREEN [33], provides information about the relationship between genetic variation and gene expression and the chromatin-chromatin interactions between neighboring genetic regions, respectively. Collectively, each element-gene linking approach mentioned relies on different kinds of genomic data to make predictions of possible cCRE-gene links. These predictions can vary dramatically in some genomic regions but remain highly consistent in other regions. None of the existing methods provides universally more accurate element-gene linking than the others. To date, RVAT methods such as STAAR have not reflected the uncertainty in element-gene linking by utilizing multiple linking approaches to produce a robust association p-values for gene-specific regulatory elements and phenotypes.

Existing approaches which integrate Whole Genome Sequencing (WGS) data and single-cell data have focused on integrating GWAS common variant summary statistics with scRNA-seq data [34–37] or scATAC-seq data [38] to improve the biological interpretation of the disease/trait associated common variant findings. To our knowledge, no method exists which integrates single-cell data with WGS to perform rare variant association analysis. To fill this gap, we developed cellSTAAR as an extension of the STAAR framework with the goal of improving RVAT performance in non-coding regions by integrating single-cell data to create cell-type-specific variant functional annotations and variant sets and accounting for element-gene linking uncertainty.

cellSTAAR boosts statistical power and aids in biological interpretation of significant loci through the ability to (1) integrate single-cell-sequencing-based functional annotations to weight genetic variants based on their activity in a cell type of interest, (2) use single-cell-sequencing-based epigenetic data to remove genomic regions with little activity in the cell type of interest and create custom cell-type-level variant sets, and (3) produce an omnibus p-value by aggregating multiple gene linking approaches to account for the uncertainty in element-gene linking across methods. To define enhancer and promoter regions, cellSTAAR uses version three of the catalog of cCREs from the ENCODE consortium [33]. This registry represents an expanded and updated collection of candidate regulatory elements as compared to the definitions used in other RVATs, such as the intersection of Genehancer [39] with CAGE [40, 41] or DHS [42] sites used in STAAR. The cellSTAAR framework is flexible in that it can accommodate the ever-increasing collection of single-cell data being generated through large-scale efforts such as the IGVF consortium [43] and the Human Cell Atlas [44, 45].

We applied cellSTAAR to detect rare variant (RV) associations in enhancer and promoter categories as defined by the ENCODE consortium [33] in four lipids traits—low-density lipoprotein cholesterol (LDL-C), a binary measure corresponding to high LDL-C levels, high-density lipoprotein cholesterol (HDL-C), and triglycerides (TG). Freeze 8 of the NHLBI Trans-Omics for Precision Medicine (TOPMed) consortium (N approximately 60,000 with lipids trait data) was used for discovery and replication was performed using UK Biobank (N approximately 180,000 whole genome samples with lipids trait data). We also performed simulation studies designed to evaluate the type-I error and power of cellSTAAR as compared with other popular RVAT methods in a variety of scenarios.

## Results

### Overview of Methods

We propose cellSTAAR (**Figure 1**) as a new statistical method to incorporate single-cell epigenetic datasets into gene-centric rare variant association tests (RVATs) of candidate cis-regulatory elements (cCREs). cCREs definitions were used exactly as provided from the ENCODE consortium [33], with separate analysis being reported for three mutually-exclusive regulatory element categories: proximal enhancer-like signatures (pELS), distal enhancer-like signatures (dELS), and promoter-like signatures (PLS). cellSTAAR simultaneously (1) incorporates cell-type-specific variant functional annotation scores computed using single-cell-sequencing-based assays such as single-cell ATAC-seq to weight variants based on activity in an individual cell type (**Figure 1b**), (2) incorporates single-cell-sequencing-based epigenomic information to prioritize and filter active cCREs in relevant cell types for association testing through variant set definition (**Figure 1c**), and (3) combines multiple cCRE-gene linking approaches to account for the variability in cCRE-gene linking (**Figure 1d**, see the “Enhancer-Gene Linking” and “Promoter-Gene Linking” in the Methods section for full details).

**Figure 1:**
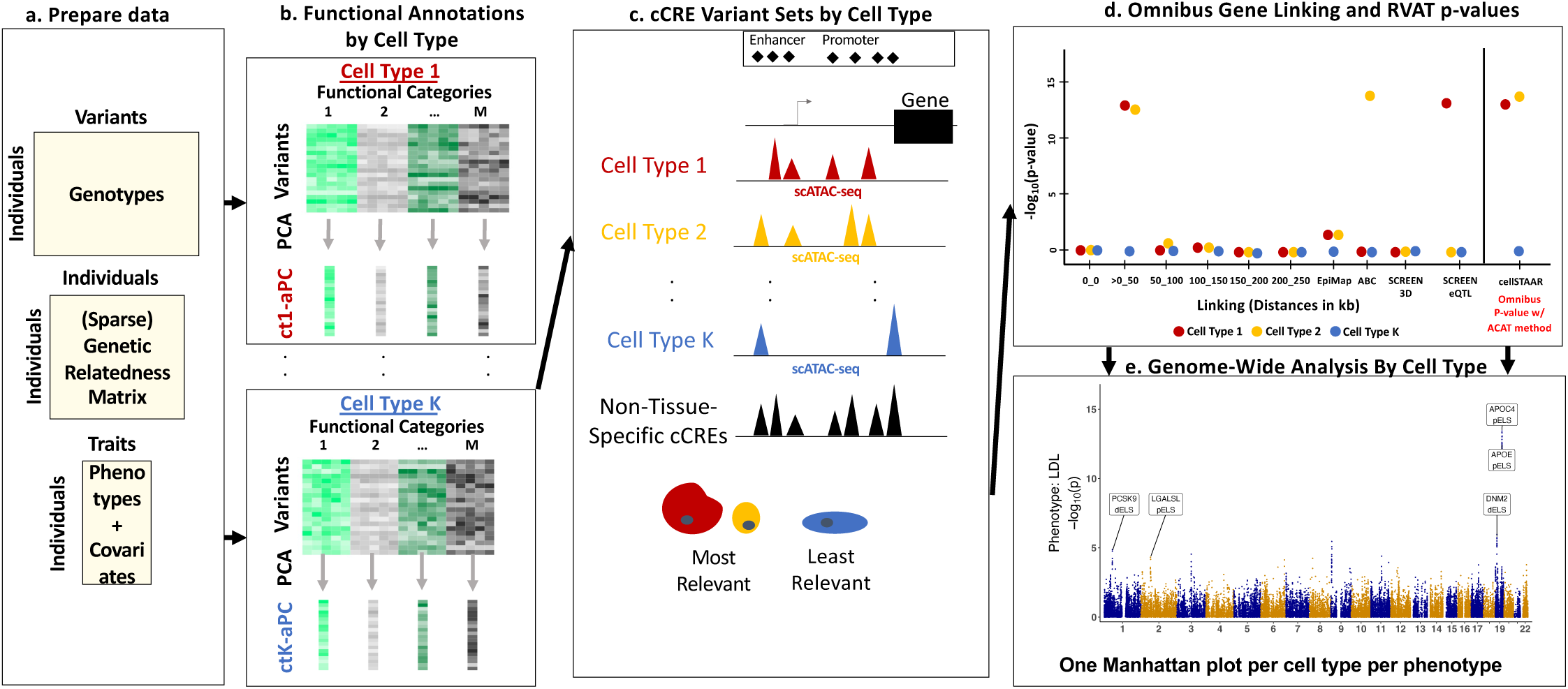
cellSTAAR overview. **a,** Input data of cellSTAAR, including genotypes, phenotype, covariates, (sparse) Genetic Relatedness Matrix (GRM), and external single-cell epigenetic data are prepared. **b,** Functional annotations are computed over each functional category for each of cell type being analyzed. **c,** scATAC-seq data are used to create gene-centric variant sets containing only regions with activity in a given cell type. **d,** The cellSTAAR omnibus p-value is obtained by aggregating results over a variety of different cCRE-gene linking approaches. **e,** For each cell type, genome-wide results are obtained.

Specifically, cellSTAAR incorporates single-cell-based variant functional annotations and employs a novel omnibus approach which uses the ACAT method [10] to aggregate results from a variety of linking approaches, each of which was chosen based on their overall performance and use of differing kinds of genomic data and computational techniques: (1) mutually-exclusive 50 kb distance-based intervals stretching to 250kb, weighted based on exponential decay [46] (see section “Element-gene linking and omnibus p-value calculation in cellSTAAR” below), (2) Activity-By-Contact (ABC, based on enhancer-promoter contacts), (3) EpiMap (based on the correlation of gene expression and epigenetic activity), and (4) SCREEN (both eQTL and 3D-based).

We demonstrate cellSTAAR on four lipids phenotypes, LDL-C, high LDL-C (binary), HDL-C, and TG, using Freeze 8 of the TOPMed consortium (N approximately 60,000) for discovery and UK Biobank (N approximately 180,000) for replication. In each case, we incorporate single-cell ATAC-seq data from the CATlas repository [23] generated from 19 cell types (see Lipids Phenotype Analysis section for the full list) representing a diversity of tissues. In our real data analysis of lipids phenotypes, we found that different methods often link the same region to multiple genes, and in simulation studies we found that by ensembling multiple linking approaches, cellSTAAR robustly boosts statistical power by maximizing the chances that a true linking is captured. We also found that the most discoveries are found in the most relevant cell types, such as those from liver and adipose tissues.

### Simulation Studies

We performed extensive simulation studies to evaluate the type-I error and power of cellSTAAR in a variety of settings designed to reflect realistic data scenarios (see ‘Data Simulation’ in the Methods section for full details of both type-I error and power simulations). Genotypes were simulated using the calibration coalescent model using a linkage disequilibrium (LD) structure representative of African-American populations [47]; a total of 20,000 sequences of 1Mb in length were simulated. Random 5kb regions were selected for testing based on 10,000 individuals, and simulations focused on aggregating rare and low frequency variants (MAF < 5%). Simulation scenarios include both relevant and irrelevant functional annotations, representing both single-cell-based and non-tissue-specific annotations.

### Type-I Error Simulations

Empirical type-I error rates for SKAT, burden, ACAT-V, STAAR, and cellSTAAR were evaluated. A total of 13 unassociated variant-level annotations were randomly simulated for each simulation replicate: 3 cell-type-specific annotations which represent categories which vary by cell type, 3 tissue-specific annotations which represent categories that vary by tissue, and 7 non-tissue-specific annotations which represent categories that do not vary across tissue. SKAT, burden, and ACAT-V were implemented using only minor allele frequency (MAF) as variant-level weights. STAAR was implemented using MAF, the three tissue-specific annotations and the 7 non-tissue-specific annotations as variant-level weights. cellSTAAR was implemented using MAF, the three cell-type-specific annotations, and the 7 non-tissue-specific annotations as variant-level weights.

SKAT, burden, ACAT-V, and STAAR test all variants in the 5kb regions. cellSTAAR combines results over four simulated linking approaches, each of which selects 70% of variants, to reflect the use of cell-type-level variant sets based on only regions with activity in the simulated cell type. Results based on 10^8^ simulation replicates show that the type-I error rate for cellSTAAR is particularly well controlled at smaller significance thresholds, but still retains nearly the nominal rate at all thresholds between 10^−3^ and 10^−6^ (Supplementary Table 1).

### Power Simulations

Empirical power, estimated as the proportion of p-values less than the 10^−7^ significance threshold, was compared between five methods: SKAT, burden, ACAT-V, STAAR, and cellSTAAR. Variant causality was assumed to be proportional to the MAF, with effect sizes dependent on simulated functional annotations. A total of 13 variant-level annotations were randomly simulated for each replicate: 3 cell-type-specific functional annotations which represent associated categories which exhibit true functional variability by cell type, 3 tissue-specific “polluted” annotations which represent the functional categories of the cell-type-specific annotations but with a signal diluted by the presence of multiple cell types in a given tissue, 2 non-tissue-specific functional annotations which represent associated categories that do not vary across tissue, and 5 irrelevant which represent uninformative categories.

SKAT, burden, and ACAT-V used only MAF as variant-level weights. STAAR was implemented using MAF, the 3 tissue-specific polluted annotations, the 2 non-tissue-specific functional annotations and the 5 irrelevant annotations as variant-level weights. cellSTAAR was implemented using MAF, the 3 cell-type-specific functional annotations, the 2 non-tissue-specific functional annotations and the 5 irrelevant annotations as variant-level weights. SKAT, burden, ACAT-V, and STAAR test all variants in the 5kb testing region. cellSTAAR combines results over four simulated linking approaches, each of which selects 90% of causal variants and 70% of non-causal variants, to reflect the use of cell-type-level variant sets based on only regions with activity in the simulated cell type.

Overall, results from our simulations suggest that cellSTAAR provides considerably higher power over rare variant association test (RVAT) approaches that do not incorporate cell-type specific functional annotations and improves power over STAAR-O through the use of multiple element-gene linking approaches and single-cell-based cell-type-specific annotations (**Supplementary Figure 1**). This result was consistent across combinations of the percentage of causal variants and the percentage of positive (vs. negative) effect sizes varying within (.05, .15, .35) and (100%, 80%, and 50%), respectively.

### Lipids Phenotype Analysis using TOPMed Freeze 8 and UK Biobank

We applied cellSTAAR to identify enhancer and promoter associations for three quantitative lipids traits, LDL-C (N = 61,861), HDL-C (N = 65,127), and TG (N = 63,848) levels, and one binary trait, high LDL-C (N = 26,604/61,861, defined as adjusted LDL-C > 130 mg/dL). Freeze 8 of the TOPMed WGS data was used for discovery and the most recent release of 200,000 samples with WGS data from UK Biobank for replication (N = 185,346 for LDL-C and 122,273/185,346 for highLDL-C, 174,380 for HDL-C, and 189,951 for TG). As was done previously [4], we applied rank-based inverse normal transformations to each continuous phenotype, including a log-transformation of TG levels, and adjusted for age, age squared, sex, self-reported race/ethnicity, sequencing center, and the first ten ancestral principal components. We additionally controlled for relatedness through heteroscedastic linear mixed models using sparse genetic relatedness matrices (GRMs) and study-race specific residual variance components. LDL-C measurements were adjusted for the presence of medications such as statins as has been done previously [29]. Results for the high LDL-C phenotype are shown in the supplementary files due to similarity with the continuous LDL-C phenotype, but still demonstrate cellSTAAR’s suitability to be applied to binary phenotypes.

Variants within a variant set were weighted in the STAAR framework (Methods) using cell-type-specific functional annotations (**Figure 1b**, see Methods) and additional annotations available in the FAVOR [19] database: MAF, three predictive scores (CADD, LINSIGHT, and FATHMM), and five non-tissue-specific aPCs (conservation, local diversity, mappability, transcription factor activity, and protein function). Enhancer and promoter definitions were taken from Version 3 of the ENCODE registry of candidate cis-regulatory elements (cCREs) [33] (more description in the Methods section). In total, we tested three kinds of variant sets, defined using ENCODE terminology: proximal and distal enhancer like signatures (“pELS” & “dELS”) and promoter like signatures (“PLS”).

Data from 19 cell types from a variety of tissues were used for analysis with cellSTAAR: hepatocytes, fetal hepatic endothelial cells, fetal hepatoblasts, and hepatic fibroblasts (liver), adipocytes (adipose), enterocytes (small intestine), atrial and ventral cardiomyocytes and cardiac fibroblasts (heart), alpha, beta, fetal islet, and delta/gamma islet cells (pancreas), microglia and astrocytes (brain), type I and II skeletal myocytes (muscle), and CD4 and Plasma B cells (blood).

Our goal was to select a comprehensive set of tissues and representative cell types including those known a priori to be highly relevant to lipids traits and some cell types which are likely less relevant to these traits. Although the number of cells used for each cell type did vary, we did not find substantial evidence that differences in results between cell types were attributable to sample size (**Supplementary Figure 2**). We chose to include data from the single cell ATAC-seq repository CATlas [23] because it is the most comprehensive single repository of cell types from major tissues available, including both fetal and adult samples. Relying on a single repository is preferred to multiple data sources because uniform processing allows us to make fairer comparisons between cell types and minimize concerns that differences between cell types are due to non-biological factors such as sequencing depth, batch effects, or data processing choices.

### cellSTAAR finds and replicates associations in non-coding regions

**Table 1** shows conditional results from cellSTAAR for the three continuous phenotypes by conditioning on known GWAS lipids-associated SNPs for significant combinations of gene, cCRE element class (as defined by ENCODE), and phenotype based on a Bonferroni-adjusted threshold of 3.01E-07 in the TOPMed discovery phase and with p-values of < .05 in the UKB replication phase (see **Supplementary Figures 3 and 4** for Manhattan plots and qq plots from the top cell type for each continuous phenotype, and **Supplementary Table 2 and Supplementary Figure 5** for results from analyzing the binary high LDL-C phenotype). The omnibus linking approach used by cellSTAAR accounts for variability in regulatory element-gene linking through non-mutually-exclusive mapping of associated regulatory elements to multiple genes within the same loci. Individual loci can then be analyzed in more detail (as in the next paragraph) to work towards an understanding of the “true” causal gene and the cell types in which the regulatory element is active.

**Table 1:** Lipids Discovery and Replication Results for Significant Loci. Shows significant, replicated conditional gene-element class-phenotype combinations from cellSTAAR for the three continuous lipids phenotypes (LDL-C, HDL-C, and TG) based on the Bonferroni threshold of .05/(18457 genes*3 element classes*4 phenotypes analyzed) = 2.26E-07 in the TOPMed discovery phase using the p-value combined over all 19 analyzed cell types and using a replication threshold of .05 in the UKB replication phase p-value combined over all 19 cell types. Results from the cell type with the minimum p-value in TOPMed and from STAAR using DHS sites to classify enhancers and promoters are also shown. Variants used in conditional analysis were chosen using the forward-selection based procedure described previously [3]. dELS: distal Enhancer-Like Signature, pELS: proximal Enhancer-Like Signature, PLS: Promoter-Like Signature

| trait | chr | gene | ENCODE element class | cellSTAAR p-values aggregated over 19 cell types |  | Cell type with minimum cellSTAAR p-value in TOPMed | cellSTAAR p-values from minimum TOPMed cell type |  | STAAR DHS p-value (no single-cell data used) |  | rsIDs used in conditional analysis |
| --- | --- | --- | --- | --- | --- | --- | --- | --- | --- | --- | --- |
|  |  |  |  | TOPMed | UKB |  | TOPMed | UKB | TOPMed | UKB |  |
| LDL-C | 19 | PVR | pELS | 7.47e-12 | 1.04e-04 | Enterocyte | 1.32e-12 | 8.85e-06 | 2.05e-02 | 6.88e-01 | rs429358, rs7412, rs12721054 |
| LDL-C | 19 | CEACAM19 | pELS | 3.11e-12 | 6.09e-05 | Type_I_Skeletal_Myocyte | 5.86e-13 | 1.40e-05 | 3.24e-03 | 9.80e-01 | "" |
| LDL-C | 19 | CEACAM16 | pELS | 8.19e-13 | 2.13e-06 | Adipocyte | 1.34e-13 | 8.22e-07 | 3.58e-02 | 5.52e-02 | "" |
| LDL-C | 19 | BCL3 | pELS | 3.60e-12 | 2.18e-05 | Adipocyte | 5.77e-13 | 4.43e-06 | 8.27e-01 | 3.28e-01 | "" |
| LDL-C | 19 | CBLC | pELS | 2.99e-12 | 7.05e-05 | Adipocyte | 8.67e-13 | 2.60e-05 | 7.25e-02 | 8.50e-06 | "" |
| LDL-C | 19 | BCAM | pELS | 2.32e-12 | 1.07e-04 | Adipocyte | 6.85e-13 | 3.82e-05 | 2.20e-05 | 8.24e-01 | "" |
| LDL-C | 19 | NECTIN2 | pELS | 9.59e-13 | 3.72e-05 | Adipocyte | 2.28e-13 | 8.60e-06 | 1.88e-01 | 1.03e-01 | "" |
| LDL-C | 19 | TOMM40 | pELS | 6.64e-13 | 2.53e-05 | Adipocyte | 1.51e-13 | 5.67e-06 | 9.74e-07 | 1.54e-01 | "" |
| LDL-C | 19 | APOE | pELS | 3.14e-13 | 8.06e-07 | Adipocyte | 5.13e-14 | 3.09e-07 | 7.77e-16 | 4.89e-07 | "" |
| LDL-C | 19 | APOC4 | pELS | 9.28e-14 | 2.57e-07 | Adipocyte | 1.60e-14 | 9.72e-08 | 7.14e-01 | 2.47e-01 | "" |
| LDL-C | 19 | APOC4-APOC2 | pELS | 9.28e-14 | 2.57e-07 | Adipocyte | 1.60e-14 | 9.72e-08 | 1.91e-01 | 9.64e-01 | "" |
| LDL-C | 19 | APOC2 | pELS | 1.44e-13 | 1.73e-06 | Adipocyte | 4.06e-14 | 5.98e-07 | 4.09e-02 | 1.06e-02 | "" |
| LDL-C | 19 | CLPTM1 | pELS | 7.31e-13 | 1.63e-05 | Adipocyte | 2.13e-13 | 8.40e-06 | 3.12e-03 | 2.06e-02 | "" |
| LDL-C | 19 | RELB | pELS | 8.57e-13 | 2.50e-05 | Adipocyte | 2.50e-13 | 1.03e-05 | 6.35e-03 | 2.38e-01 | "" |
| LDL-C | 19 | CLASRP | pELS | 5.21e-13 | 4.95e-05 | Adipocyte | 1.45e-13 | 2.20e-05 | 1.79e-01 | 5.12e-03 | "" |
| LDL-C | 19 | ZNF296 | pELS | 3.42e-12 | 1.29e-04 | Adipocyte | 1.00e-12 | 4.94e-05 | 3.43e-01 | 5.08e-01 | "" |
| LDL-C | 19 | GEMIN7 | pELS | 4.03e-12 | 1.52e-04 | Adipocyte | 1.18e-12 | 5.80e-05 | 7.56e-06 | 6.89e-05 | "" |
| LDL-C | 19 | PPP1R37 | pELS | 3.67e-12 | 1.38e-04 | Adipocyte | 9.60e-13 | 4.22e-05 | 1.85e-07 | 8.25e-04 | "" |
| LDL-C | 19 | NKPD1 | pELS | 9.47e-12 | 1.11e-04 | Adipocyte | 2.85e-12 | 3.24e-05 | 3.81e-04 | 7.46e-04 | "" |
| LDL-C | 19 | MARK4 | pELS | 5.76e-12 | 2.36e-04 | Type_I_Skeletal_Myocyte | 1.67e-12 | 6.36e-05 | 5.99e-03 | 1.19e-01 | "" |
| LDL-C | 19 | DMPK | pELS | 2.58e-13 | 6.00e-07 | Enterocyte | 7.72e-14 | 2.14e-07 | 2.49e-01 | 6.36e-02 | "" |
| HDL-C | 11 | BUD13 | dELS | 3.55e-34 | 1.89e-02 | Type_I_Skeletal_Myocyte | 8.69e-35 | 3.84e-02 | 1.03e-01 | 5.54e-03 | rs964184,<br>rs662799,<br>rs75919952,<br>rs525028,<br>rs11216230 |
| HDL-C | 11 | ZPR1 | dELS | 2.39e-34 | 3.61e-02 | Fetal_Endothelial_Hepatic | 6.08e-35 | 1.12e-02 | 7.22e-02 | 6.26e-05 | "" |
| HDL-C | 11 | APOA5 | dELS | 8.03e-34 | 3.38e-02 | Beta | 1.52e-34 | 2.82e-02 | 7.13e-01 | 1.20e-02 | "" |
| HDL-C | 11 | APOA1 | dELS | 5.11e-34 | 3.13e-02 | Beta | 1.15e-34 | 2.51e-02 | 4.94e-35 | 3.48e-10 | "" |
| HDL-C | 11 | SIK3 | dELS | 2.04e-33 | 4.33e-02 | Fetal_Endothelial_Hepatic | 4.99e-34 | 4.41e-02 | 1.49e-01 | 1.76e-03 | "" |
| HDL-C | 11 | SIDT2 | dELS | 7.37e-34 | 3.28e-02 | Fetal_Endothelial_Hepatic | 1.88e-34 | 4.62e-02 | 5.52e-01 | 2.08e-04 | "" |
| HDL-C | 16 | HERPUD1 | PLS | 3.95e-13 | 2.48e-09 | Enterocyte | 8.82e-14 | 1.44e-06 | 2.01e-03 | 8.88e-01 | rs9989419,<br>rs247617,<br>rs17231520,<br>rs12720922,<br>rs118146573,<br>rs5883,<br>rs1801706 |
| HDL-C | 16 | CETP | PLS | 7.89e-13 | 2.49e-09 | Enterocyte | 1.76e-13 | 2.88e-06 | 2.64e-06 | 4.35e-04 | "" |
| HDL-C | 16 | NLRC5 | PLS | 7.89e-13 | 2.49e-09 | Enterocyte | 1.76e-13 | 2.88e-06 | 7.23e-04 | 2.96e-03 | "" |
| TG | 11 | BUD13 | dELS | 6.07e-25 | 1.33e-08 | Type_I_Skeletal_Myocyte | 1.35e-25 | 1.31e-08 | 3.18e-02 | 3.38e-05 | rs28927680,<br>rs964184,<br>rs35120633,<br>rs2266788,<br>rs12287066,<br>rs662799,<br>rs9804646,<br>rs633389 |
| TG | 11 | ZPR1 | dELS | 3.94e-25 | 2.03e-08 | Fetal_Endothelial_Hepatic | 1.04e-25 | 1.74e-08 | 4.92e-06 | 3.25e-03 | "" |
| TG | 11 | APOA5 | dELS | 1.36e-24 | 1.17e-08 | Beta | 2.81e-25 | 9.00e-09 | 2.97e-01 | 1.40e-04 | "" |
| TG | 11 | APOA1 | dELS | 8.72e-25 | 2.46e-08 | Beta | 2.12e-25 | 2.34e-08 | 9.35e-26 | 3.86e-18 | "" |
| TG | 11 | SIK3 | dELS | 3.22e-24 | 2.20e-08 | Fetal_Endothelial_Hepatic | 8.01e-25 | 1.93e-08 | 1.45e-04 | 1.37e-08 | "" |
| TG | 11 | PAFAH1B2 | dELS | 2.87e-25 | 1.46e-08 | Alpha | 6.69e-26 | 1.54e-08 | 1.92e-03 | 9.86e-01 | "" |
| TG | 11 | SIDT2 | dELS | 1.14e-24 | 1.02e-08 | Fetal_Endothelial_Hepatic | 2.89e-25 | 1.34e-08 | 1.43e-01 | 1.77e-01 | "" |
| TG | 11 | TAGLN | dELS | 4.33e-25 | 1.58e-08 | Beta | 1.10e-25 | 1.56e-08 | 3.97e-02 | 7.38e-11 | "" |
| TG | 11 | PCSK7 | dELS | 1.17e-12 | 6.68e-08 | Beta | 9.41e-14 | 4.79e-08 | 8.48e-01 | 1.45e-01 | "" |
| TG | 11 | RNF214 | dELS | 9.45e-13 | 1.41e-08 | Beta | 7.54e-14 | 1.33e-08 | 5.00e-02 | 7.62e-02 | "" |
| TG | 11 | BACE1 | dELS | 5.66e-11 | 1.88e-08 | Delta_Gamma | 2.06e-11 | 1.91e-08 | 1.65e-02 | 1.09e-05 | "" |
| TG | 11 | CEP164 | dELS | 6.76e-10 | 1.02e-07 | Beta | 2.43e-10 | 6.13e-08 | 7.99e-01 | 9.13e-11 | "" |
| TG | 19 | MEF2B | dELS | 9.95e-08 | 3.31e-02 | Fetal_Endothelial_Hepatic | 1.69e-08 | 7.61e-02 | 9.53e-01 | 3.34e-01 | rs58542926,<br>rs8102280 |
| TG | 19 | BORCS8 | dELS | 8.37e-08 | 4.91e-02 | Fetal_Endothelial_Hepatic | 1.72e-08 | 9.58e-02 | 6.92e-02 | 8.03e-01 | "" |
| TG | 19 | HAPLN4 | dELS | 3.43e-08 | 1.95e-02 | Adipocyte | 5.90e-09 | 1.84e-02 | 9.16e-02 | 9.51e-01 | "" |
| TG | 19 | PVR | pELS | 4.36e-24 | 8.82e-03 | Enterocyte | 6.78e-25 | 1.16e-02 | 2.51e-02 | 9.06e-08 | rs429358,<br>rs7412,<br>rs438811,<br>rs12721054,<br>rs5112 |
| TG | 19 | CEACAM16 | pELS | 4.72e-25 | 1.44e-02 | Adipocyte | 8.79e-26 | 1.01e-02 | 1.86e-01 | 6.32e-02 | "" |
| TG | 19 | TOMM40 | pELS | 1.98e-26 | 1.59e-02 | Enterocyte | 4.76e-27 | 1.21e-02 | 1.35e-02 | 2.52e-02 | "" |
| TG | 19 | APOE | pELS | 1.81e-25 | 7.93e-03 | Adipocyte | 3.38e-26 | 6.39e-03 | 4.30e-23 | 2.70e-01 | "" |
| TG | 19 | APOC4 | pELS | 5.39e-26 | 1.08e-02 | Adipocyte | 1.04e-26 | 1.01e-02 | 5.59e-05 | 2.80e-01 | "" |
| TG | 19 | APOC4-APOC2 | pELS | 5.39e-26 | 3.94e-02 | Adipocyte | 1.04e-26 | 8.44e-02 | 5.65e-01 | 8.97e-01 | "" |
| TG | 19 | APOC2 | pELS | 8.36e-26 | 9.04e-03 | Adipocyte | 2.49e-26 | 8.06e-03 | 7.67e-01 | 9.78e-01 | "" |
| TG | 19 | RELB | pELS | 4.72e-25 | 3.05e-02 | Enterocyte | 1.40e-25 | 5.28e-02 | 1.40e-04 | 9.69e-01 | "" |
| TG | 19 | CLASRP | pELS | 1.24e-24 | 1.72e-02 | Enterocyte | 3.72e-25 | 1.35e-02 | 1.64e-01 | 7.11e-01 | "" |
| TG | 19 | MARK4 | pELS | 2.07e-24 | 4.16e-02 | Hepatocyte | 4.49e-25 | 2.97e-02 | 5.76e-02 | 8.27e-01 | "" |

After conditional analysis, cellSTAAR found and replicated significant element-gene pairs in loci which were previously seen in conditional rare variant association analyses of the TOPMed data using STAAR [29], including dELS elements linked to the *APOA1* locus for HDL-C and PLS elements linked to the *CETP* locus for HDL-C. cellSTAAR also revealed novel significant element-gene-phenotype combinations after conditional analysis in lipids-associated loci [29, 48, 49] for rare variant associations in pELS elements linked to the *APOE* locus for both LDL-C and TG and dELS elements linked to the *APOA1* locus for TG. Many additional genes with previously established [29, 48, 49] associations in non-coding regions with lipids phenotypes, including *LDLR, PCSK9, LPA,* and *APOB* have significant cellSTAAR p-values in the UKB replication phase only or when using a relaxed significance threshold of 1E-4 in either the discovery or replication phases (**Supplementary Tables 3 and 4 and Supplementary Figures 6 and 7**). This pattern of significance, in which many potential rare variant association discoveries in cCREs are only conditionally significant at a reduced threshold, was also seen in previous conditional analyses of the TOPMed and UKB WGS data using STAAR for non-coding association testing (see Supplementary Data 15 from [29]).

As compared to STAAR, results from cellSTAAR are more significant in the TOPMed discovery cohort for all three phenotypes and are also more significant in the UKB replication cohort for both LDL-C and TG. Replication p-values from cellSTAAR are more consistent and less random than STAAR in the analysis, particularly in analyzing HDL-C (for example, in comparing *APOA1* and *APOA5* using cellSTAAR and STAAR). STAAR tended to have more random replication p-values and failed to identify many of these associations and failed to replicate many discoveries found by cellSTAAR across all three phenotypes.

### cellSTAAR addresses uncertainty in element-gene linking and outperforms bulk data-based element-gene linking

**Figure 2** shows the variability in association for pELS variants in the *APOE* locus when analyzing LDL-C. Although the enhancer driving the association is contained with the *APOE* gene, which has been previously associated with LDL-C levels [29], 3D-based evidence from SCREEN suggests possible regulation of nearby genes *APOC2* and *APOC4.* The ability to demonstrate uncertainty in regulatory element to gene linking is not accounted for in existing RVAT methods such as STAAR. Further experimental validation of this locus may be required for a complete biological understanding of which variant(s) and genes are causal.

**Figure 2:**
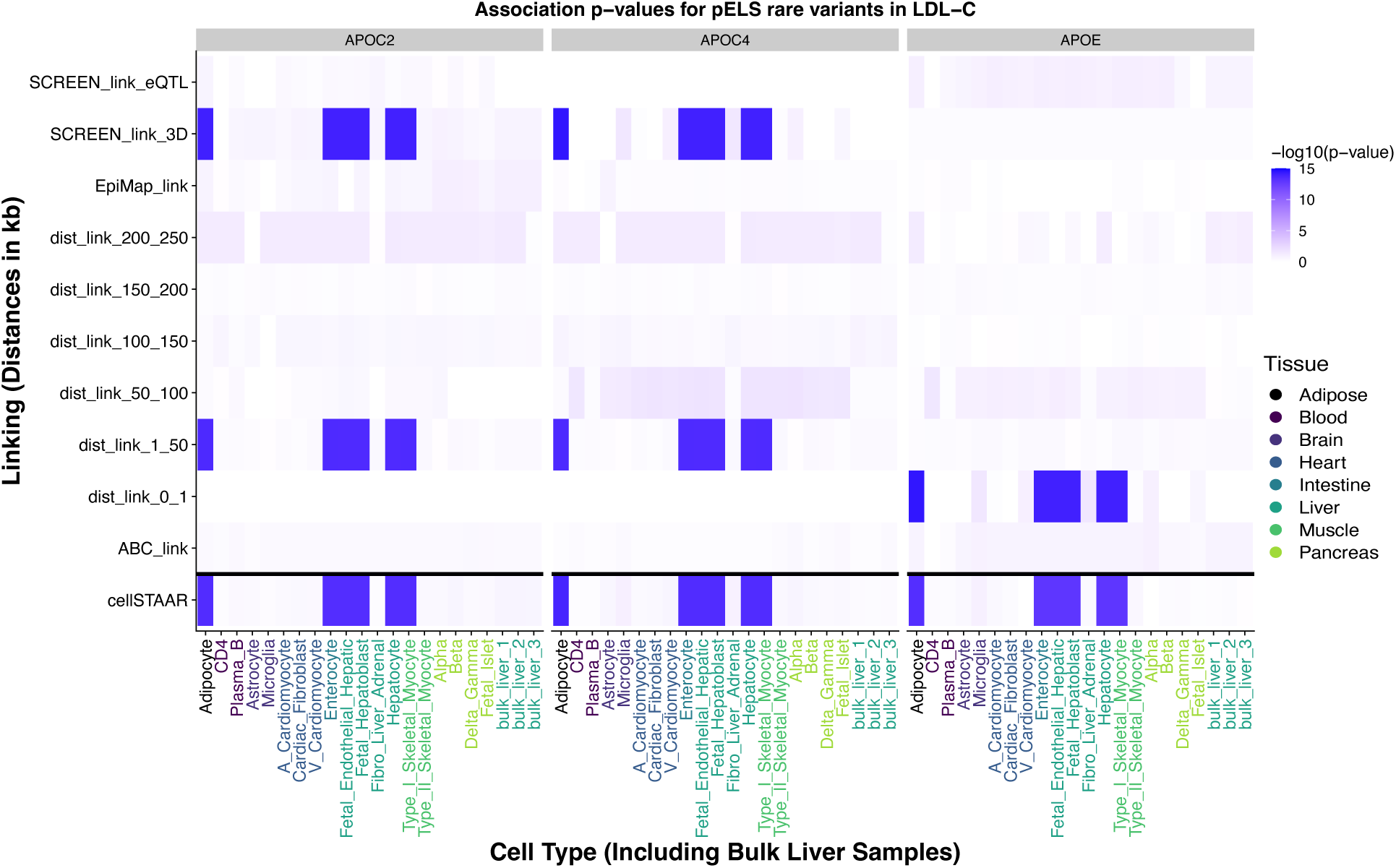
Variability in Association Across Linking Approaches and Cell Types (including 3 bulk comparisons from liver) for *APOC2, APOC4* and *APOE* in TOPMed Discovery Phase for LDL-C. For each gene, the first 19 columns correspond the cell-type-specific results using the scATAC-seq data, and the last three columns correspond to the bulk ATAC-seq-based results. These results show that significant associations are found by incorporating scATAC seq data, but no significant RVAT findings are found when incorporating bulk ATAC-seq data. pELS: proximal Enhancer-Like Signature.

Out of the 19 cell types analyzed, the significant pELS association is found when using data from 6 cell types, of which 5 are known a priori to be highly relevant to lipids: hepatocytes and fetal hepatoblasts [50–52], adipocytes [53], fetal liver endothelial cells [54], and enterocytes from the small intestine [55]. This suggests the possibility that the associated pELS regions are exhibiting a differential effect across cell types concentrated in cell types that are the most involved in regulating LDL-C levels. This association was missed completely when using bulk ATAC-seq data from three liver biosamples from ENCODE (see section “Use of ENCODE bulk liver ATAC-seq data**”** and columns bulk_liver_1, bulk_liver_2, and bulk_liver_3 in **Figure 2**), demonstrating one example in which the use of single-cell data provides improved discovery beyond the use of bulk data.

For LDL-C, results in the *APOE* locus for pELS elements, including significant cell types and the variation among linking approaches, were fully replicated using the UKB replication phase (**Supplementary Figure 8**). **Figure 2** shows that cellSTAAR can robustly capture both the differential regulation of associated enhancer regions across cell types and the uncertainty of the target gene(s) of regulation, collectively allowing for in-depth investigation of signals from RVATs involving cCREs. Although we did not typically find as much variation by cell type when testing PLS regions with cellSTAAR, a significant PLS association for HDL-C was found in the *CETP* gene with replication in some cell types in UKB (**Supplementary Figure 9**).

### cellSTAAR prioritizes cell types and increases discoveries

**Figure 3** shows the “percent enrichment” by cell type for the continuous lipids phenotypes, calculated by comparing the number of discoveries at a reduced significance threshold (10^−3^) with the average number within each phenotype over all 19 cell types analyzed. The x-axis is ordered by the total sum of percent enrichment over all three phenotypes, and therefore cell types on the left part of the x-axis show the most overall enrichment for lipids phenotypes. In the TOPMed discovery phase (**Figure 3a**), the top three cell types based on this enrichment were fetal hepatoblasts, fetal liver endothelial cells, and hepatocytes. All three are all non-immune cell types from liver, and all are expected *a priori* to be highly relevant to lipids levels. Conversely, the three least relevant cell types are microglial cells, fibroblasts from liver/adrenal gland, and astrocytes. None of these cell types are expected to be relevant for lipids levels.

**Figure 3:**
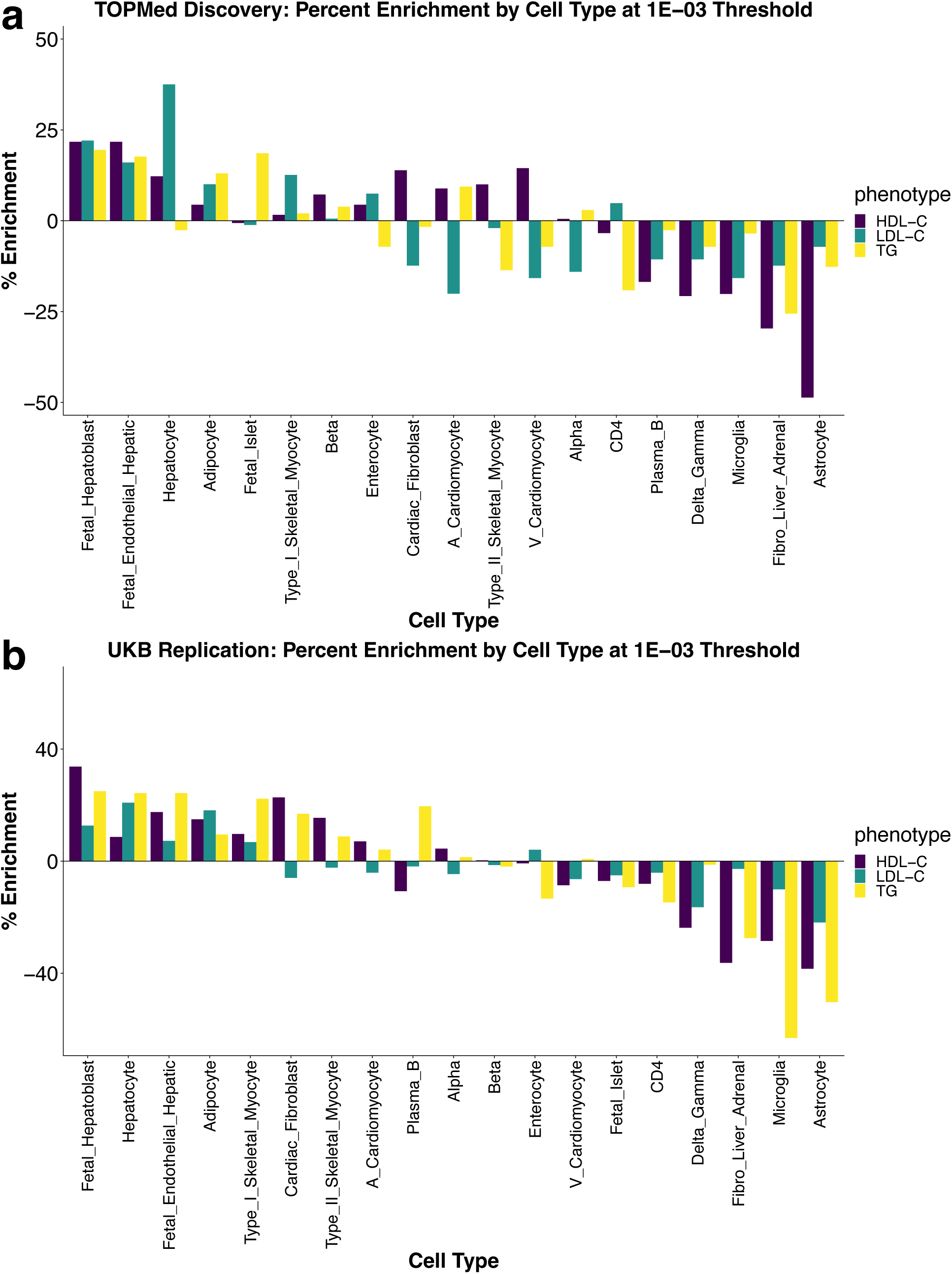
Percent enrichment by cell type. **a,** TOPMed discovery phase; **b,** UKB replication phase. Percent enrichment is calculated as the difference between the observed number of discoveries using cellSTAAR for a given cell type and the average number of discoveries over all 19 cell types within a given phenotype. The x-axis ranks cell types based on the sum of these percent enrichment values over all three continuous lipids phenotypes.

Results from analyzing the UKB replication phase (**Figure 3b**) broadly concur: the top three cell types from TOPMed rank 1, 3, and 2 using UKB. Overall, **Figure 3** provides evidence that incorporating chromatin accessibility measured at the cellular level can reveal additional discoveries when data from the most relevant cell types is used.

Visualizing the discoveries by phenotype instead of by cell type (**Supplementary Figure 10**) further reinforces the result that lipids-related cell types tended to show the most discoveries. This is also true when compared to STAAR [4, 13], which does not incorporate single-cell data (bars “STAAR_DHS” and “STAAR_CAGE”), when compared to three bulk liver ATAC-seq samples generated from ENCODE (bars “bulk_liver_1”, “bulk_liver_2”, and “bulk_liver_3”, see Methods for full details) and when comparing to an aggregated cellSTAAR p-value over all 19 analyzed cell types (“all_ct_ACAT”). Analyzing common variant associations across cell types in the TOPMed discovery phase revealed that the three liver cell types mentioned previously rank one, two, and four (**Supplementary Figures 11 and 12**) in the total number of discoveries. Collectively, these results suggest that incorporating single-cell-based chromatin accessibility data produces a larger number of more interpretable discoveries in more relevant cell types as compared to less relevant cell types.

### Integration of scRNA-seq data into cellSTAAR

Finally, in addition to scATAC-seq data, we further integrated scRNA-seq data from the Tabula Sapiens consortium [56] in cellSTAAR to investigate whether the scRNA-seq data can further boost the RVAT power. Because scRNA-seq data provides gene-level, and not variant-level information, it cannot be integrated to inform variant set construction or create variant-level functional annotations like scATAC-seq data. Rather, we used the “side-information” based approach IHW [57] to integrate scRNA-seq data in cellSTAAR. Briefly, IHW uses external covariates which are independent under the association null hypothesis to up or down weight the gene-centric cellSTAAR omnibus p-values (**Supplementary Figure 13**).

From Tabula Sapiens, we used data from 14 cell types which closely match 16 of the 19 cell types from which we used scATAC-seq data (**Supplementary Table 5**). For each cell type, we then computed measures of absolute expression (percentage of cells in a cell type expressing each gene) and differential expression (log fold change comparing each cell type to all others) using the FindMarkers function in the Seurat R package [58]. We found that the number of genome-wide discoveries was slightly reduced when using either absolute or differential gene expression as side-information covariates (**Supplementary Figure 14**). This is likely primarily driven by a weak relationship between cellSTAAR association p-values and either absolute or differential expression (**Supplementary Figures 15 and 16**). We also found limited success in using scRNA-seq data to provide a finer resolution to the significant association in *APOE* locus for pELS elements in the LDL-C phenotype (**Supplementary Figure 17**).

Cell types perform multiple biological functions, and highly expressed genes in relevant cell types may not have an association with lipids traits. Hence, using scRNA-seq-based cell-type-specific expression data to differentiate genes as gene-level weights may not increase significance in the cellSTAAR p-values. In the discussion section below, we expand on what additional data are needed to effectively integrate scRNA-seq data into cellSTAAR to boost the power of RVATs.

## Discussion

We developed a statistical method, cellSTAAR, to perform gene-centric non-coding rare variant association testing in cCREs by incorporating cell-type-specific variant functional annotations and variant sets based on single-cell data. cellSTAAR allows for the analysis of binary and continuous traits and adjusts for population structure and relatedness in a generalized linear mixed modelling framework. It is especially tailored to non-coding regions through: (1) the incorporation of single-cell ATAC-seq data to prioritize regions with accessible chromatin in a given cell type of interest, (2) an omnibus approach to account for uncertainty in linking cCREs to their target genes, and (3) the ability to incorporate cell-type-level functional annotations to upweight variants based on their activity in cell types of interest.

To perform unconditional and conditional genome-wide cellSTAAR analyses from 19 cell types for the HDL phenotype, analysis of the 65,127 samples from TOPMed took 14.41 hours using 100 computing cores for all 19 cell types including a maximum memory requirement of 41.52 GB (**Supplementary Table 6**). Analysis of the 174,380 samples from the UK Biobank using the Research Analysis Platform took 156.8 hours using 100 computing cores for all 19 cell types including a maximum memory usage of 62.07 GB.

Using the TOPMed WGS data, cellSTAAR revealed discoveries in the cCREs of several lipids-related genes, including *APOE*, *APOA1,* and *CETP*. In the *APOE* locus, which is known to be highly relevant to LDL-C levels, we replicated a result which showed that cellSTAAR is a robust RVAT that accounts for considerable variability between cell types and between element-gene linking approaches, which is not accounted for in existing methods such as STAAR but is crucial to consider for a more complete biological understanding. This locus shows that the uncertainty in element-gene linking is important to consider as the same associated region was plausibly linked to multiple genes, including *APOE*, *APOC2,* and *APOC4* using different element-gene linking approaches.

Fully resolving the uncertainty inherent in element-gene mapping in such loci remains a substantial challenge in determining casual variant to gene to phenotype pathways because there is no gold standard linkage approach. Use of the ACAT method to construct the cellSTAAR omnibus p-value provides a robust way to capture element-gene links that may be missed by a single, pre-chosen linking approach (**Supplementary Figure 18**).

In the coming years, large scale efforts like the IGVF consortium [59] will provide more reliable, validated genome-wide element-to-gene linkages based on multiome data and perturbation experiments and state-of-the-art computational approaches. These improved non-coding variant annotations and element-to-gene linkings can be incorporated in cellSTAAR to empower its performance for RVATs and improve result interpretation. The results of the cellSTAAR findings can also help prioritize experimental data generation by IGVF.

Using lipids phenotypes, we demonstrate that the incorporation of single-cell-based functional annotations and variant sets increases the number of non-coding rare variant discoveries of associations with traits and diseases and reveals relevant cell types based on the number of discoveries.

An important step in applying cellSTAAR is the choice of relevant cell types and tissues from which to prioritize single-cell data. For a good number of phenotypes, this choice is scientifically well understood. Other phenotypes may benefit from the ability to use complementary single-cell data to determine relevant cell types and tissues. In particular, one potential future methodological synergy is to use methods which link scRNA-seq data and GWAS results, such as sc-linker [34] and scDRS [35], to help reveal relevant cell types for association testing of a particular phenotype. Such an approach will benefit from the ever-increasing generation of single-cell sequencing data.

Future generation of genome-wide single-cell-sequencing datasets which include multiple cell types and tissues will increase the applicability of cellSTAAR in several ways. First, cellSTAAR can incorporate replicate datasets from a given cell type to reduce the noise that may be present in single-cell sequencing datasets, particularly in rare cell types. Second, new datasets representing different aspects of biological function can be incorporated. For example, datasets generated in the future will allow RVATs to prioritize genetic variants based on cell-type-specific chromatin interactions (single-cell Hi-C) or single-cell epigenetic mark activity (single-cell ChIP-seq).

Our efforts to integrate scRNA-seq data from the Tabula Sapiens consortium did not markedly improve rare variant association discoveries or biological interpretation at the genome-wide level. One likely reason is that cell types have multiple biological functions and, even in the most relevant cell types, the vast majority (>90%) of highly expressed genes were not found to be associated with lipids levels as seen in, for example, the GWAS catalog (p-value < 5E-8, **Supplementary Table 7**). A second reason is that there is an inconsistent relationship between gene expression and lipids association when ranking based on differential expression across cell types, even in 21 lipids Mendelian genes [60]. For example, *APOE* is implicated in both the GWAS catalog and cellSTAAR but is ranked 1,215 or worse in differential expression (**Supplementary Table 8**) in all cell types.

Taken together, these two facts limit the potential to differentiate genes or cell types based on only measures of scRNA-seq-based gene expression to boost the power of cellSTAAR. They also explain the weak relationship between either absolute and differential expression and the association p-value from cellSTAAR when analyzed genome-wide (**Supplementary Figures 15 and 16**).

Another contributing factor to the apparent lack of improvement from integrating scRNA-seq data is that the context of the Tabula Sapiens scRNA-seq data is not optimal for analyses of lipids traits. An ideal dataset would provide gene expression by cell type contrasting high lipids levels versus normal lipids levels. Associated lipids levels for the cells from Tabula Sapiens dataset are unknown, and to our knowledge there is not scRNA-seq study of expression changes due to high lipids levels collected over multiple different tissues. If such scRNA-seq data was available, using it to construct side-information weights applied to our lipids association p-values would likely improve the power of cellSTAAR.

The rapid expansion of sample sizes in WGS datasets and biobanks has led to the identification of hundreds of millions of non-coding rare variants. We introduce cellSTAAR, which provides a robust framework to perform association testing of these variants while accounting for the variability which exists across cell types in the non-coding genome. cellSTAAR therefore allows researchers to explore and quantify the heterogeneity in non-coding genome across multiple cell types and multiple regulatory element to gene linking approaches. As we demonstrated, cellSTAAR boosts statistical power and aids in more informative biological interpretation of rare variant associations in candidate cis-regulatory elements. Software to implement cellSTAAR is freely available as an R package at https://github.com/edvanburen/cellSTAAR.

## Supporting information

cellSTAAR Supplementary Tables

cellSTAAR Supplement

## Data Availability

This paper used the TOPMed Freeze 8 WGS data and lipids phenotype data. Genotype and phenotype data are both available in database of Genotypes and Phenotypes. The TOPMed WGS data were from the following twenty study phases (accession numbers provided in parentheses): Old Order Amish (phs000956.v1.p1), Atherosclerosis Risk in Communities Study (phs001211), Mt Sinai BioMe Biobank (phs001644), Coronary Artery Risk Development in Young Adults (phs001612), Cleveland Family Study (phs000954), Cardiovascular Health Study (phs001368), Diabetes Heart Study (phs001412), Framingham Heart Study (phs000974), Genetic Study of Atherosclerosis Risk (phs001218), Genetic Epidemiology Network of Arteriopathy (phs001345), Genetic Epidemiology Network of Salt Sensitivity (phs001217), Genetics of Lipid Lowering Drugs and Diet Network (phs001359), Hispanic Community Health Study - Study of Latinos (phs001395), Hypertension Genetic Epidemiology Network and Genetic Epidemiology Network of Arteriopathy (phs001293), Jackson Heart Study (phs000964), Multi-Ethnic Study of Atherosclerosis (phs001416), San Antonio Family Heart Study (phs001215), Genome-wide Association Study of Adiposity in Samoans (phs000972), Taiwan Study of Hypertension using Rare Variants (phs001387), and Women’s Health Initiative (phs001237). UKB WGS is available from the UKB RAP. The single-cell ATAC-seq used from CATlas is publicly available at http://catlas.org/humanenhancer/.

## Code Availability

cellSTAAR is freely available as an R package at https://github.com/edvanburen/cellSTAAR.

## Genome build

All genome coordinates are given in NCBI GRCh38/UCSC hg38.

## Acknowledgements

This work was supported by grants R35-CA197449, U19-CA203654, U01-HG012064, and U01-HG009088 (X. Lin), R01-HL142711 and R01-HL127564 (P.N. and G.M.P.), 75N92020D00001, HHSN268201500003I, N01-HC-95159, 75N92020D00005, N01-HC-95160, 75N92020D00002, N01-HC-95161, 75N92020D00003, N01-HC-95162, 75N92020D00006, N01-HC-95163, 75N92020D00004, N01-HC-95164, 75N92020D00007, N01-HC-95165, N01-HC-95166, N01-HC-95167, N01-HC-95168, N01-HC-95169, UL1-TR-000040, UL1-TR-001079, UL1-TR-001420, UL1-TR001881, DK063491, R01-HL071051, R01-HL071205, R01-HL071250, R01-HL071251, R01-HL071258, R01-HL071259, and UL1-RR033176 (J.R. and Y.C.), 1R35-HL135818, R01-HL113338, and HL046389 (S.R.), HL105756 (B.P.), HHSN268201600018C, HHSN268201600001C, HHSN268201600002C, HHSN268201600003C, and HHSN268201600004C (C.K.), R01-MD012765 and R01-DK117445 (N.F.), R01-HL153805, R03-HL154284 (B.E.C.), HHSN268201700001I, HHSN268201700002I, HHSN268201700003I, HHSN268201700005I, and HHSN268201700004I (E.B.), U01-HL072524, R01-HL104135-04S1, U01-HL054472, U01-HL054473, U01-HL054495, U01-HL054509, and R01-HL055673-18S1 (D.K.A.), U01-HL72518, HL087698, HL49762, HL59684, HL58625, HL071025, HL112064, NR0224103, M01-RR000052 (to the Johns Hopkins General Clinical Research Center), and NHLBI TOPMed Fellowship 75N92021F00229 (X.Li and M.S.S.). The Cardiovascular Health Study research was supported by NHLBI contracts HHSN268201200036C, HHSN268200800007C, HHSN268201800001C, N01HC55222, N01HC85079, N01HC85080, N01HC85081, N01HC85082, N01HC85083, N01HC85086, 75N92021D00006; and NHLBI grants U01HL080295, R01HL087652, R01HL105756, R01HL103612, R01HL120393, and U01HL130114 with additional contribution from the National Institute of Neurological Disorders and Stroke (NINDS). Additional support was provided through R01AG023629 from the National Institute on Aging (NIA). A full list of principal CHS investigators and institutions can be found at <u>CHS-NHLBI.</u> This work was also supported by R01-HL92301, R01-HL67348, R01-NS058700, R01-AR48797, R01-DK071891, R01-AG058921, the General Clinical Research Center of the Wake Forest University School of Medicine (M01-RR07122, F32 HL085989), the American Diabetes Association, and a pilot grant from the Claude Pepper Older Americans Independence Center of Wake Forest University Health Sciences (P60 AG10484). The Coronary Artery Risk Development in Young Adults Study (CARDIA) is conducted and supported by the National Heart, Lung, and Blood Institute (NHLBI) in collaboration with the University of Alabama at Birmingham (75N92023D00002 & 75N92023D00005), Northwestern University (75N92023D00004), University of Minnesota (75N92023D00006), and Kaiser Foundation Research Institute (75N92023D00003). The Framingham Heart Study (FHS) acknowledges the support of contracts NO1-HC-25195, HHSN268201500001I and 75N92019D00031 from the National Heart, Lung and Blood Institute and grant supplement R01 HL092577-06S1 for this research. We also acknowledge the dedication of the FHS study participants without whom this research would not be possible. R.S.V. is supported in part by the Evans Medical Foundation and the Jay and Louis Coffman Endowment from the Department of Medicine, Boston University School of Medicine. The Jackson Heart Study (JHS) is supported and conducted in collaboration with Jackson State University (HHSN268201800013I), Tougaloo College (HHSN268201800014I), the Mississippi State Department of Health (HHSN268201800015I) and the University of Mississippi Medical Center (HHSN268201800010I, HHSN268201800011I and HHSN268201800012I) contracts from the National Heart, Lung, and Blood Institute (NHLBI) and the National Institute on Minority Health and Health Disparities (NIMHD). The authors also wish to thank the staffs and participants of the JHS. Support for GENOA was provided by the National Heart, Lung, and Blood Institute (U01HL054457, U01HL054464, U01HL054481, R01HL119443, and R01HL087660) of the National Institutes of Health. Collection of the San Antonio Family Study data was supported in part by National Institutes of Health (NIH) grants P01 HL045522, MH078143, MH078111 and MH083824; and whole genome sequencing of SAFS subjects was supported by U01 DK085524 and R01 HL113323. The Diabetes Heart Study was supported by R01 HL92301, R01 HL67348, R01 NS058700, R01 AR48797, R01 DK071891, R01 AG058921, the General Clinical Research Center of the Wake Forest University School of Medicine (M01 RR07122, F32 HL085989), the American Diabetes Association, and a pilot grant from the Claude Pepper Older Americans Independence Center of Wake Forest University Health Sciences (P60 AG10484). Molecular data for the Trans-Omics in Precision Medicine (TOPMed) program was supported by the National Heart, Lung, and Blood Institute (NHLBI). Genome sequencing for “NHLBI TOPMed: Coronary Artery Risk Development in Young Adults (CARDIA)” (phs001612.v1.p1) was performed at the Baylor Sequencing Center (HHSN268201600033I). Core support including centralized genomic read mapping and genotype calling, along with variant quality metrics and filtering were provided by the TOPMed Informatics Research Center (3R01HL-117626-02S1; contract HHSN268201800002I). Core support including phenotype harmonization, data management, sample-identity QC, and general program coordination were provided by the TOPMed Data Coordinating Center (R01HL-120393; U01HL-120393; contract HHSN268201800001I). We gratefully acknowledge the studies and participants who provided biological samples and data for TOPMed. The full study specific acknowledgements are detailed in the **Supplementary Note**.

## Author Contributions

E.V.B., Y.Z., X. Li, Z. Li, and X. Lin designed the experiments. E.V.B., Y.Z., X. Li, and X. Lin performed the experiments. E.V.B., Y.Z., X. Li, Z. Li, H.Z., M.S.S., N.P.A, D.K.A., J.B., E.B., B.E.C., J.C.C, J.P.C, Y.D.I.C, J.C., R.D, M.F., N.F., M.G., C.G., X.G., J.H., N.H.C, L.H., Y.J.H, S.K., E.K., C.K., L.L., D.L., C.L., S.L., D.L.J., R.J.F.L., A.W.M, L.M.,R.M., R.M., B.M., J.C.M., T.N., K.N., J.O., J.P., P.P., B.P., L.R., V.S.R., S.R., A.R., S.S.R., J.S., B.S., H.T., K.D.T., R.T., S.V.,W.Z., J.R., G.M.P., P.N., and X. Lin acquired, analyzed, or interpreted data. J.R., G.M.P., P.N. and the NHLBI TOPMed Lipids Working Group provided administrative, technical, or material support. E.V.B. and X. Lin drafted the manuscript and revised it according to suggestions by the coauthors. All authors critically reviewed the manuscript, suggested revisions as needed and approved the final version.

## Competing Interests

E.K. has received personal fees from Regeneron Pharmaceuticals, 23&Me, Allelica, and Illumina; has received research funding from Allelica; and serves on the advisory boards for Encompass Biosciences, Overtone, and Galateo Bio. P.N. reports research grants from Allelica, Amgen, Apple, Boston Scientific, Genentech / Roche, and Novartis, personal fees from Allelica, Apple, AstraZeneca, Blackstone Life Sciences, Creative Education Concepts, CRISPR Therapeutics, Eli Lilly & Co, Foresite Labs, Genentech / Roche, GV, HeartFlow, Magnet Biomedicine, Merck, and Novartis, scientific advisory board membership of Esperion Therapeutics, Preciseli, and TenSixteen Bio, scientific co-founder of TenSixteen Bio, equity in MyOme, Preciseli, and TenSixteen Bio, and spousal employment at Vertex Pharmaceuticals, all unrelated to the present work. B.M.P. serves on the Steering Committee of the Yale Open Data Access Project funded by Johnson & Johnson. L.M.R. is a consultant for the TOPMed Administrative Coordinating Center (ACC) through Westat. X. Lin is a consultant of AbbVie Pharmaceuticals and Verily Life Sciences. The remaining authors declare no competing interests.

## NHLBI Trans-Omics for Precision Medicine (TOPMed) Consortium

Namiko Abe^53^, Gonçalo Abecasis^54^, Francois Aguet^55^, Christine Albert^56^, Laura Almasy^57^, Alvaro Alonso^58^, Seth Ament^59^, Peter Anderson^60^, Pramod Anugu^61^, Deborah Applebaum-Bowden^62^, Kristin Ardlie^55^, Dan Arking^63^, Allison Ashley-Koch^64^, Stella Aslibekyan^65^, Tim Assimes^66^, Paul Auer^67^, Dimitrios Avramopoulos^63^, Najib Ayas^68^, Adithya Balasubramanian^69^, John Barnard^70^, Kathleen Barnes^71^, R. Graham Barr^72^, Emily Barron-Casella^63^, Lucas Barwick^73^, Terri Beaty^63^, Gerald Beck^74^, Diane Becker^75^, Lewis Becker^63^, Rebecca Beer^76^, Amber Beitelshees^59^, Emelia Benjamin^77^, Takis Benos^78^, Marcos Bezerra^79^, Larry Bielak^54^, Joshua Bis^80^, Thomas Blackwell^54^, Nathan Blue^81^, Donald W. Bowden^82^, Russell Bowler^83^, Jennifer Brody^80^, Ulrich Broeckel^84^, Jai Broome^60^, Deborah Brown^85^, Karen Bunting^53^, Esteban Burchard^86^, Carlos Bustamante^87^, Erin Buth^88^, Jonathan Cardwell^89^, Vincent Carey^90^, Julie Carrier^91^, Cara Carty^92^, Richard Casaburi^93^, Juan P Casas Romero^94^, James Casella^63^, Peter Castaldi^95^, Mark Chaffin^55^, Christy Chang^59^, Yi-Cheng Chang^96^, Daniel Chasman^97^, Sameer Chavan^89^, Bo-Juen Chen^53^, Wei-Min Chen^98^, Michael Cho^90^, Seung Hoan Choi^55^, Lee-Ming Chuang^99^, Mina Chung^100^, Ren-Hua Chung^101^, Clary Clish^102^, Suzy Comhair^103^, Matthew Conomos^88^, Elaine Cornell^104^, Adolfo Correa^105^, Carolyn Crandall^93^, James Crapo^106^, L. Adrienne Cupples^107^, Jeffrey Curtis^108^, Brian Custer^109^, Coleen Damcott^59^, Dawood Darbar^110^, Sean David^111^, Colleen Davis^60^, Michelle Daya^89^, Michael DeBaun^112^, Dawn DeMeo^90^, Ranjan Deka^113^, Scott Devine^59^, Huyen Dinh^69^, Harsha Doddapaneni^114^, Qing Duan^115^, Shannon Dugan-Perez^116^, Jon Peter Durda^117^, Susan K. Dutcher^118^, Charles Eaton^119^, Lynette Ekunwe^61^, Adel El Boueiz^120^, Patrick Ellinor^121^, Leslie Emery^60^, Serpil Erzurum^122^, Charles Farber^98^, Jesse Farek^69^, Tasha Fingerlin^123^, Matthew Flickinger^54^, Chris Frazar^60^, Mao Fu^59^, Stephanie M. Fullerton^60^, Lucinda Fulton^124^, Stacey Gabriel^55^, Weiniu Gan^76^, Shanshan Gao^89^, Yan Gao^61^, Margery Gass^125^, Heather Geiger^126^, Bruce Gelb^127^, Mark Geraci^128^, Soren Germer^53^, Robert Gerszten^129^, Auyon Ghosh^90^, Richard Gibbs^69^, Chris Gignoux^66^, Mark Gladwin^130^, David Glahn^131^, Stephanie Gogarten^60^, Da-Wei Gong^59^, Harald Goring^132^, Sharon Graw^133^, Kathryn J. Gray^134^, Daniel Grine^89^, Colin Gross^54^, Yue Guan^59^, Namrata Gupta^135^, Jeff Haessler^125^, Michael Hall^136^, Yi Han^69^, Patrick Hanly^137^, Daniel Harris^138^, Nicola L. Hawley^139^, Ben Heavner^88^, Susan Heckbert^140^, Ryan Hernandez^86^, David Herrington^141^, Craig Hersh^142^, Bertha Hidalgo^65^, James Hixson^143^, Brian Hobbs^144^, John Hokanson^89^, Elliott Hong^59^, Karin Hoth^145^, Chao (Agnes) Hsiung^146^, Jianhong Hu^69^, Yi-Jen Hung^147^, Haley Huston^148^, Chii Min Hwu^149^, Marguerite Ryan Irvin^65^, Rebecca Jackson^150^, Deepti Jain^60^, Cashell Jaquish^151^, Jill Johnsen^152^, Andrew Johnson^76^, Craig Johnson^60^, Rich Johnston^58^, Kimberly Jones^63^, Hyun Min Kang^153^, Robert Kaplan^154^, Shannon Kelly^155^, Michael Kessler^59^, Alyna Khan^60^, Ziad Khan^69^, Wonji Kim^156^, John Kimoff^157^, Greg Kinney^158^, Barbara Konkle^159^, Holly Kramer^160^, Christoph Lange^161^, Ethan Lange^89^, Cathy Laurie^60^, Cecelia Laurie^60^, Meryl LeBoff^90^, Jonathon LeFaive^54^, Jiwon Lee^90^, Sandra Lee^69^, Wen-Jane Lee^149^, David Levine^60^, Joshua Lewis^59^, Xiaohui Li^162^, Yun Li^115^, Henry Lin^162^, Honghuang Lin^163^, Yongmei Liu^164^, Yu Liu^165^, Steven Lubitz^121^, Kathryn Lunetta^166^, James Luo^76^, Ulysses Magalang^167^, Michael Mahaney^168^, Barry Make^63^, Alisa Manning^169^, JoAnn Manson^90^, Melissa Marton^126^, Susan Mathai^89^, Susanne May^88^, Patrick McArdle^59^, Merry-Lynn McDonald^170^, Sean McFarland^156^, Stephen McGarvey^171^, Daniel McGoldrick^172^, Caitlin McHugh^88^, Becky McNeil^173^, Hao Mei^61^, James Meigs^174^, Vipin Menon^69^, Luisa Mestroni^133^, Ginger Metcalf^69^, Deborah A Meyers^175^, Emmanuel Mignot^176^, Julie Mikulla^76^, Nancy Min^61^, Mollie Minear^177^, Matt Moll^95^, Zeineen Momin^69^, May E. Montasser^59^, Courtney Montgomery^178^, Donna Muzny^69^, Girish Nadkarni^127^, Rakhi Naik^63^, Sergei Nekhai^179^, Sarah C. Nelson^88^, Bonnie Neltner^89^, Caitlin Nessner^69^, Deborah Nickerson^180^, Osuji Nkechinyere^69^, Tim O’Connor^59^, Heather Ochs-Balcom^181^, Geoffrey Okwuonu^69^, Allan Pack^182^, David T. Paik^183^, Nicholette Palmer^184^, James Pankow^185^, George Papanicolaou^76^, Cora Parker^186^, Juan Manuel Peralta^187^, Marco Perez^66^, Ulrike Peters^188^, Lawrence S Phillips^58^, Jacob Pleiness^54^, Toni Pollin^59^, Wendy Post^189^, Julia Powers Becker^190^, Meher Preethi Boorgula^89^, Michael Preuss^127^, Pankaj Qasba^76^, Dandi Qiao^90^, Zhaohui Qin^58^, Nicholas Rafaels^191^, Mahitha Rajendran^69^, D.C. Rao^124^, Laura Rasmussen-Torvik^192^, Aakrosh Ratan^98^, Robert Reed^59^, Catherine Reeves^193^, Elizabeth Regan^194^, Muagututi‘a Sefuiva Reupena^195^, Ken Rice^60^, Rebecca Robillard^196^, Nicolas Robine^126^, Dan Roden^197^, Carolina Roselli^55^, Ingo Ruczinski^63^, Alexi Runnels^126^, Pamela Russell^89^, Sarah Ruuska^198^, Kathleen Ryan^59^, Ester Cerdeira Sabino^199^, Danish Saleheen^200^, Shabnam Salimi^201^, Sejal Salvi^69^, Steven Salzberg^63^, Kevin Sandow^202^, Vijay G. Sankaran^203^, Jireh Santibanez^69^, Karen Schwander^124^, David Schwartz^89^, Frank Sciurba^130^, Christine Seidman^204^, Jonathan Seidman^205^, Vivien Sheehan^206^, Stephanie L. Sherman^207^, Amol Shetty^59^, Aniket Shetty^89^, Wayne Hui-Heng Sheu^149^, M. Benjamin Shoemaker^208^, Brian Silver^209^, Edwin Silverman^90^, Robert Skomro^210^, Albert Vernon Smith^211^, Josh Smith^60^, Nicholas Smith^212^, Tanja Smith^53^, Sylvia Smoller^154^, Beverly Snively^213^, Michael Snyder^214^, Tamar Sofer^129^, Nona Sotoodehnia^80^, Adrienne M. Stilp^60^, Garrett Storm^215^, Elizabeth Streeten^59^, Jessica Lasky Su^216^, Yun Ju Sung^124^, Jody Sylvia^90^, Adam Szpiro^60^, Frédéric Sériès^217^, Daniel Taliun^54^, Margaret Taub^63^, Matthew Taylor^133^, Simeon Taylor^59^, Marilyn Telen^64^, Timothy A. Thornton^60^, Machiko Threlkeld^218^, Lesley Tinker^219^, David Tirschwell^60^, Sarah Tishkoff^220^, Hemant Tiwari^221^, Catherine Tong^222^, Michael Tsai^185^, Dhananjay Vaidya^63^, David Van Den Berg^223^, Peter VandeHaar^54^, Scott Vrieze^185^, Tarik Walker^89^, Robert Wallace^145^, Avram Walts^89^, Fei Fei Wang^60^, Heming Wang^224^, Jiongming Wang^225^, Karol Watson^93^, Jennifer Watt^69^, Daniel E. Weeks^226^, Joshua Weinstock^153^, Bruce Weir^60^, Scott T Weiss^227^, Lu-Chen Weng^121^, Jennifer Wessel^228^, Cristen Willer^108^, Kayleen Williams^88^, L. Keoki Williams^229^, Scott Williams^230^, Carla Wilson^90^, James Wilson^231^, Lara Winterkorn^126^, Quenna Wong^60^, Baojun Wu^232^, Joseph Wu^183^, Huichun Xu^59^, Lisa Yanek^63^, Ivana Yang^89^, Ketian Yu^54^, Seyedeh Maryam Zekavat^55^, Yingze Zhang^233^, Snow Xueyan Zhao^106^, Xiaofeng Zhu^234^, Elad Ziv^235^, Michael Zody^53^, Sebastian Zoellner^54^, Mariza de Andrade^236^, Paul de Vries^237^, Lisa de las Fuentes^238^

53 - New York Genome Center, New York, New York, 10013, US; 54 - University of Michigan, Ann Arbor, Michigan, 48109, US; 55 - Broad Institute, Cambridge, Massachusetts, 2142, US; 56 Cedars Sinai, Boston, Massachusetts, 2114, US; 57 - Children’s Hospital of Philadelphia, University of Pennsylvania, Philadelphia, Pennsylvania, 19104, US; 58 - Emory University, Atlanta, Georgia, 30322, US; 59 - University of Maryland, Baltimore, Maryland, 21201, US; 61 - University of Mississippi, Jackson, Mississippi, 38677, US; 62 - National Institutes of Health, Bethesda, Maryland, 20892, US; 63 - Johns Hopkins University, Baltimore, Maryland, 21218, US; 64 - Duke University, Durham, North Carolina, 27708, US; 65 - University of Alabama, Birmingham, Alabama, 35487, US; 66 - Stanford University, Stanford, California, 94305, US; 67 - Medical College of Wisconsin, Milwaukee, Wisconsin, 53211, US; 68 - Providence Health Care, Medicine, Vancouver, CA; 69 - Baylor College of Medicine Human Genome Sequencing Center, Houston, Texas, 77030, US; 70 - Cleveland Clinic, Cleveland, Ohio, 44195, US; 71 - Tempus, University of Colorado Anschutz Medical Campus, Aurora, Colorado, 80045, US; 72 - Columbia University, New York, New York, 10032, US; 73 - The Emmes Corporation, LTRC, Rockville, Maryland, 20850, US; 74 - Cleveland Clinic, Quantitative Health Sciences, Cleveland, Ohio, 44195, US; 75 - Johns Hopkins University, Medicine, Baltimore, Maryland, 21218, US; 76 - National Heart, Lung, and Blood Institute, National Institutes of Health, Bethesda, Maryland, 20892, US; 77 - Boston University, Massachusetts General Hospital, Boston University School of Medicine, Boston, Massachusetts, 2114, US; 78 - University of Florida, Epidemiology, Gainesville, Florida, 32610, US; 79 - Fundação de Hematologia e Hemoterapia de Pernambuco - Hemope, Recife, 52011-000, BR; 80 - University of Washington, Cardiovascular Health Research Unit, Department of Medicine, Seattle, Washington, 98195, US; 81 - University of Utah, Obstetrics and Gynecology, Salt Lake City, Utah, 84132, US; 82 - Wake Forest University School of Medicine, Department of Biochemistry, Winston-Salem, North Carolina, 27157, US; 83 - National Jewish Health, National Jewish Health, Denver, Colorado, 80206, US; 84 - Medical College of Wisconsin, Pediatrics, Milwaukee, Wisconsin, 53226, US; 85 - University of Texas Health at Houston, Pediatrics, Houston, Texas, 77030, US; 86 - University of California, San Francisco, San Francisco, California, 94143, US; 87 - Stanford University, Biomedical Data Science, Stanford, California, 94305, US; 88 - University of Washington, Biostatistics, Seattle, Washington, 98195, US; 89 - University of Colorado at Denver, Denver, Colorado, 80204, US; 90 - Brigham & Women’s Hospital, Boston, Massachusetts, 2115, US; 91 - University of Montreal, US; 92 - Washington State University, Pullman, Washington, 99164, US; 93 - University of California, Los Angeles, Los Angeles, California, 90095, US; 94 - Brigham & Women’s Hospital, US; 95 - Brigham & Women’s Hospital, Medicine, Boston, Massachusetts, 2115, US; 96 - National Taiwan University, Taipei, 10617, TW; 97 - Brigham & Women’s Hospital, Division of Preventive Medicine, Boston, Massachusetts, 2215, US; 98 - University of Virginia, Charlottesville, Virginia, 22903, US; 99 - National Taiwan University, National Taiwan University Hospital, Taipei, 10617, TW; 100 - Cleveland Clinic, Cleveland Clinic, Cleveland, Ohio, 44195, US; 101 - National Health Research Institute Taiwan, Miaoli County, 350, TW; 102 - Broad Institute, Metabolomics Platform, Cambridge, Massachusetts, 2142, US; 103 - Cleveland Clinic, Immunity and Immunology, Cleveland, Ohio, 44195, US; 104 - University of Vermont, Burlington, Vermont, 5405, US; 105 - University of Mississippi, Population Health Science, Jackson, Mississippi, 39216, US; 106 - National Jewish Health, Denver, Colorado, 80206, US; 107 - Boston University, Biostatistics, Boston, Massachusetts, 2115, US; 108 - University of Michigan, Internal Medicine, Ann Arbor, Michigan, 48109, US; 109 - Vitalant Research Institute, San Francisco, California, 94118, US; 110 - University of Illinois at Chicago, Chicago, Illinois, 60607, US; 111 - University of Chicago, Chicago, Illinois, 60637, US; 112 - Vanderbilt University, Nashville, Tennessee, 37235, US; 113 - University of Cincinnati, Cincinnati, Ohio, 45220, US; 114 - Baylor College of Medicine Human Genome Sequencing Center, Houston, Texas, 77030; 115 - University of North Carolina, Chapel Hill, North Carolina, 27599, US; 116 - Baylor College of Medicine Human Genome Sequencing Center, BCM, Houston, Texas, 77030, US; 117 - University of Vermont, Pathology and Laboratory Medicine, Burlington, Vermont, 5405, US; 118 - Washington University in St Louis, Genetics, St Louis, Missouri, 63110, US; 119 - Brown University, Providence, Rhode Island, 2912, US; 120 - Harvard University, Channing Division of Network Medicine, Cambridge, Massachusetts, 2138, US; 121 - Massachusetts General Hospital, Boston, Massachusetts, 2114, US; 122 - Cleveland Clinic, Lerner Research Institute, Cleveland, Ohio, 44195, US; 123 - National Jewish Health, Center for Genes, Environment and Health, Denver, Colorado, 80206, US; 124 - Washington University in St Louis, St Louis, Missouri, 63130, US; 125 - Fred Hutchinson Cancer Research Center, Seattle, Washington, 98109, US; 126 - New York Genome Center, New York City, New York, 10013, US; 127 - Icahn School of Medicine at Mount Sinai, New York, New York, 10029, US; 128 - University of Pittsburgh, Pittsburgh, Pennsylvania, US; 129 - Beth Israel Deaconess Medical Center, Boston, Massachusetts, 2215, US; 130 - University of Pittsburgh, Pittsburgh, Pennsylvania, 15260, US; 131 - Boston Children’s Hospital, Harvard Medical School, Department of Psychiatry, Boston, Massachusetts, 2115, US; 132 - University of Texas Rio Grande Valley School of Medicine, San Antonio, Texas, 78229, US; 133 - University of Colorado Anschutz Medical Campus, Aurora, Colorado, 80045, US; 134 - Mass General Brigham, Obstetrics and Gynecology, Boston, Massachusetts, 2115, US; 135 - Broad Institute, Broad Institute, Cambridge, Massachusetts, 2142, US; 136 - University of Mississippi, Cardiology, Jackson, Mississippi, 39216, US; 137 - University of Calgary, Medicine, Calgary, CA; 138 - University of Maryland, Genetics, Philadelphia, Pennsylvania, 19104, US; 139 - Yale University, Department of Chronic Disease Epidemiology, New Haven, Connecticut, 6520, US; 140 - University of Washington, Epidemiology, Seattle, Washington, 98195-9458, US; 141 - Wake Forest Baptist Health, Winston-Salem, North Carolina, 27157, US; 142 - Brigham & Women’s Hospital, Channing Division of Network Medicine, Boston, Massachusetts, 2115, US; 143 - University of Texas Health at Houston, Houston, Texas, 77225, US; 144 - Regeneron Genetics Center, Boston, Massachusetts, 2115, US; 145 - University of Iowa, Iowa City, Iowa, 52242, US; 146 - National Health Research Institute Taiwan, Institute of Population Health Sciences, NHRI, Miaoli County, 350, TW; 147 - Tri-Service General Hospital National Defense Medical Center, TW; 148 - Blood Works Northwest, Seattle, Washington, 98104, US; 149 - Taichung Veterans General Hospital Taiwan, Taichung City, 407, TW; 150 - Oklahoma State University Medical Center, Internal Medicine, DIvision of Endocrinology, Diabetes and Metabolism, Columbus, Ohio, 43210, US; 151 - National Heart, Lung, and Blood Institute, National Institutes of Health, NHLBI, Bethesda, Maryland, 20892, US; 152 - University of Washington, Medicine, Seattle, Washington, 98109, US; 153 - University of Michigan, Biostatistics, Ann Arbor, Michigan, 48109, US; 154 - Albert Einstein College of Medicine, New York, New York, 10461, US; 155 - University of California, San Francisco, San Francisco, California, 94118, US; 156 - Harvard University, Cambridge, Massachusetts, 2138, US; 157 - McGill University, Montréal, QC H3A 0G4, CA; 158 - University of Colorado at Denver, Epidemiology, Aurora, Colorado, 80045, US; 159 - Blood Works Northwest, Medicine, Seattle, Washington, 98104, US; 160 - Loyola University, Public Health Sciences, Maywood, Illinois, 60153, US; 161 - Harvard School of Public Health, Biostats, Boston, Massachusetts, 2115, US; 162 - Lundquist Institute, Torrance, California, 90502, US; 163 - Boston University, University of Massachusetts Chan Medical School, Worcester, Massachusetts, 1655, US; 164 - Duke University, Cardiology, Durham, North Carolina, 27708, US; 165 - Stanford University, Cardiovascular Institute, Stanford, California, 94305, US; 166 - Boston University, Boston, Massachusetts, 2215, US; 167 - The Ohio State University, Division of Pulmonary, Critical Care and Sleep Medicine, Columbus, Ohio, 43210, US; 168 - University of Texas Rio Grande Valley School of Medicine, Brownsville, Texas, 78520, US; 169 - Broad Institute, Harvard University, Massachusetts General Hospital; 170 - University of Alabama, University of Alabama at Birmingham, Birmingham, Alabama, 35487, US; 171 - Brown University, Epidemiology, Providence, Rhode Island, 2912, US; 172 - University of Washington, Genome Sciences, Seattle, Washington, 98195, US; 173 - RTI International, US; 174 - Massachusetts General Hospital, Medicine, Boston, Massachusetts, 2114, US; 175 - University of Arizona, Tucson, Arizona, 85721, US; 176 - Stanford University, Center For Sleep Sciences and Medicine, Palo Alto, California, 94304, US; 177 - National Institute of Child Health and Human Development, National Institutes of Health, Bethesda, Maryland, 20892, US; 178 - Oklahoma Medical Research Foundation, Genes and Human Disease, Oklahoma City, Oklahoma, 73104, US; 179 - Howard University, Washington, District of Columbia, 20059, US; 180 - University of Washington, Department of Genome Sciences, Seattle, Washington, 98195, US; 181 - University at Buffalo, Buffalo, New York, 14260, US; 182 - University of Pennsylvania, Division of Sleep Medicine/Department of Medicine, Philadelphia, Pennsylvania, 19104-3403, US; 183 - Stanford University, Stanford Cardiovascular Institute, Stanford, California, 94305, US; 184 - Wake Forest Baptist Health, Biochemistry, Winston-Salem, North Carolina, 27157, US; 185 - University of Minnesota, Minneapolis, Minnesota, 55455, US; 186 - RTI International, Biostatistics and Epidemiology Division, Research Triangle Park, North Carolina, 27709-2194, US; 187 - University of Texas Rio Grande Valley School of Medicine, Edinburg, Texas, 78539, US; 188 - Fred Hutchinson Cancer Research Center, Fred Hutch and UW, Seattle, Washington, 98109, US; 189 - Johns Hopkins University, Cardiology/Medicine, Baltimore, Maryland, 21218, US; 190 - University of Colorado at Denver, Medicine, Denver, Colorado, 80204, US; 191 - University of Colorado at Denver, CCPM, Denver, Colorado, 80045, US; 192 - Northwestern University, Chicago, Illinois, 60208, US; 193 - New York Genome Center, New York Genome Center, New York City, New York, 10013, US; 194 - National Jewish Health, Medicine, Denver, Colorado, 80206, US; 195 - Lutia I Puava Ae Mapu I Fagalele, Apia, WS; 196 - University of Ottawa, Sleep Research Unit, University of Ottawa Institute for Mental Health Research, Ottawa, ON K1Z 7K4, CA; 197 - Vanderbilt University, Medicine, Pharmacology, Biomedicla Informatics, Nashville, Tennessee, 37235, US; 198 - University of Washington, Seattle, Washington, 98104, US; 199 - Universidade de Sao Paulo, Faculdade de Medicina, Sao Paulo, 1310000, BR; 200 - Columbia University, New York, New York, 10027, US; 201 - University of Maryland, Pathology, Seattle, Washington, 98195, US; 202 - Lundquist Institute, TGPS, Torrance, California, 90502, US; 203 - Harvard University, Division of Hematology/Oncology, Boston, Massachusetts, 2115, US; 204 - Harvard Medical School, Genetics, Boston, Massachusetts, 2115, US; 205 - Harvard Medical School, Boston, Massachusetts, 2115, US; 206 - Emory University, Pediatrics, Atlanta, Georgia, 30307, US; 207 - Emory University, Human Genetics, Atlanta, Georgia, 30322, US; 208 - Vanderbilt University, Medicine/Cardiology, Nashville, Tennessee, 37235, US; 209 - UMass Memorial Medical Center, Worcester, Massachusetts, 1655, US; 210 - University of Saskatchewan, Saskatoon, SK S7N 5C9, CA; 211 - University of Michigan; 212 - University of Washington, Epidemiology, Seattle, Washington, 98195, US; 213 - Wake Forest Baptist Health, Biostatistical Sciences, Winston-Salem, North Carolina, 27157, US; 214 - Stanford University, Genetics, Stanford, California, 94305, US; 215 - University of Colorado at Denver, Genomic Cardiology, Aurora, Colorado, 80045, US; 216 - Brigham & Women’s Hospital, Channing Department of Medicine, Boston, Massachusetts, 2115, US; 217 - Université Laval, Quebec City, G1V 0A6, CA; 218 - University of Washington, University of Washington, Department of Genome Sciences, Seattle, Washington, 98195, US; 219 - Fred Hutchinson Cancer Research Center, Cancer Prevention Division of Public Health Sciences, Seattle, Washington, 98109, US; 220 - University of Pennsylvania, Genetics, Philadelphia, Pennsylvania, 19104, US; 221 - University of Alabama, Biostatistics, Birmingham, Alabama, 35487, US; 222 - University of Washington, Department of Biostatistics, Seattle, Washington, 98195, US; 223 - University of Southern California, USC Methylation Characterization Center, University of Southern California, California, 90033, US; 224 - Brigham & Women’s Hospital, Mass General Brigham, Boston, Massachusetts, 2115, US; 225 - University of Michigan, US; 226 - University of Pittsburgh, Department of Human Genetics, Pittsburgh, Pennsylvania, 15260, US; 227 - Brigham & Women’s Hospital, Channing Division of Network Medicine, Department of Medicine, Boston, Massachusetts, 2115, US; 228 - Indiana University, Epidemiology, Indianapolis, Indiana, 46202, US; 229 - Henry Ford Health System, Detroit, Michigan, 48202, US; 230 - Case Western Reserve University; 231 - Beth Israel Deaconess Medical Center, Cardiology, Cambridge, Massachusetts, 2139, US; 232 - Henry Ford Health System, Department of Medicine, Detroit, Michigan, 48202, US; 233 - University of Pittsburgh, Medicine, Pittsburgh, Pennsylvania, 15260, US; 234 - Case Western Reserve University, Department of Population and Quantitative Health Sciences, Cleveland, Ohio, 44106, US; 235 - University of California, San Francisco, Medicine, San Francisco, California, 94143, US; 236 - Mayo Clinic, Health Quantitative Sciences Research, Rochester, Minnesota, 55905, US; 237 - University of Texas Health at Houston, Human Genetics Center, Department of Epidemiology, Human Genetics, and Environmental Sciences, Houston, Texas, 77030, US; 238 - Washington University in St Louis, Department of Medicine, Cardiovascular Division, St. Louis, Missouri, 63110, US

## Methods

### Ethics Statement

This study relied on analyses of genetic data from TOPMed phases. The study has been approved by the TOPMed Publications Committee, TOPMed Lipids Working Group and all the participating phases, including Old Order Amish (phs000956.v1.p1), Atherosclerosis Risk in Communities Study (phs001211), Mt Sinai BioMe Biobank (phs001644), Coronary Artery Risk Development in Young Adults (phs001612), Cleveland Family Study (phs000954), Cardiovascular Health Study (phs001368), Diabetes Heart Study (phs001412), Framingham Heart Study (phs000974), Genetic Study of Atherosclerosis Risk (phs001218), Genetic Epidemiology Network of Arteriopathy (phs001345), Genetic Epidemiology Network of Salt Sensitivity (phs001217), Genetics of Lipid Lowering Drugs and Diet Network (phs001359), Hispanic Community Health Study - Study of Latinos (phs001395), Hypertension Genetic Epidemiology Network and Genetic Epidemiology Network of Arteriopathy (phs001293), Jackson Heart Study (phs000964), Multi-Ethnic Study of Atherosclerosis (phs001416), San Antonio Family Heart Study (phs001215), Genome-wide Association Study of Adiposity in Samoans (phs000972), Taiwan Study of Hypertension using Rare Variants (phs001387), and Women’s Health Initiative (phs001237), where the accession numbers are provided in parenthesis. The use of human genetics data from TOPMed cohorts was approved by the Harvard T.H. Chan School of Public Health IRB (IRB13-0353).

### Use of CATlas single-cell ATAC-seq data in cellSTAAR

We used the CATlas repository (http://catlas.org/catlas_downloads/humantissues/) [23] as downloaded on February 17, 2022 to download single-cell ATAC-seq data from 19 representative cell types in both .bed (ATAC-seq peaks accessible in each cell type) and .bigwig (position-level scores for each cell type) formats. If a cell type has multiple samples, the cellSTAAR package averages the data by genomic position and weights all samples equally.

### cCRE Collection

We downloaded Version 3 of the ENCODE registry of candidate cis-regulatory elements (cCREs) from https://api.wenglab.org/screen_v13/fdownloads/V3/GRCh38-cCREs.bed on February 22, 2022. This registry contains a comprehensive list of cCREs classified into three categories described in more detail elsewhere [33]: Promoter-Like Signature (PLS), proximal Enhancer Like Signature (pELS), and distal Enhancer Like Signature (dELS). We did not modify category assignments or remove any regions belonging to these three categories from consideration.

### Use of ENCODE bulk liver ATAC-seq data

To compare performance when using bulk ATAC-seq data to performance using single-cell ATAC-seq data, on December 31, 2024, we downloaded ATAC-seq data from liver samples as available from the ENCODE consortium at https://www.encodeproject.org/matrix/?type=Experiment&searchTerm=liver+atac-seq. We downloaded both raw scores and peak calls from one sample from each of the three “tissues” listed at the above link: left lobe of liver (ENCFF318PKW (track), ENCFF188RKV (peak)), liver (ENCFF435JGG (track), ENCFF488BRH (peak)), and right lobe of liver (ENCFF295COO (track), ENCFF318SNW (peak)). Then, naming the respective samples as “bulk_liver_1”, “bulk_liver_2”, and “bulk_liver_3”, we used these samples to create functional annotations and variant sets using the *create_ct_annotations* and *create_cellSTAAR_mapping_file* functions in the *cellSTAAR* R package and run association analyses analogously to those which used the CATlas scATAC-seq data.

### Enhancer-Gene Linking

cellSTAAR combines results from five different approaches to link enhancer-like signatures (“pELS” and “dELS”) from V3 of the ENCODE registry to their target genes. First, we defined mutually-exclusive distance-based windows surrounding the transcription start and end sites for each gene as follows: [0,0] (contained within the gene), (0, 50] kb, (50, 100] kb, (100,150] kb, (150, 200] kb, and (200, 250] kb. Distance was calculated using the *gr.dist* function within the *gUtils R* package.

Second, we use all gene-enhancer links available on the SCREEN website as of February 22, 2022. Links are separated depending on whether they are from eQTL studies (approximately 70% of all links) or from either ChIA-PET, Hi-C data, or other 3D-structure-based assays (remaining 30% percent).

Third, we collected links provided from EpiMap [32], which uses the correlation between gene expression and imputed epigenomic activity. EpiMap links were first converted to hg38 using version 1.18.0 of the *liftOver* Bioconductor package. Then, we used the *findOverlaps* function in the *GenomicRanges* Bioconductor package to select ENCODE cCREs which have at least a 50bp overlap with a linked EpiMap enhancer, the collection of which was downloaded from https://personal.broadinstitute.org/cboix/epimap/links/pergroup/ on April 9, 2022.

Fourth, we used links from the ABC [21] method, which relies on chromatin activity and promoter-enhancer contact data to link candidate enhancer regions to genes. Once again, we used the *findOverlaps* function in the *GenomicRanges* Bioconductor package to find ENCODE cCREs which have at least 50bp overlap with the dataset linked ABC enhancer regions calculated on ENCODE biosamples, as downloaded from https://www.engreitzlab.org/resources on February 4, 2022. Because our aim is in evaluating the benefits of variant set construction and cell-type-specific functional annotations, and not a specific linking approach, we do not use tissue-specific or cell-type-specific links from ABC, EpiMap, and SCREEN. Any differences seen in association with lipids between two cell types or tissues is therefore not due to differences in linking. All enhancer-gene links for ENCODE V3 distal and proximal enhancer-like signature elements are available within the *cellSTAAR R* package (https://github.com/edvanburen/cellSTAAR) and used in the *create_cellSTAAR_mapping_file* function. See the “Element-Gene Linking in cellSTAAR” section below for additional details in mathematical notation.

### Promoter-Gene Linking

cellSTAAR combines results from three different approaches to link promoters (“PLS”) from V3 of the ENCODE registry to their target genes. First, as above, we define a single distance window of (0,4] kb using the *gr.dist* function from the *gUtils* R package. Second, we use all links available on SCREEN as of February 22, 2022, as discussed in more detail above. All promoter-gene links for ENCODE V3 promoter like signature elements are available within the *cellSTAAR R* package (https://github.com/edvanburen/cellSTAAR) and used in the *create_cellSTAAR_mapping_file* function. See the “Element-Gene Linking in cellSTAAR” section below for additional details in mathematical notation.

### Cauchy Combination Test and ACAT Method

ACAT [10, 61] is a flexible method to combine p-values of arbitrary dependence structure while preserving type-I error while increasing statistical power over alternative approaches. To combine M possibly dependent p-values, using weights ***w***_***m***_ such that ∑_***m***_ ***w***_***m***_ = **1**, ACAT uses the following approximation: 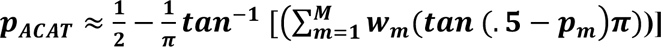. cellSTAAR employs the ACAT method to create a consensus p-value by combining p-values from differing linking approaches as shown in Figure 1.

### Conditional Analysis

We downloaded version 1.0 of the GWAS catalog on February 25, 2021. For each trait, we searched for associated variants by filtering based on the following trait matches: LDL-C (“LDL cholesterol”, “LDL cholesterol levels”, “Low density lipoprotein cholesterol levels”), HDL-C (“HDL cholesterol”, “HDL cholesterol levels”, “High density lipoprotein cholesterol levels”), TG (“Triglycerides”,“Triglyceride levels”,“Total triglycerides levels”). We then used the variant selection procedure described in our STAARpipeline paper [4] implemented in the *LD_pruning* function of the STAARpipeline *R* package to select a subset of variants that represent the signal in known variants from the GWAS catalog to adjust for in the conditional analyses.

### Notations and model

Suppose there are *n* subjects with *M* total variants sequenced across the whole genome. Given a genetic set of *p* variants, for subject *i*, let *Y*_*i*_ denote a continuous or dichotomous trait with mean *μi*; ***X****i* = (*Xi*1, …, *Xiq*)^*T*^ denote *q* covariates, such as age, gender, ancestral principal components; and ***G***_*i*_ = (*G*_*i*1_, …, *G*_*ip*_) denote the genotype information of the *p* genetic variants in a variant-set.

When the data consist of unrelated samples, we consider the following Generalized Linear Model (GLM)

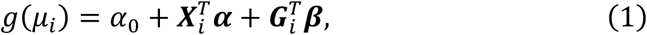

where *g*(*μ*) = *μ* for a continuous normally distributed trait, *g*(*μ*) = logit(*μ*) for a dichotomous trait, *α*_0_ is an intercept, ***α*** = (*α*_1_, …, *α*_*q*_)^*T*^ is a vector of regression coefficients for ***X***_*i*_, and ***β*** = (*β*_1_, …, *βp*)^*T*^ is a vector of regression coefficients for ***G****i*.

When the data consist of related samples, we consider the following Generalized Linear Mixed Model (GLMM) [62–64]

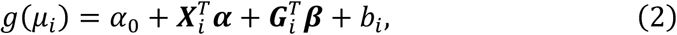

where the random effects *b*_*i*_ account for remaining population structure unaccounted by ancestral principal components (PCs), relatedness, and other between-observation correlation. We assume that 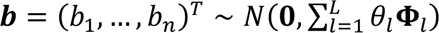 with variance components *θ*_*l*_ and known covariance matrices 𝚽_*l*_. The random effects ***b*** can be decomposed into a sum of multiple random effects to account for different sources of relatedness and correlation as 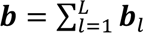 with ***b***_*l*_ ∼ 𝑁(𝟎, *θ*_*l*_𝚽_*l*_). For example, ***b***_1_ accounts for population structure and family relatedness by using the Genetic Relatedness Matrices (GRMs) as its covariance matrix 𝚽_1_ [65, 66]. A sparse GRM can be used to scale up computation [18]. Additional random effects ***b***_2_, ⋯, ***b***_𝐿_ can be used to account for complex sampling designs, such as correlation between repeated measures from longitudinal studies using subject-specific random intercepts and slopes and hierarchical designs. The remaining variables are defined in the same way as those in the GLM in equation (1). Under both the GLM and the GLMM, we are interested in testing the null hypothesis of whether the variant-set is associated with the phenotype, adjusting for covariates and relatedness, which corresponds to 𝐻_0_: ***β*** = 𝟎, that is, *β*_1_ = *β*_2_ = ⋯ = *β*_*p*_ = 0.

### cellSTAAR: Empowering non-coding rare variant association testing by incorporating single-cell-sequencing data

cellSTAAR improves the power of non-coding rare variant association testing through the following steps: (1) it incorporates cell-type-specific variant functional annotation scores computed using single-cell-sequencing-based assays such as single-cell ATAC-seq to weight variants based on activity in an individual cell type (Figure 1b), (2) it incorporates single-cell-sequencing-based epigenomic information to prioritize and filter active cCREs in relevant cell types for association testing through variant set definition (Figure 1c), and (3) it combines multiple cCRE-gene linking approaches to reflect the variability in cCRE-gene linking (Figure 1d).

### Rare variant association testing using cellSTAAR

cellSTAAR extends the STAAR framework to conduct non-coding gene-centric rare variant association testing by incorporating single-cell-based cell-type specific variant sets and functional annotations. Briefly, cellSTAAR calculates gene-centric p-values by integrating the burden, SKAT, and ACAT-V methods and incorporating single-cell based cell-type specific variant functional annotation scores [13] and the multi-faceted annotations used in the original STAAR method, for each combination of ENCODE cCRE element class and element-gene linking approach (discussed below). These functional annotation scores can be non-tissue-specific, e.g. as available in the FAVOR [19] database, or tissue/cell-type specific.

To create the single-cell-based cell-type specific variant functional annotations (**Figure 1b**) used in cellSTAAR, scores from the (possibly averaged) .bigwig files as discussed in the “Use of CATlas scATAC-seq data in cellSTAAR” section above are converted to the PHRED scale, defined as 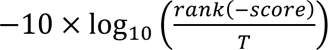, where *T* is the total number of variants sequenced across the whole genome. In this way, construction of the cell-type-level annotations does not depend on the absolute values of the peaks and the scATAC-seq peak calling approach or threshold, but only on the raw scores provided in the .bigwig files.

Suppose that, for the *m*th regulatory element to gene linkage method (*m* ∈ {*dist*_0–0𝑘*b*_, *dist*_0–4𝑘*b*_, *dist*_0–50𝑘*b*_, *dist*_50–100𝑘*b*_, *dist*_100–150𝑘*b*_, *dist*_150–200𝑘*b*_,, *dist*_200–250𝑘*b*_, *A*𝐵𝐶, 𝐸*piM*𝑎*p*, 𝑆𝐶𝑅𝐸𝐸𝑁_3𝐷_, 𝑆𝐶𝑅𝐸𝐸𝑁_𝑒Q*T*𝐿_}) (see their definitions in the Element-gene linking section below), there are 𝐾 functional annotations (including non-tissue-specific and single-cell-based). Define 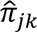 as the estimated probability of the *j*th variant being causal using the *k*th annotation (*k* = 0,…, *K*). We estimated 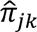 using the empirical cumulative distribution function of the *k*th annotation for variant *j* using its rank among all variants as:

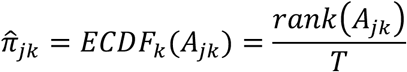

where *A*_j𝑘_ is the 𝑘th annotation for the 𝑗th variant (𝑘 = 1, ⋯, 𝐾; 𝑗 = 1, ⋯, *p*). For 𝑘 = 0, we assume 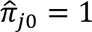. As was suggested previously [8, 13], we weight variants using two different beta densities 𝑤_j*l*_ = 𝐵𝑒𝑡𝑎(MAF_j_; 𝑎_1*l*_, 𝑎_2*l*_1, where (𝑎_11_, 𝑎_21_) = (1,25), (𝑎_12_, 𝑎_22_) = (1,1) (see the SKAT [8] and STAAR [13] papers for more discussion of these choices) and MAF_j_ is the MAF of the 𝑗th variant (𝑗 = 1, ⋯, *p*). The first density weights rarer variants more heavily than the second density, which weights variants equally based on MAF. To construct the cellSTAAR p-value, first define three different variant set statistics.

The burden test statistic using 𝑘th variant functional annotation and *l*th beta density (𝑘 = 0, ⋯, 𝐾; *l* = 1,2) to weight variants is given by

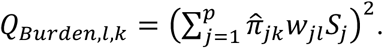

The SKAT test statistic using 𝑘th variant functional annotation and *l*th beta density to weight variants is given by

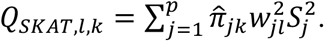

The ACAT-V test statistic using 𝑘th variant functional annotation and *l*th beta density to weight variants is given by

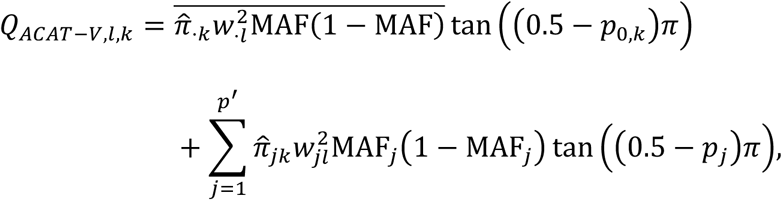

where 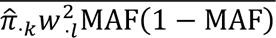 is the average of the weights 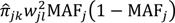 among the extremely rare variants with MAC ≤ 10, and *p*′ is the number of variants with MAC > 10 in the variant set.

For example, for the analyses presented in this paper, each statistic is based on the cell-type-specific annotations created using the CATlas scATAC-seq data and additional non-cell-type-specific annotations: MAF, three predictive scores (CADD, LINSIGHT, and FATHMM), and five non-tissue-specific aPCs (conservation, local diversity, mappability, transcription factor activity, and protein function).

Next, define the associated p-value for each of the previously defined statistics: *p*_𝐵𝑢𝑟𝑑𝑒*n*,*l*,𝑘_ is the *p*-value of 𝑄_𝐵𝑢𝑟𝑑𝑒*n*,*l*,𝑘_, *p*_𝑆𝐾*AT*,*l*,𝑘_ is the *p*-value of 𝑄_𝑆𝐾*AT*,*l*,𝑘_, and *p*_*A*𝐶*AT*–𝑉,*l*,𝑘_ is the *p*-value of 𝑄_*A*𝐶*AT*–𝑉,*l*,𝑘_ (𝑘 = 0, ⋯, 𝐾; *l* = 1,2).

Next, for each of the three methods SKAT, ACAT-V, and burden, the *p*-values are combined over all annotations *k* and both beta density weighting schemes l using the ACAT method: define STAAR-Burden (STAAR-B), STAAR-SKAT (STAAR-S), and STAAR-ACAT-V (STAAR-A) as 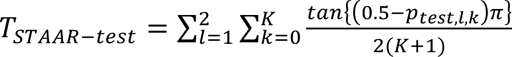, respectively. The corresponding *p*-value for each is calculated by 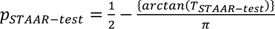, where 𝑡𝑒𝑠𝑡 ∈ {𝐵𝑢𝑟𝑑𝑒*n*, 𝑆𝐾*AT*, *A*𝐶*AT* − 𝑉}.

The omnibus test statistic used by cellSTAAR is constructed using the ACAT method as the equally weighted combination of the three *p*_𝑆*TAA*𝑅–𝑡𝑒𝑠𝑡_ *p*-values defined above:

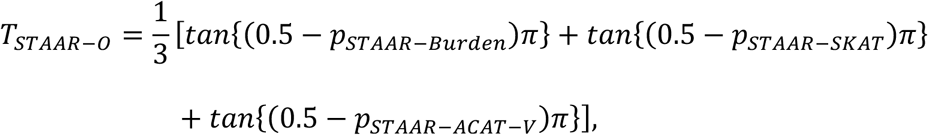

and the corresponding *p-*value to *T*_𝑆*TAA*𝑅–[_ is given by

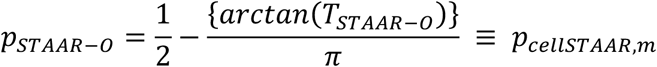

Each element-gene linking approach, indexed as (*m* = 1, ⋯, *M*), produces its own *p*_𝑐𝑒*ll*𝑆*TAA*𝑅,*m*_. cellSTAAR produces an omnibus p-value, *p*_𝑐𝑒*ll*𝑆*TAA*𝑅_, which aggregates over the *p*_𝑐𝑒*ll*𝑆*TAA*𝑅,*m*_’s as described in more detail in the section “Element-gene linking and omnibus p-value calculation in cellSTAAR below.

### Creation of cell-type-level variant sets

Variants were selected for inclusion in the cell-type level variant sets (**Figure 1c**) using the union of two criteria: (a) in a called scATAC-seq peak (as provided in the input data) for that cell type (.bed file) or (b) in the top 20% of non-zero scores for that cell type (.bigwig file). These criteria are not mutually-exclusive, but together ensure that we are capturing all peaks while also including flanking regions to provide robustness against the peak calling method and threshold used.

### Creation of cell-type-level functional annotations

Using the *create_ct_annotations* function of the cellSTAAR package, we created genome-wide PHRED-scaled annotation weights for each cell type using the raw track (.bigwig files) scATAC-seq data as provided by CATlas.

### Element-gene linking and omnibus p-value calculation in cellSTAAR

#### Enhancer regions

For testing proximal and distal enhancer like signatures (pELS/dELS) as defined by ENCODE), cellSTAAR aggregates multiple element-gene linking approaches, including distance-based (comprising six windows: 0-0 kb (contained within the gene), >0-50 kb, 50-100 kb, 100-150 kb, 150-200 kb, and 200-250 kb), ABC [21], EpiMap [32], SCREEN [33] eQTL-based, and SCREEN 3D-based, to perform rare variant association testing.

To define the distance-based p-value used to compute the cellSTAAR omnibus *p*-value, let 𝑑 = 1, …, 6 index the mutually exclusive distance-based windows used for element-gene linking, each of which has an overall association p-value given by *p*_𝑐𝑒*ll*𝑆*TAA*𝑅,𝑑_ as defined above. Then, the exponential-decay-based weighted distance-linking [46] p-value used in the omnibus cellSTAAR p-value calculation is computed using the ACAT method as:

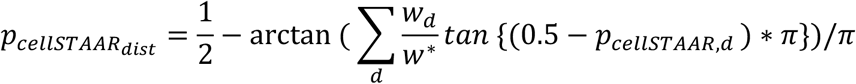

where the weights 𝑤_𝑑_ of each interval are given below and their sum is denoted by 𝑤^∗^(weights are standardized to ensure the sum of all weights used in the formula equals 1). In this way, cellSTAAR constructs an overall distance-based linking association *p*-value using distances ranging from 0 – 250 kb using the exponential distance based weight, which weights linkages which are closer in physical distance more heavily, and has been found to work well in the literature [46].

The cellSTAAR omnibus p-value is then calculated as:

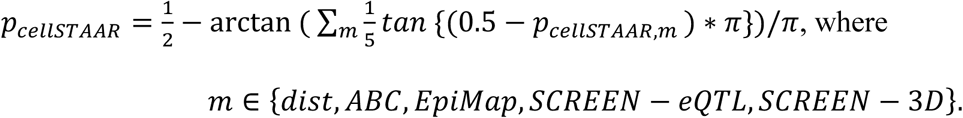

### Promoter regions

For testing promoter regions, cellSTAAR aggregates three element-gene linking approaches, including a 0-4 kb distance-based window, SCREEN eQTL-based, and SCREEN 3D-based, to perform rare variant association testing. Within a linking approach, cellSTAAR uses the STAAR-O p-value as defined above in its calculation. Define the individual association p-values for the three linking approaches as follows, *p*_𝑐𝑒*ll*𝑆*TAA*𝑅,*dist*_0_4𝑘*b*_ for the 0-4kb distance based window, *p*_𝑐𝑒*ll*𝑆*TAA*𝑅,𝑆𝐶𝑅𝐸𝐸𝑁–𝑒Q*T*𝐿_ for SCREEN eQTL-based, and *p*_𝑐𝑒*ll*𝑆*TAA*𝑅,𝑆𝐶𝑅𝐸𝐸𝑁–3𝐷_ for SCREEN 3D-based. Then, we define the cellSTAAR omnibus p-value as:

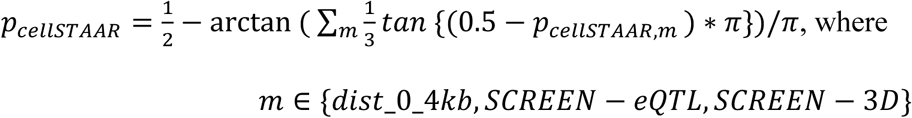

### Data Simulation

#### Type-I Error Simulations

We performed extensive simulation studies to evaluate the type-I error rate of cellSTAAR. Phenotypes were simulated under the null hypothesis from a linear model defined by

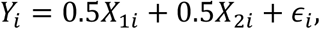

with *X*_1*i*_ ∼ 𝑁(0,1), *X*_2*i*_ ∼ Bernoulli(0.5), and 𝜖_*i*_ ∼ 𝑁(0,1). Genotypes were simulated as in STAAR [13]: we simulated 20,000 sequences for 100 different regions each spanning 1Mb from which randomly selected 5kb regions were used for type-I error testing. Data were generated to mimic the linkage disequilibrium (LD) structure of an African American population by using the calibration coalescent model (COSI) [47]. In each simulation replicate, a total of 13 unassociated annotations (*A*_1_, …, *A*_13_) were randomly simulated: 3 cell-type-specific (*A*_1_, *A*_2_, *A*_3_) which represent categories which vary by cell type, 3 tissue-specific (*A*_4_, *A*_5_, *A*_6_) which represent categories which vary at the tissue level, and 7 non-tissue-specific (*A*_7_, *A*_8_, *A*_9_, *A*_10_, *A*_11_, *A*_12_, *A*_13_) which represent categories that do not vary across tissue. Comparisons were made between ACAT, Burden, SKAT, STAAR, and cellSTAAR using 10^8^ replicates to examine type-I error at the following levels: *α* = 10^−3^, 10^−4^, 10^−5^, 10^−6^. STAAR was implemented using MAF, the tissue-specific annotations (*A*_4_, *A*_5_, *A*_6_) and the non-tissue-specific annotations (*A*_7_, *A*_8_, *A*_9_, *A*_10_, *A*_11_, *A*_12_, *A*_13_) as variant-level weights. cellSTAAR was implemented using MAF, the cell-type-specific annotations (*A*_1_, *A*_2_, *A*_3_) and the non-tissue-specific annotations (*A*_7_, *A*_8_, *A*_9_, *A*_10_, *A*_11_, *A*_12_, *A*_13_) as variant-level weights. ACAT, Burden, SKAT, and STAAR tested all variants in the 5kb testing window. For cellSTAAR, we simulated four different linking approaches each of which chooses 70% of all variants to reflect the use of cell-type-level variant sets which select only regions with activity in the given cell type. A sample size of 10,000 individuals was used.

### Empirical Power Simulations

Next, we carried out extensive simulations under a variety of alternative hypotheses to evaluate the power improvement from incorporating single-cell-based variant sets and functional annotations using cellSTAAR. As before, each simulation replicate used a randomly sampled 5kb genetic region from a global 1Mb regions. Each replicate also used different sets of analogously-simulated annotations designed to reflect differences in data source and functional relevance. A total of 13 annotations (*A*_1_, …, *A*_13_) were randomly simulated for each replicate: 3 cell-type-specific functional (*A*_1_, *A*_2_, *A*_3_) which represent categories which exhibit true functional variability by cell type, 3 tissue-specific polluted (*A*_4_, *A*_5_, *A*_6_) which represent the functional categories of (*A*_1_, *A*_2_, *A*_3_) but with a signal diluted by the presence of multiple cell types in a given tissue, 2 non-tissue-specific functional (*A*_7_, *A*_8_) which represent categories that do not vary across tissue, and 5 irrelevant (*A*_9_, *A*_10_, *A*_11_, *A*_12_, *A*_13_).

Variant-level (indexed by *j*) causal probabilities were simulated according to a logistic model defined by:

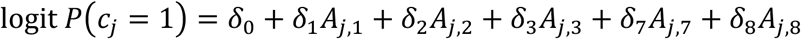

In this way we are assuming that causality depends on only 5 of the 13 annotations: 3 cell-type-specific functional annotations and 2 non-tissue-specific functional annotations. We set 𝛿_1_, 𝛿_2_, 𝛿_3_, 𝛿_7_, 𝛿_8_ = log (5) for all annotations as previously [13] and varied the proportions of causal variants in the signal region by setting 𝛿_0_ = logit(0.0015), logit(0.015), and logit(0.18) for averaging 5%, 15% and 35% causal variants in the signal region, respectively.

Phenotypes were generated from a linear model defined by:

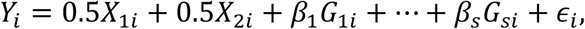

where *X*_1*i*_, *X*_2*i*_, 𝜖_*i*_ were defined identically as in the type-I error simulations, *G*_1*i*_, …, *G*_𝑠*i*_ were the genotypes of the *s* causal variants in the 5kb signal region, and *β*_1_, …, *β*_𝑠_ were the corresponding effect sizes of causal variants. These are modeled using an assumption that *β*_j_ = 𝛾_j_ = 𝑐_0_| log_10_ *MA*𝐹_j_ |, meaning that effect size is a decreasing function of MAF. c0 was set as 0.05 for all simulations. We additionally varied the direction of causal variant effect sizes by setting either 50%, 80%, or 100% of variants effects to be positive.

SKAT, Burden, and ACAT were implemented using only MAF as weights and including all variants in the 5kb testing region. STAAR was implemented including all variants in the 5kb testing region and using MAF and using the following 10 functional annotations as weights (as described above): 3 tissue-specific polluted (*A*_4_, *A*_5_, *A*_6_), 2 non-tissue-specific functional (*A*_7_, *A*_8_) and 5 irrelevant (*A*_9_, *A*_10_, *A*_11_, *A*_12_, *A*_13_). To capture the ability of cellSTAAR to use single-cell epigenomic data to create cell-type-level variant sets and use multiple element-gene linking approaches, we randomly selected 90% of causal variants and 70% of non-causal variants for inclusion into the cellSTAAR variant sets for each of four simulated linking approaches, as in the type-I error simulations. cellSTAAR used MAF as well as the following 10 annotations as weights (as described above): 3 cell-type-specific functional (*A*_1_, *A*_2_, *A*_3_), 2 non-tissue-specific functional (*A*_7_, *A*_8_), and 5 irrelevant (*A*_9_, *A*_10_, *A*_11_, *A*_12_, *A*_13_). To summarize, STAAR and cellSTAAR use 7 identical and 3 differing annotations and differ in the exact variant sets being tested and through cellSTAAR’s use of the omnibus element-gene linking p-value. A sample size of 10,000 individuals was used with 10^4^ replicates to evaluate power at the threshold. P-values used for each method are the same as in the table in the type-I error simulations description.

## Notes

https://github.com/edvanburen/cellSTAAR

### Competing Interest Statement

The authors have declared no competing interest.

