## Supplementary material for "cellSTAAR: Incorporating single-cell-sequencing-based functional data to boost power in rare variant association testing of non-coding regions": cellSTAAR Supplement

**Supplementary Figure 1**: Empirical power from simulation studies. Genotypes were simulated using the calibration coalescent model using an LD structure representative of African-American populations [1]; a total of 20,000 sequences of 1Mb in length were simulated. A total of 13 annotations ($A_1, \ldots, A_{13}$) were randomly simulated for each replicate: 3 cell-type-specific functional ($A_1, A_2, A_3$) which represent categories which exhibit true functional variability by cell type, 3 tissue-specific polluted ($A_4, A_5, A_6$) which represent the functional categories of ($A_1, A_2, A_3$) but with a signal diluted by the presence of multiple cell types in a given tissue, 2 non-tissue-specific functional ($A_7, A_8$) which represent categories that do not vary across tissue, and 5 irrelevant ($A_9, A_{10}, A_{11}, A_{12}, A_{13}$). SKAT, Burden, and ACAT included all variants in the randomly selected 5kb testing region and used only MAFs as weights. STAAR was implemented including all variants in the 5kb testing region and using MAF and the following 10 functional annotations as weights: 3 tissue-specific polluted ($A_4, A_5, A_6$), 2 non-tissue-specific functional ($A_7, A_8$) and 5 irrelevant ($A_9, A_{10}, A_{11}, A_{12}, A_{13}$). To capture the ability of cellSTAAR to use single-cell epigenomic data to create cell-type-specific variant sets, we randomly selected 90% of causal variants and 70% of non-causal variants for inclusion into the cellSTAAR variant sets for each of 4 simulated linking approaches. cellSTAAR used MAF as well as the following 10 annotations as weights: 3 cell-type-specific functional ($A_1, A_2, A_3$), 2 non-tissue-specific functional ($A_7, A_8$), and 5 irrelevant ($A_9, A_{10}, A_{11}, A_{12}, A_{13}$). Note that in this setup cellSTAAR and STAAR differ in (1) the variant sets being tested and (2) the use of 3 differing annotations as variant-level weights. Simulations were based on 10,000 individuals and $10^4$ simulation replicates.

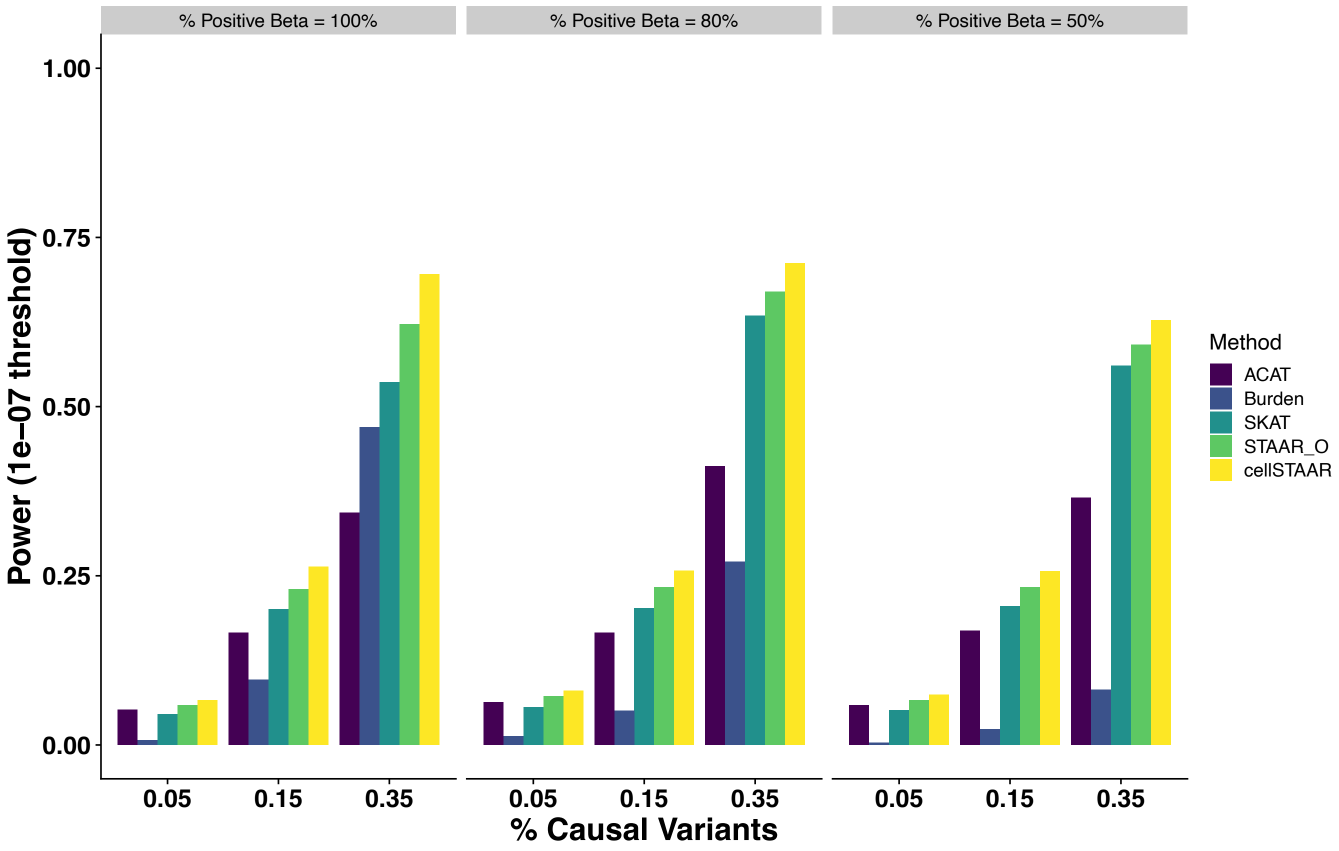

**Supplementary Figure 2**: Number of Discoveries using cellSTAAR (defined as p-value < $1x10^{-3}$) by CATlas nuclei count. **a,** TOPMed discovery phase; **b,** UKB Replication phase. Single-cell ATAC-seq data was used to select accessible genomic regions in each cell type by combining ATAC-seq peaks with regions in the top 20% of non-zero scores. Within the selected regions, rare variants within each of three ENCODE V3 element classes (dELS, pELS, PLS) were tested for association with the lipids phenotypes using cellSTAAR, which performs functionally-informed cell-type-level association testing using a generalized linear mixed model and multiple element-gene linking approaches, as described in the main text. Discoveries were then tallied for HDL-C, LDL-C, and TG in both the TOPMed discovery and UKB replication phases for the 19 cell types analyzed. The best linear fit along with the 95% confidence band is plotted and select cell types are labeled (most labels omitted for brevity).

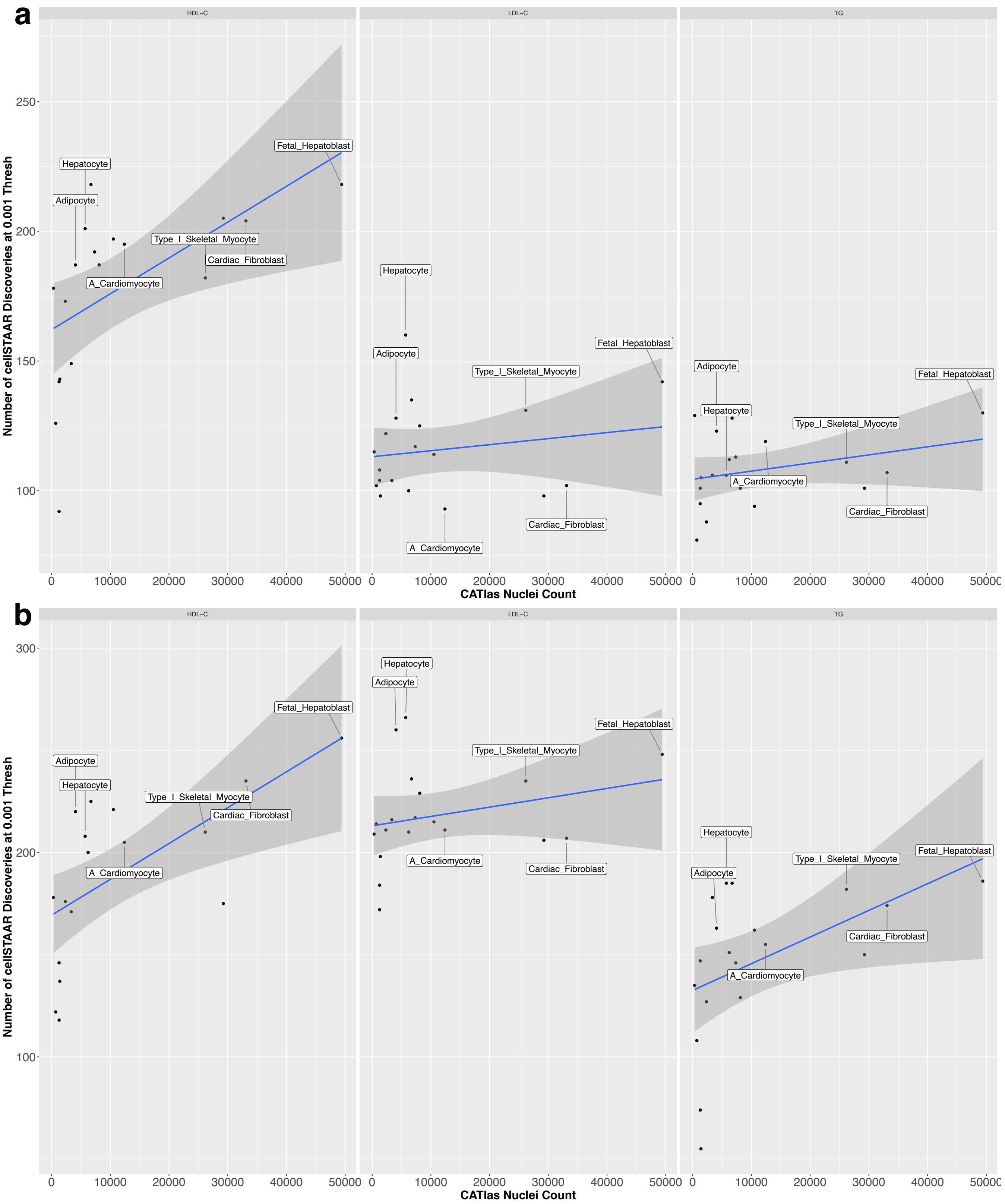
a
b
HDL–C
LDL–C
TG
Number of cellSTAAR Discoveries at 0.001 Thresh
CATlas Nuclei Count
Hepatocyte
Adipocyte
Fetal_Hepatoblast
Type_I_Skeletal_Myocyte
Cardiac_Fibroblast
A_Cardiomyocyte

**Supplementary Figure 3**: Manhattan and qq plots for cellSTAAR p-values from TOPMed for LDL-C, HDL-C, and TG, using the cell type with the most discoveries for each phenotype. Single-cell ATAC-seq data was used to select accessible genomic regions in each cell type by combining ATAC-seq peaks with regions in the top 20% of non-zero scores. Within the selected regions, rare variants within each of three ENCODE V3 element classes (dELS, pELS, PLS) were tested for association with the lipids phenotypes using cellSTAAR, which performs functionally-informed cell-type-level association testing using a generalized linear mixed model and multiple element-gene linking approaches, as described in the main text. **a/b,** Using Hepatocyte single-cell-sequencing-based data for LDL-C. **c/d**, Using Fetal_Hepatoblast single-cell-sequencing-based data for HDL-C. **e/f**, Using Beta single-cell-sequencing-based data for TG.

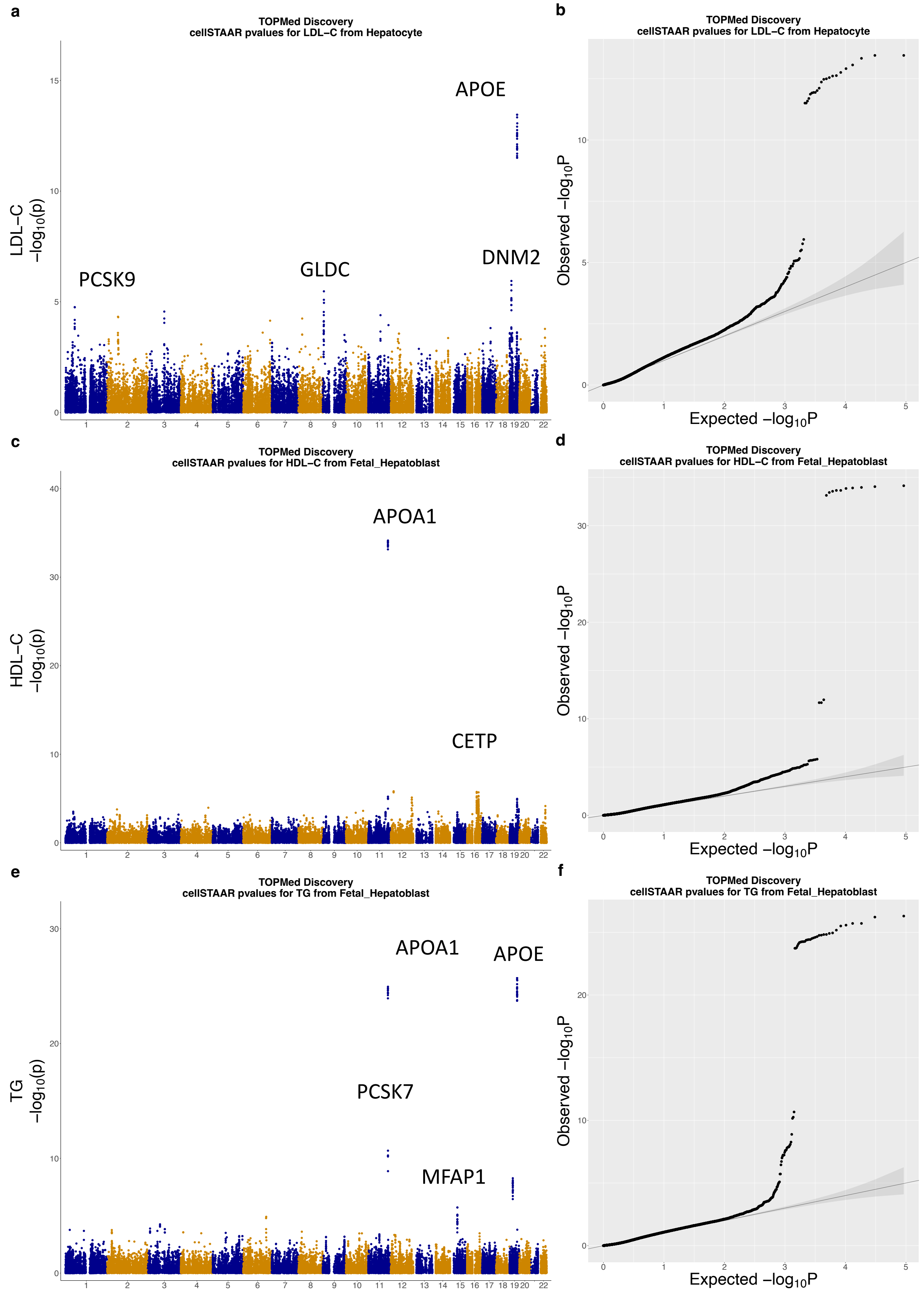
a
TOPMed Discovery
cellSTAAR pvalues for LDL−C from Hepatocyte
LDL−C
−log10(p)
APOE
PCSK9
GLDC
DNM2
b
TOPMed Discovery
cellSTAAR pvalues for LDL−C from Hepatocyte
Observed −log10P
Expected −log10P
c
TOPMed Discovery
cellSTAAR pvalues for HDL−C from Fetal_Hepatoblast
HDL−C
−log10(p)
APOA1
CETP
d
TOPMed Discovery
cellSTAAR pvalues for HDL−C from Fetal_Hepatoblast
Observed −log10P
Expected −log10P
e
TOPMed Discovery
cellSTAAR pvalues for TG from Fetal_Hepatoblast
TG
−log10(p)
APOA1
APOE
PCSK7
MFAP1
f
TOPMed Discovery
cellSTAAR pvalues for TG from Fetal_Hepatoblast
Observed −log10P
Expected −log10P

**Supplementary Figure 4**: Manhattan and qq plots for cellSTAAR p-values from the UKB replication phase for LDL-C, HDL-C, and TG, using the cell type with the most discoveries for each phenotype. Single-cell ATAC-seq data was used to select accessible genomic regions in each cell type by combining ATAC-seq peaks with regions in the top 20% of non-zero scores. Within the selected regions, rare variants within each of three ENCODE V3 element classes (dELS, pELS, PLS) were tested for association with the lipids phenotypes using cellSTAAR, which performs functionally-informed cell-type-level association testing using a generalized linear mixed model and multiple element-gene linking approaches, as described in the main text. **a,** Using Hepatocyte single-cell-sequencing-based data for LDL-C. **b**, Using Fetal_Hepatoblast single-cell-sequencing-based data for HDL-C. **c**, Using Hepatocyte single-cell-sequencing-based data for TG.

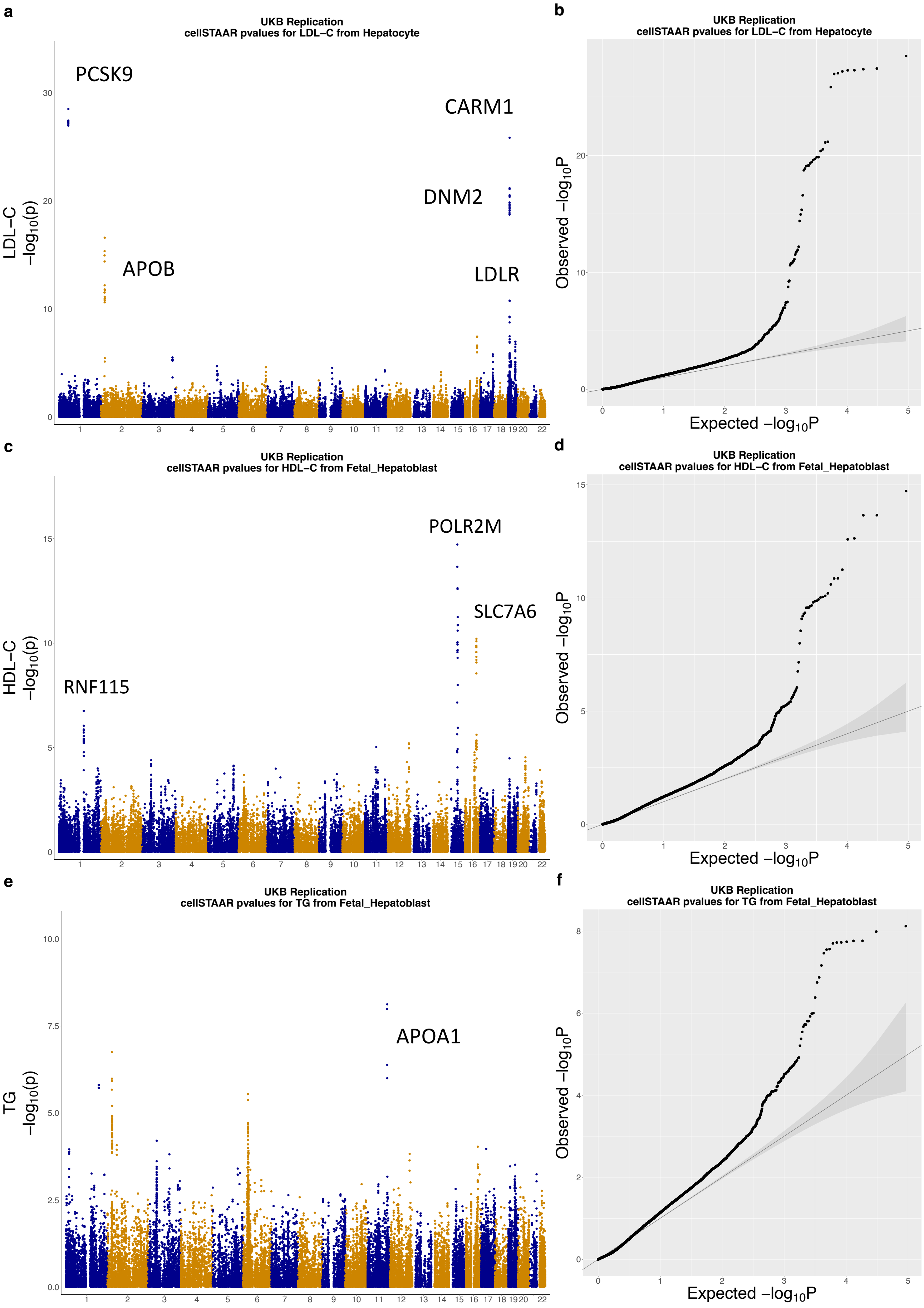
a
UKB Replication
cellSTAAR pvalues for LDL−C from Hepatocyte
LDL−C
$-\log_{10}(p)$
PCSK9
APOB
CARM1
DNM2
LDLR
b
UKB Replication
cellSTAAR pvalues for LDL−C from Hepatocyte
Observed $-\log_{10}P$
Expected $-\log_{10}P$
c
UKB Replication
cellSTAAR pvalues for HDL−C from Fetal_Hepatoblast
HDL−C
$-\log_{10}(p)$
RNF115
POLR2M
SLC7A6
d
UKB Replication
cellSTAAR pvalues for HDL−C from Fetal_Hepatoblast
Observed $-\log_{10}P$
Expected $-\log_{10}P$
e
UKB Replication
cellSTAAR pvalues for TG from Fetal_Hepatoblast
TG
$-\log_{10}(p)$
APOA1
f
UKB Replication
cellSTAAR pvalues for TG from Fetal_Hepatoblast
Observed $-\log_{10}P$
Expected $-\log_{10}P$

**Supplementary Figure 5:** Manhattan and qq plots for cellSTAAR p-values from the TOPMed Discovery and UKB replication phase for the binary highLDL phenotype, using the cell type with the most discoveries (Fetal_Endothelial_Hepatic). Single-cell ATAC-seq data was used to select accessible genomic regions in each cell type by combining  ATAC-seq peaks with regions in the top 20% of non-zero scores. Within the selected regions, rare variants within each of three ENCODE V3 element classes (dELS, pELS, PLS) were tested for association with the lipids phenotypes using cellSTAAR, which performs functionally-informed cell-type-level association testing using a generalized linear mixed model and multiple element-gene linking approaches, as described in the main text. **a/b,** TOPMed discovery cohort, **c/d** UKB replication cohort.

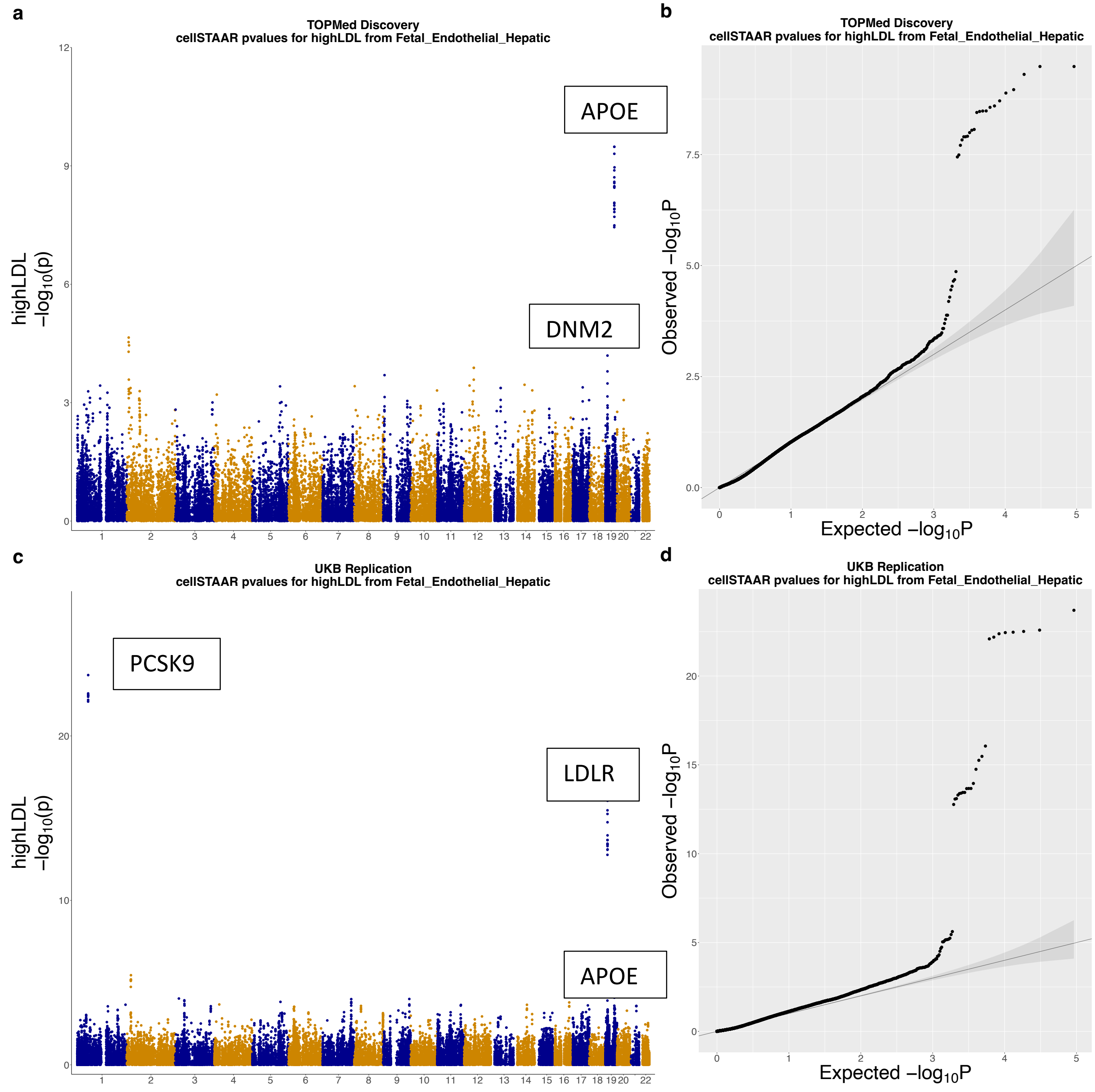

**Supplementary Figure 6:** Variability in association across linking approaches and cell types for *PCSK9, LDLR, LPA, and APOB* in the TOPMed discovery phase for LDL-C. Single-cell ATAC-seq data was used to select accessible genomic regions in each cell type by combining ATAC-seq peaks with regions in the top 20% of non-zero scores. Within the selected regions, rare variants within each of three ENCODE V3 element classes (dELS, pELS, PLS) were tested for association with the lipids phenotypes using cellSTAAR, which performs functionally-informed cell-type-level association testing using a generalized linear mixed model and multiple element-gene linking approaches, as described in the main text. **a/c/e/g,** Association results for dELS variants from distance-based, EpiMap, Activity-by-Contact (ABC), SCREEN eQTL, and SCREEN 3D linking approaches and cellSTAAR omnibus p-value across 8 representative cell types for *PCSK9, LDLR, LPA,* and *APOB*. **b/d/f/h,** cellSTAAR p-value for dELS variants across all 19 cell types analyzed for *PCSK9, LDLR, LPA,* and *APOB.*

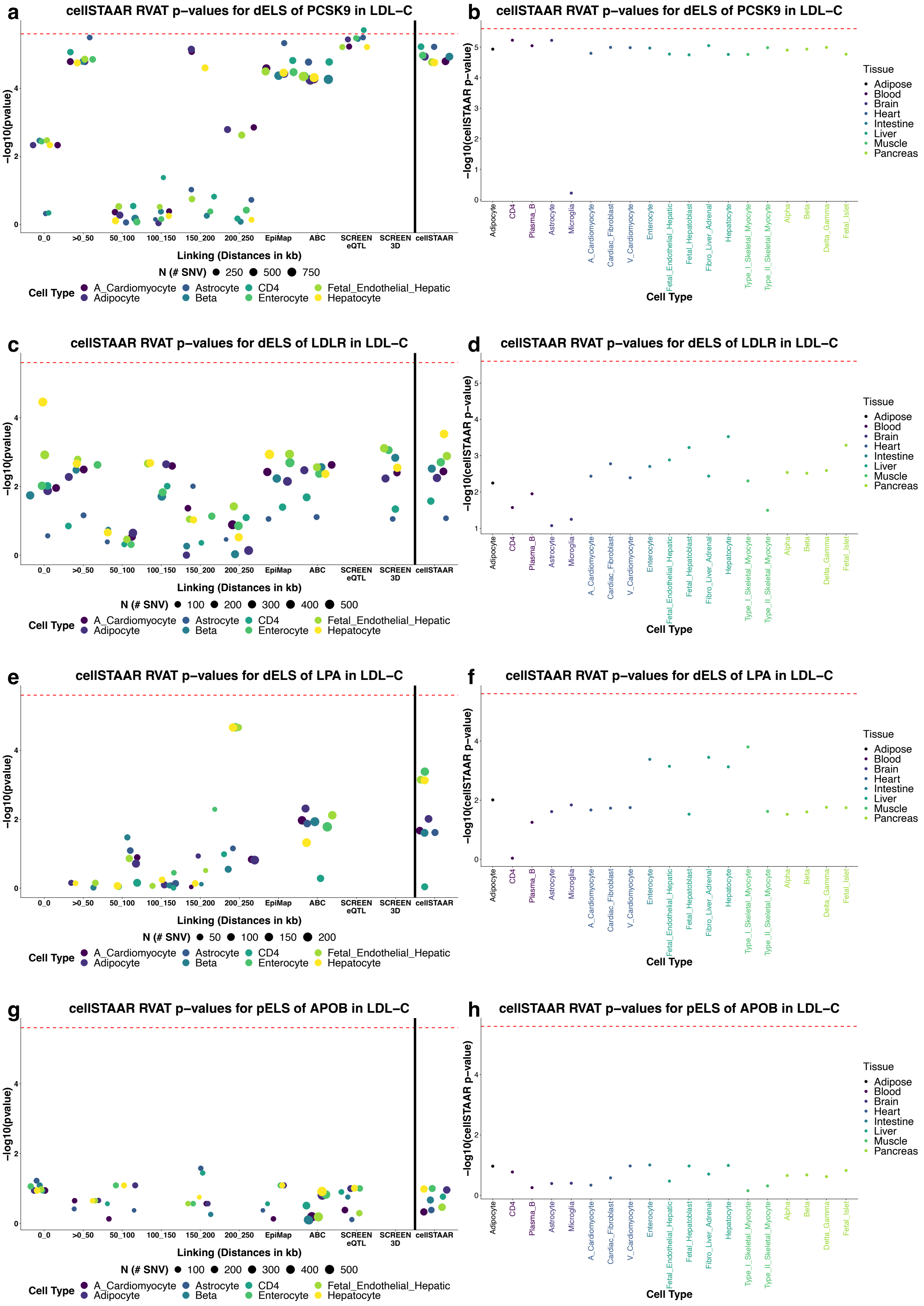

a
cellSTAAR RVAT p−values for dELS of PCSK9 in LDL−C
−log10(pvalue)
0_0
>0_50
50_100
100_150
150_200
200_250
EpiMap
ABC
SCREEN eQTL
SCREEN 3D
cellSTAAR
Linking (Distances in kb)
N (# SNV) 250 500 750
Cell Type A_Cardiomyocyte Adipocyte Astrocyte Beta CD4 Enterocyte Fetal_Endothelial_Hepatic Hepatocyte
b
cellSTAAR RVAT p−values for dELS of PCSK9 in LDL−C
−log10(cellSTAAR p−value)
Adipocyte
CD4
Plasma_B
Astrocyte
Microglia
A_Cardiomyocyte
Cardiac_Fibroblast
V_Cardiomyocyte
Enterocyte
Fetal_Endothelial_Hepatic
Fetal_Hepatoblast
Fibro_Liver_Adrenal
Hepatocyte
Type_I_Skeletal_Myocyte
Type_II_Skeletal_Myocyte
Alpha
Beta
Delta_Gamma
Fetal_Islet
Cell Type
Tissue
Adipose
Blood
Brain
Heart
Intestine
Liver
Muscle
Pancreas
c
cellSTAAR RVAT p−values for dELS of LDLR in LDL−C
N (# SNV) 100 200 300 400 500
d
cellSTAAR RVAT p−values for dELS of LDLR in LDL−C
e
cellSTAAR RVAT p−values for dELS of LPA in LDL−C
N (# SNV) 50 100 150 200
f
cellSTAAR RVAT p−values for dELS of LPA in LDL−C
g
cellSTAAR RVAT p−values for pELS of APOB in LDL−C
N (# SNV) 100 200 300 400 500
h
cellSTAAR RVAT p−values for pELS of APOB in LDL−C

**Supplementary Figure 7:** Variability in association across linking approaches and cell types for *PCSK9, LDLR, LPA, and APOB* in the UKB replication phase for LDL-C. Single-cell ATAC-seq data was used to select accessible genomic regions in each cell type by combining ATAC-seq peaks with regions in the top 20% of non-zero scores. Within the selected regions, rare variants within each of three ENCODE V3 element classes (dELS, pELS, PLS) were tested for association with the lipids phenotypes using cellSTAAR, which performs functionally-informed cell-type-level association testing using a generalized linear mixed model and multiple element-gene linking approaches, as described in the main text. **a/c/e/g,** Association results for dELS variants from distance-based, EpiMap, Activity-by-Contact (ABC), SCREEN eQTL, and SCREEN 3D linking approaches and cellSTAAR omnibus p-value across 8 representative cell types for *PCSK9, LDLR, LPA*, and *APOB*. **b/d/f/h,** cellSTAAR p-value for dELS variants across all 19 cell types analyzed for *PCSK9, LDLR, LPA*, and *APOB.*

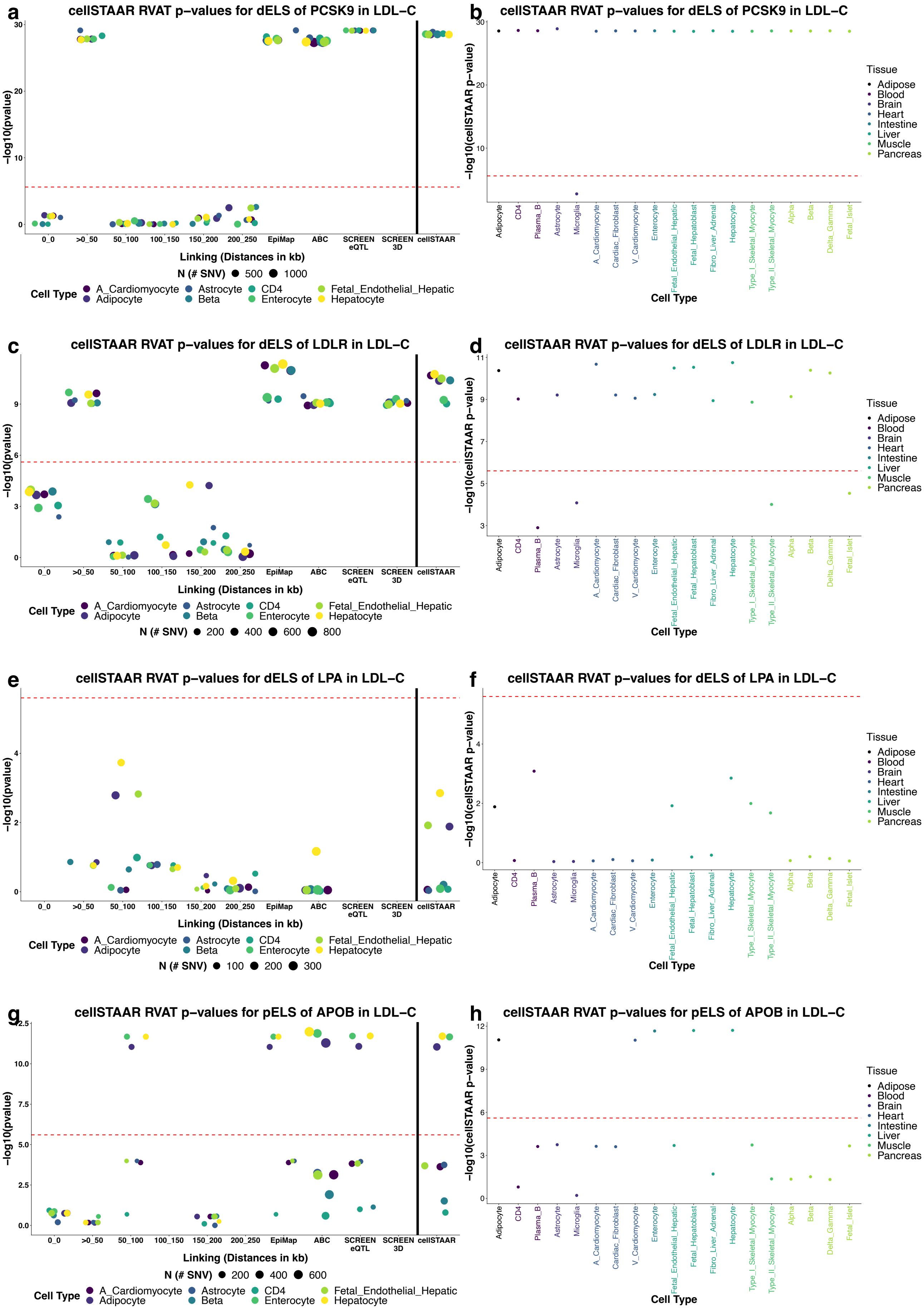
a
cellSTAAR RVAT p−values for dELS of PCSK9 in LDL−C
−log10(pvalue)
0_0
>0_50
50_100
100_150
150_200
200_250
EpiMap
ABC
SCREEN eQTL
SCREEN 3D
cellSTAAR
Linking (Distances in kb)
N (# SNV) 500 1000
Cell Type A_Cardiomyocyte Adipocyte Astrocyte Beta CD4 Enterocyte Fetal_Endothelial_Hepatic Hepatocyte
b
cellSTAAR RVAT p−values for dELS of PCSK9 in LDL−C
−log10(cellSTAAR p−value)
Adipocyte
CD4
Plasma_B
Astrocyte
Microglia
A_Cardiomyocyte
Cardiac_Fibroblast
V_Cardiomyocyte
Enterocyte
Fetal_Endothelial_Hepatic
Fetal_Hepatoblast
Fibro_Liver_Adrenal
Hepatocyte
Type_I_Skeletal_Myocyte
Type_II_Skeletal_Myocyte
Alpha
Beta
Delta_Gamma
Fetal_Islet
Cell Type
Tissue Adipose Blood Brain Heart Intestine Liver Muscle Pancreas
c
cellSTAAR RVAT p−values for dELS of LDLR in LDL−C
N (# SNV) 200 400 600 800
d
cellSTAAR RVAT p−values for dELS of LDLR in LDL−C
e
cellSTAAR RVAT p−values for dELS of LPA in LDL−C
N (# SNV) 100 200 300
f
cellSTAAR RVAT p−values for dELS of LPA in LDL−C
g
cellSTAAR RVAT p−values for pELS of APOB in LDL−C
N (# SNV) 200 400 600
h
cellSTAAR RVAT p−values for pELS of APOB in LDL−C

**Supplementary Figure 8**: Variability in association across linking approaches and cell types for the *APOE* locus in the UKB replication phase for LDL-C. Single-cell ATAC-seq data was used to select accessible genomic regions in each cell type by combining ATAC-seq peaks with regions in the top 20% of non-zero scores. Within the selected regions, rare variants within each of three ENCODE V3 element classes (dELS, pELS, PLS) were tested for association with the lipids phenotypes using cellSTAAR, which performs functionally-informed cell-type-level association testing using a generalized linear mixed model and multiple element-gene linking approaches, as described in the main text.

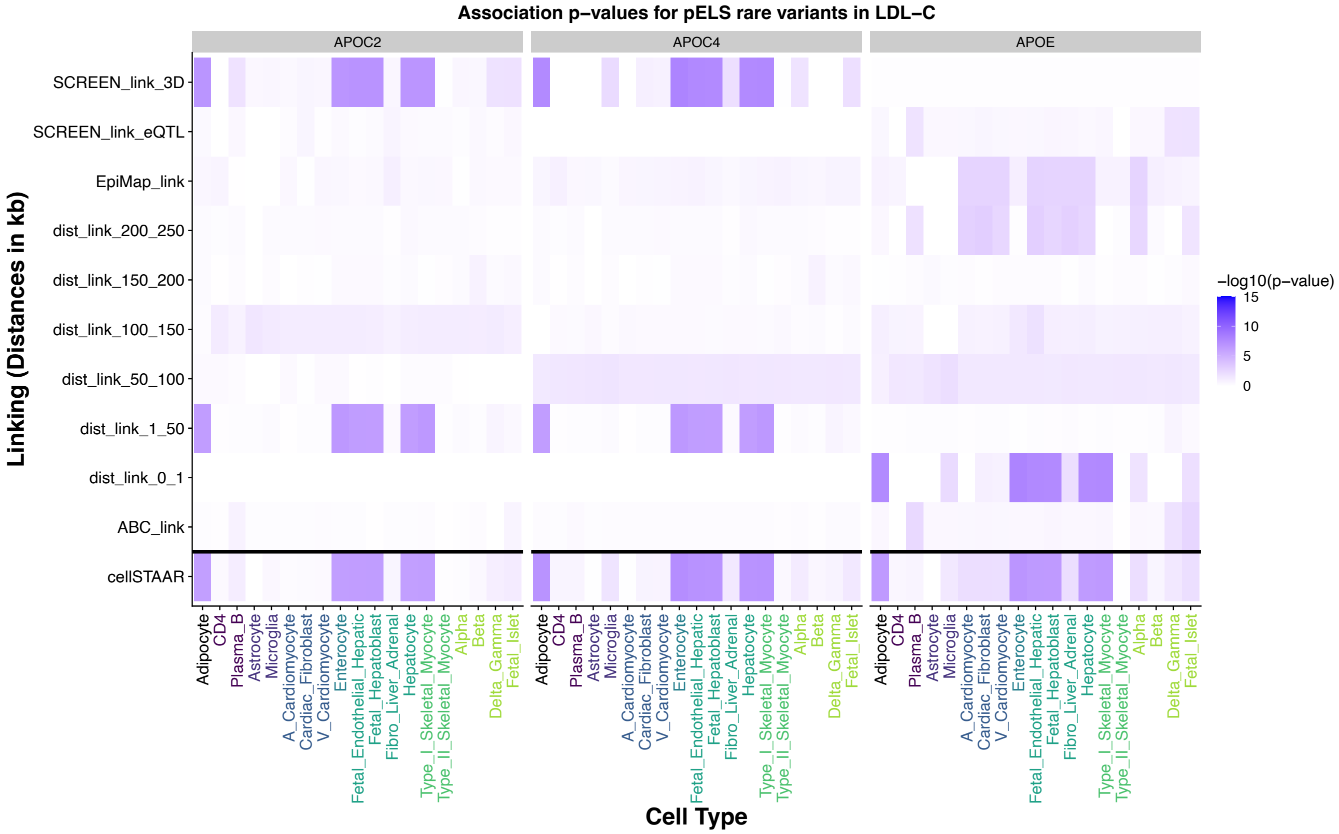

**Supplementary Figure 9:** Variability in association across linking approaches and cell types for the *CETP* locus in the TOPMed discovery and UKB replication phases in HDL-C. Single-cell ATAC-seq data was used to select accessible genomic regions in each cell type by combining ATAC-seq peaks with regions in the top 20% of non-zero scores. Within the selected regions, rare variants within each of three ENCODE V3 element classes (dELS, pELS, PLS) were tested for association with the lipids phenotypes using cellSTAAR, which performs functionally-informed cell-type-level association testing using a generalized linear mixed model and multiple element-gene linking approaches, as described in the main text. **a,** TOPMed association results for PLS variants from distance-based, SCREEN eQTL, and SCREEN 3D linking approaches and cellSTAAR omnibus p-value across 8 representative cell types for *CETP*. **b,** TOPMed cellSTAAR p-value for PLS variants linked to *CETP* across all 19 cell types analyzed. **c,** UKB association results for PLS variants from distance-based, SCREEN eQTL, and SCREEN 3D linking approaches and cellSTAAR omnibus p-value across 8 representative cell types for *CETP*. **d,** UKB cellSTAAR p-value for PLS variants linked to *CETP* across all 19 cell types analyzed.

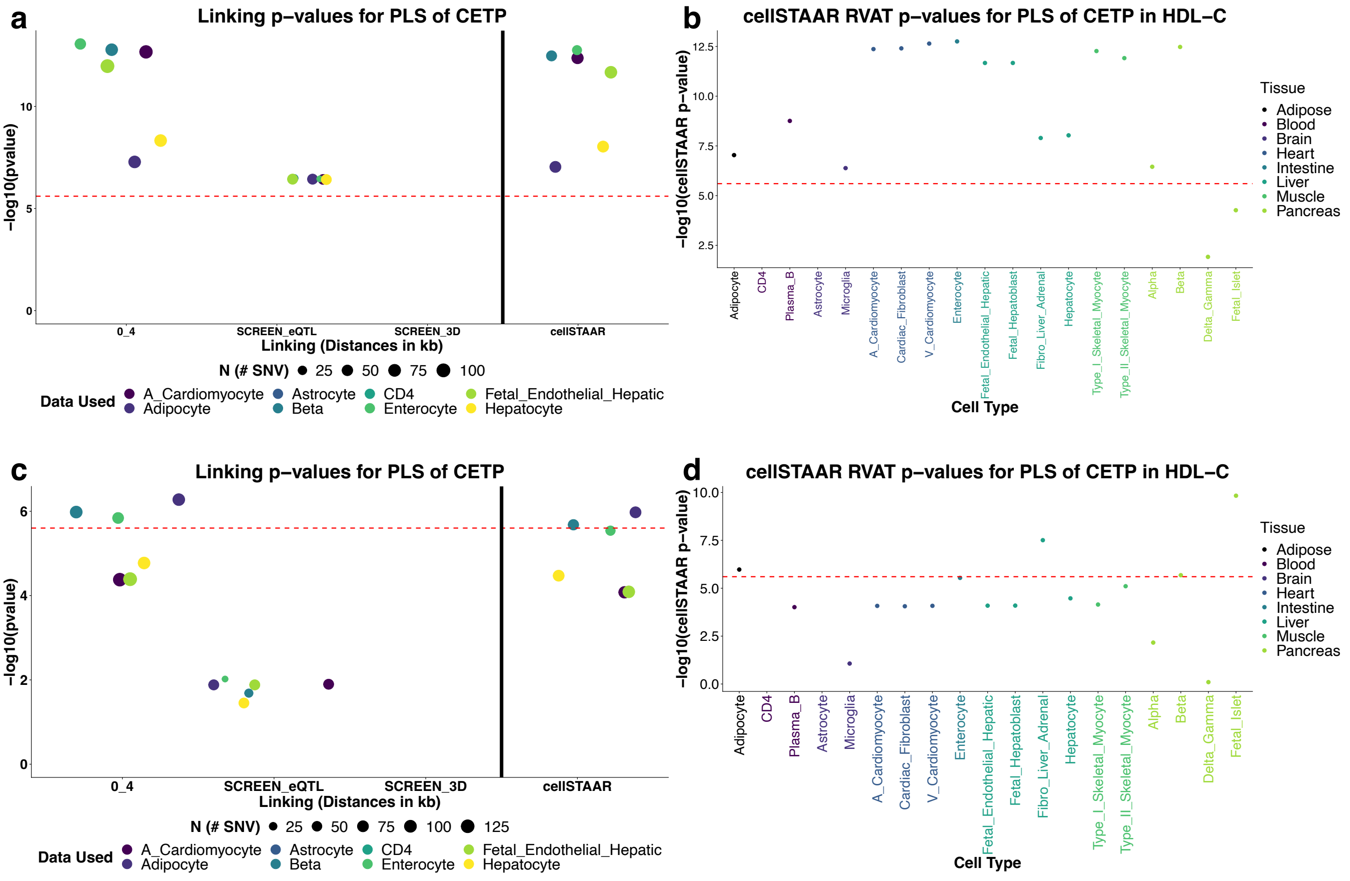

**Supplementary Figure 10**: Discoveries (cellSTAAR p-value < $1x10^{-3}$) by cell type and phenotype for TOPMed discovery and UK Biobank replication phases. Single-cell ATAC-seq data was used to select accessible genomic regions in each cell type by combining ATAC-seq peaks with regions in the top 20% of non-zero scores. Within the selected regions, rare variants within each of three ENCODE V3 element classes (dELS, pELS, PLS) were tested for association with the lipids phenotypes using cellSTAAR, which performs functionally-informed cell-type-level association testing using a generalized linear mixed model and multiple element-gene linking approaches, as described in the main text. Within a phenotype and cell type, the height of the bars corresponds to the number of discoveries, which is also printed after the cell type name. Bars "STAAR_DHS" and "STAAR_CAGE" refer to DHS and CAGE defined enhancers and promoters as tested for association using STAARPipeline [2] (no single-cell data used), and "all_ct_ACAT" refers to the p-value that results from combining cellSTAAR p-values from all 19 cell types using the ACAT method. Bars "bulk_liver_1", "bulk_liver_2", and "bulk_liver_3" refer to analyses run using ENCODE bulk liver ATAC-seq data (see section "Use of ENCODE bulk liver ATAC-seq data**"** in the main text) to compare the use of single-cell ATAC-seq data to bulk ATAC-seq data.

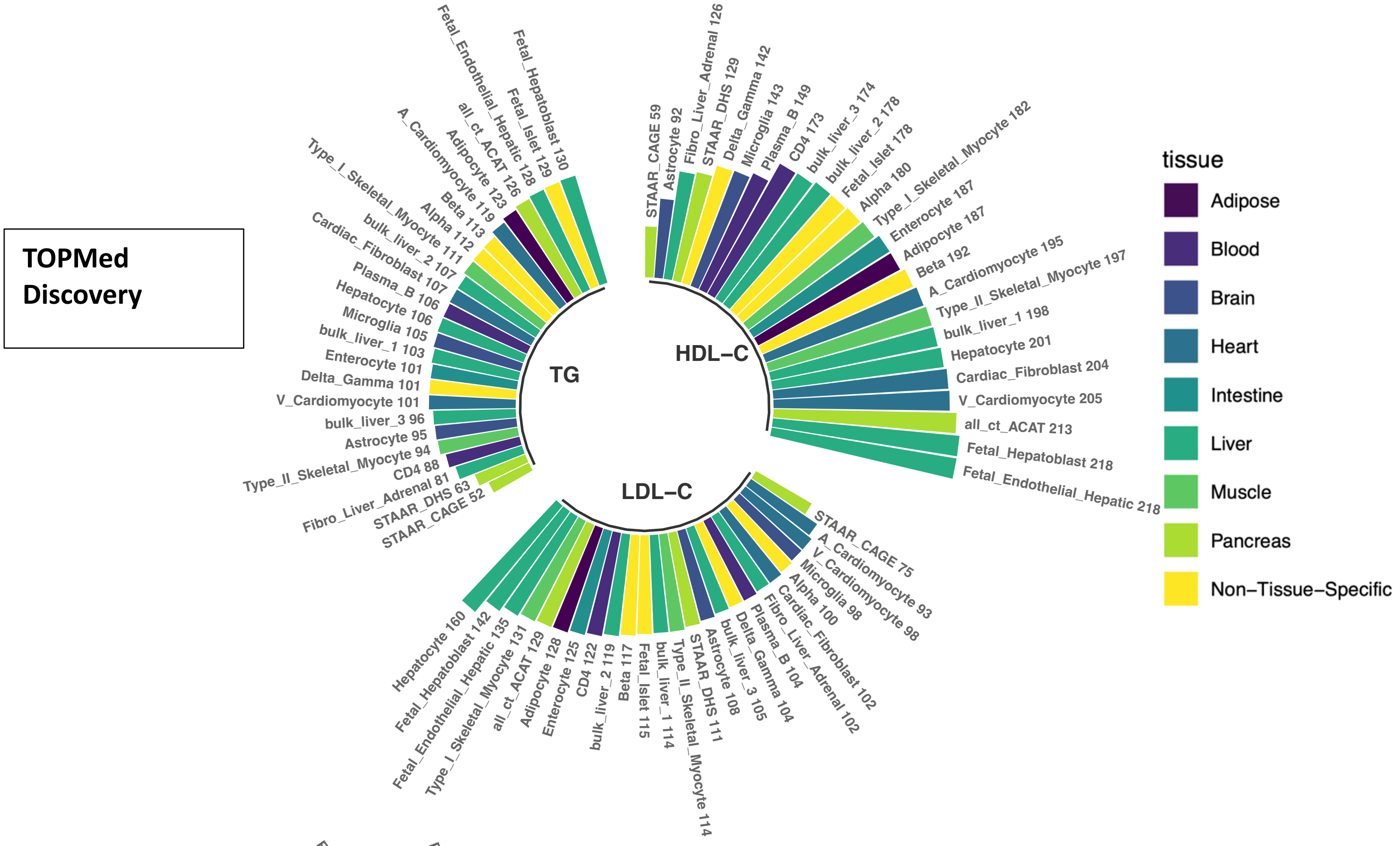
TOPMed Discovery
TG
HDL−C
LDL−C
Fetal_Hepatoblast 130
Fetal_Islet 129
Fetal_Endothelial_Hepatic 128
all_ct_ACAT 126
Adipocyte 123
A_Cardiomyocyte 119
Beta 113
Alpha 112
Type_I_Skeletal_Myocyte 111
bulk_liver_2 107
Cardiac_Fibroblast 107
Plasma_B 106
Hepatocyte 106
Microglia 105
bulk_liver_1 103
Enterocyte 101
Delta_Gamma 101
V_Cardiomyocyte 101
bulk_liver_3 96
Astrocyte 95
Type_II_Skeletal_Myocyte 94
CD4 88
Fibro_Liver_Adrenal 81
STAAR_DHS 63
STAAR_CAGE 52
STAAR_CAGE 59
Astrocyte 92
Fibro_Liver_Adrenal 126
STAAR_DHS 129
Delta_Gamma 142
Microglia 143
Plasma_B 149
CD4 173
bulk_liver_3 174
bulk_liver_2 178
Fetal_Islet 178
Alpha 180
Type_I_Skeletal_Myocyte 182
Enterocyte 187
Adipocyte 187
Beta 192
A_Cardiomyocyte 195
Type_II_Skeletal_Myocyte 197
bulk_liver_1 198
Hepatocyte 201
Cardiac_Fibroblast 204
V_Cardiomyocyte 205
all_ct_ACAT 213
Fetal_Hepatoblast 218
Fetal_Endothelial_Hepatic 218
STAAR_CAGE 75
A_Cardiomyocyte 93
V_Cardiomyocyte 98
Microglia 98
Alpha 100
Cardiac_Fibroblast 102
Fibro_Liver_Adrenal 102
Plasma_B 104
Delta_Gamma 104
bulk_liver_3 105
Astrocyte 108
STAAR_DHS 111
Type_II_Skeletal_Myocyte 114
bulk_liver_1 114
Fetal_Islet 115
Beta 117
bulk_liver_2 119
CD4 122
Enterocyte 125
Adipocyte 128
all_ct_ACAT 129
Type_I_Skeletal_Myocyte 131
Fetal_Endothelial_Hepatic 135
Fetal_Hepatoblast 142
Hepatocyte 160
tissue
Adipose
Blood
Brain
Heart
Intestine
Liver
Muscle
Pancreas
Non−Tissue−Specific

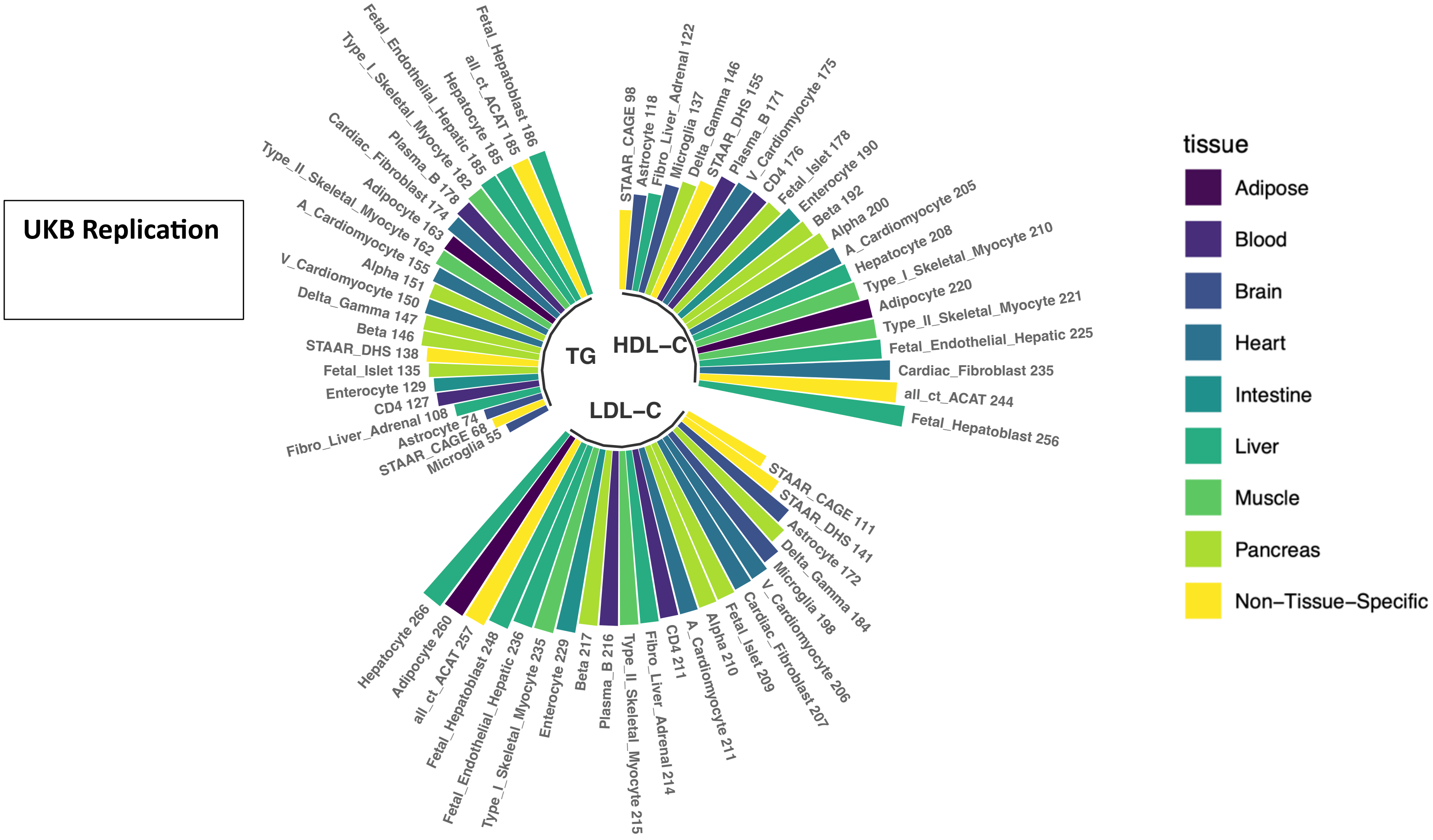
UKB Replication
TG
HDL−C
LDL−C
Fetal_Hepatoblast 186
all_ct_ACAT 185
Hepatocyte 185
Fetal_Endothelial_Hepatic 185
Type_I_Skeletal_Myocyte 182
Plasma_B 178
Cardiac_Fibroblast 174
Adipocyte 163
Type_II_Skeletal_Myocyte 162
A_Cardiomyocyte 155
Alpha 151
V_Cardiomyocyte 150
Delta_Gamma 147
Beta 146
STAAR_DHS 138
Fetal_Islet 135
Enterocyte 129
CD4 127
Fibro_Liver_Adrenal 108
Astrocyte 74
STAAR_CAGE 68
Microglia 55
STAAR_CAGE 98
Astrocyte 118
Fibro_Liver_Adrenal 122
Microglia 137
Delta_Gamma 146
STAAR_DHS 155
Plasma_B 171
V_Cardiomyocyte 175
CD4 176
Fetal_Islet 178
Enterocyte 190
Beta 192
Alpha 200
A_Cardiomyocyte 205
Hepatocyte 208
Type_I_Skeletal_Myocyte 210
Adipocyte 220
Type_II_Skeletal_Myocyte 221
Fetal_Endothelial_Hepatic 225
Cardiac_Fibroblast 235
all_ct_ACAT 244
Fetal_Hepatoblast 256
STAAR_CAGE 111
STAAR_DHS 141
Astrocyte 172
Delta_Gamma 184
Microglia 198
V_Cardiomyocyte 206
Cardiac_Fibroblast 207
Fetal_Islet 209
Alpha 210
A_Cardiomyocyte 211
CD4 211
Fibro_Liver_Adrenal 214
Type_II_Skeletal_Myocyte 215
Plasma_B 216
Beta 217
Enterocyte 229
Type_I_Skeletal_Myocyte 235
Fetal_Endothelial_Hepatic 236
Fetal_Hepatoblast 248
all_ct_ACAT 257
Adipocyte 260
Hepatocyte 266
tissue
Adipose
Blood
Brain
Heart
Intestine
Liver
Muscle
Pancreas
Non−Tissue−Specific

**Supplementary Figure 11**: Enrichment of common variant (MAF > 1%) discoveries (5E-8 threshold) in TOPMed discovery phase by cell type. Single-cell ATAC-seq data was used to select accessible genomic regions in each cell type by combining ATAC-seq peaks with regions in the top 20% of non-zero scores. Within the selected regions, all common variants belonging to one of the three ENCODE V3 element classes (dELS, pELS, PLS) were individually tested for association with the lipids phenotypes using a standard generalized linear mixed modelling framework. Variant-level p-values were produced with the "Individual_Analysis" function in the STAARPipeline R package. The x-axis is ordered by the sum of the three phenotypes, with cell types further to the left having more overall enrichment as compared to those on the right.

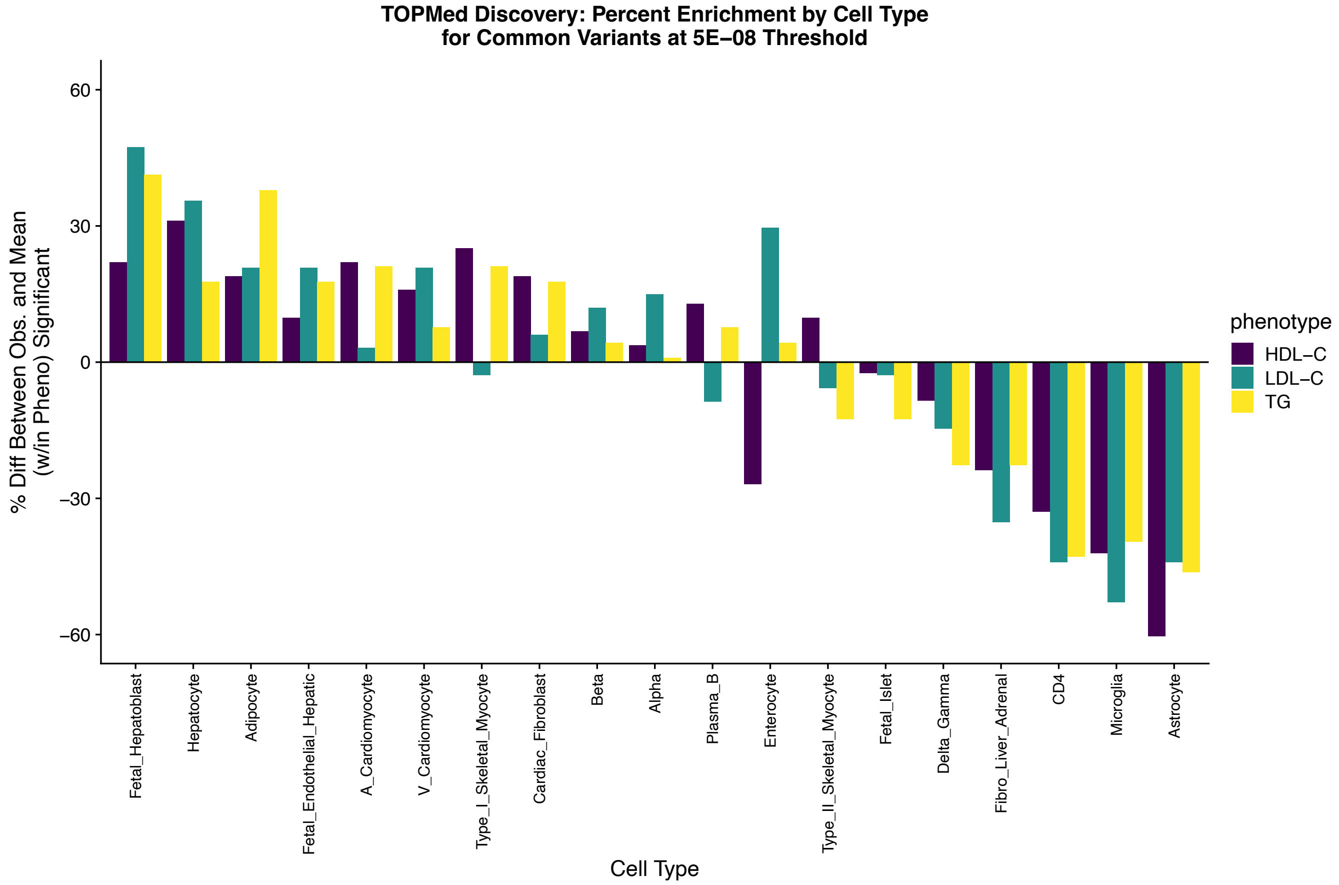

**Supplementary Figure 12**: Enrichment of common variant (MAF > 1%) discoveries (5E-8 threshold) in TOPMed discovery phase by phenotype. Single-cell ATAC-seq data was used to select accessible genomic regions in each cell type by combining ATAC-seq peaks with regions in the top 20% of non-zero scores. Within the selected regions, common variants belonging to one of the three ENCODE V3 element classes (dELS, pELS, PLS) were individually tested for association with the lipids phenotypes using a standard generalized linear mixed modelling framework. Variant-level p-values were produced with the "Individual_Analysis" function in the STAARPipeline R package. Within a phenotype, the height of the bars corresponds to the number of discoveries, which is also printed next to the cell type name.

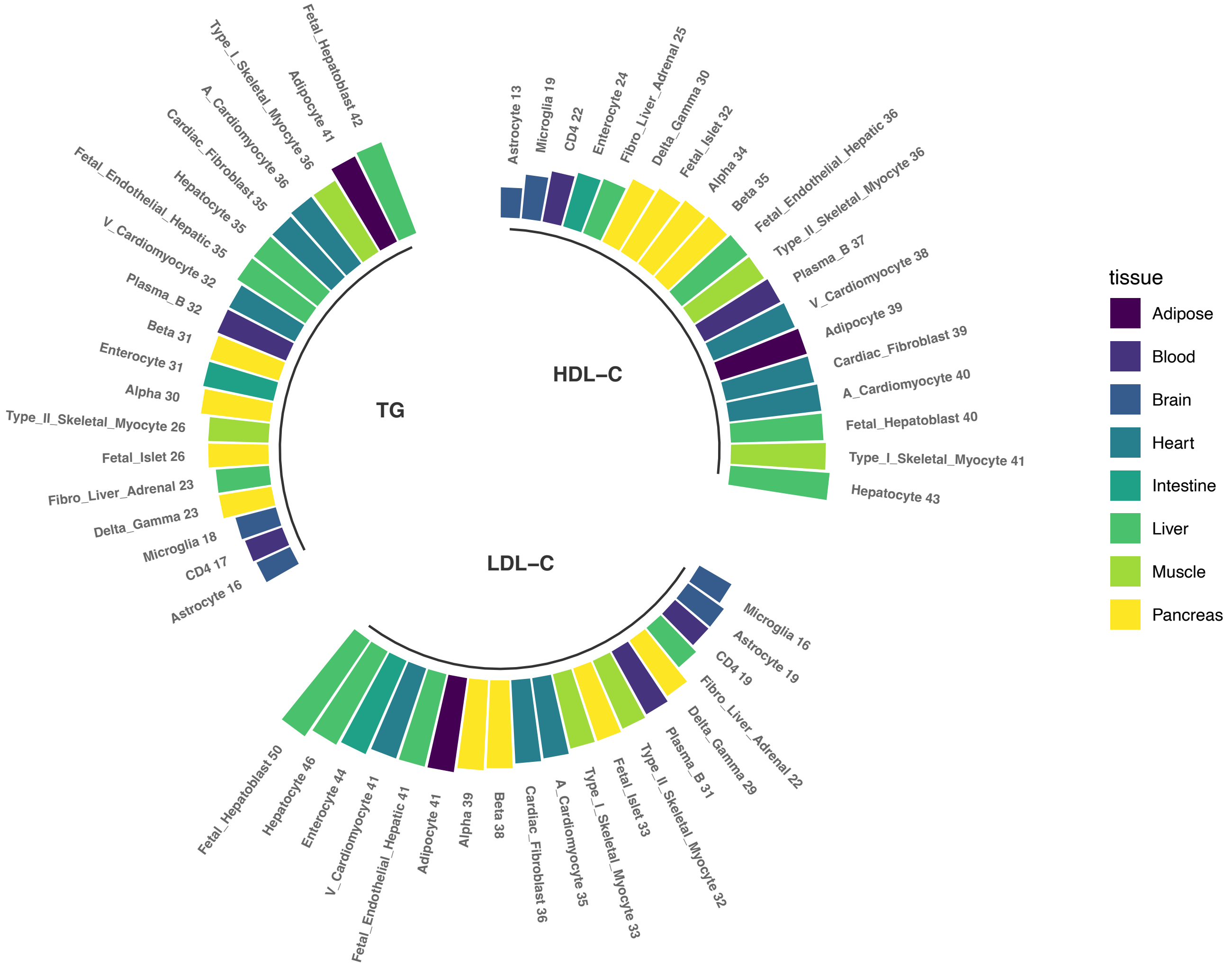

**Supplementary Figure 13**: Intuitive overview of the use of the IHW method to weight cellSTAAR gene-centric omnibus association p-values. Using a "side-information" covariate which is hypothesized to be informative about the association null hypothesis, such as a measure of absolute or differential expression, p-values $p_{cellSTAAR}$ are weighted to create a new p-values $p*_{cellSTAAR}$ which incorporates the side-information.

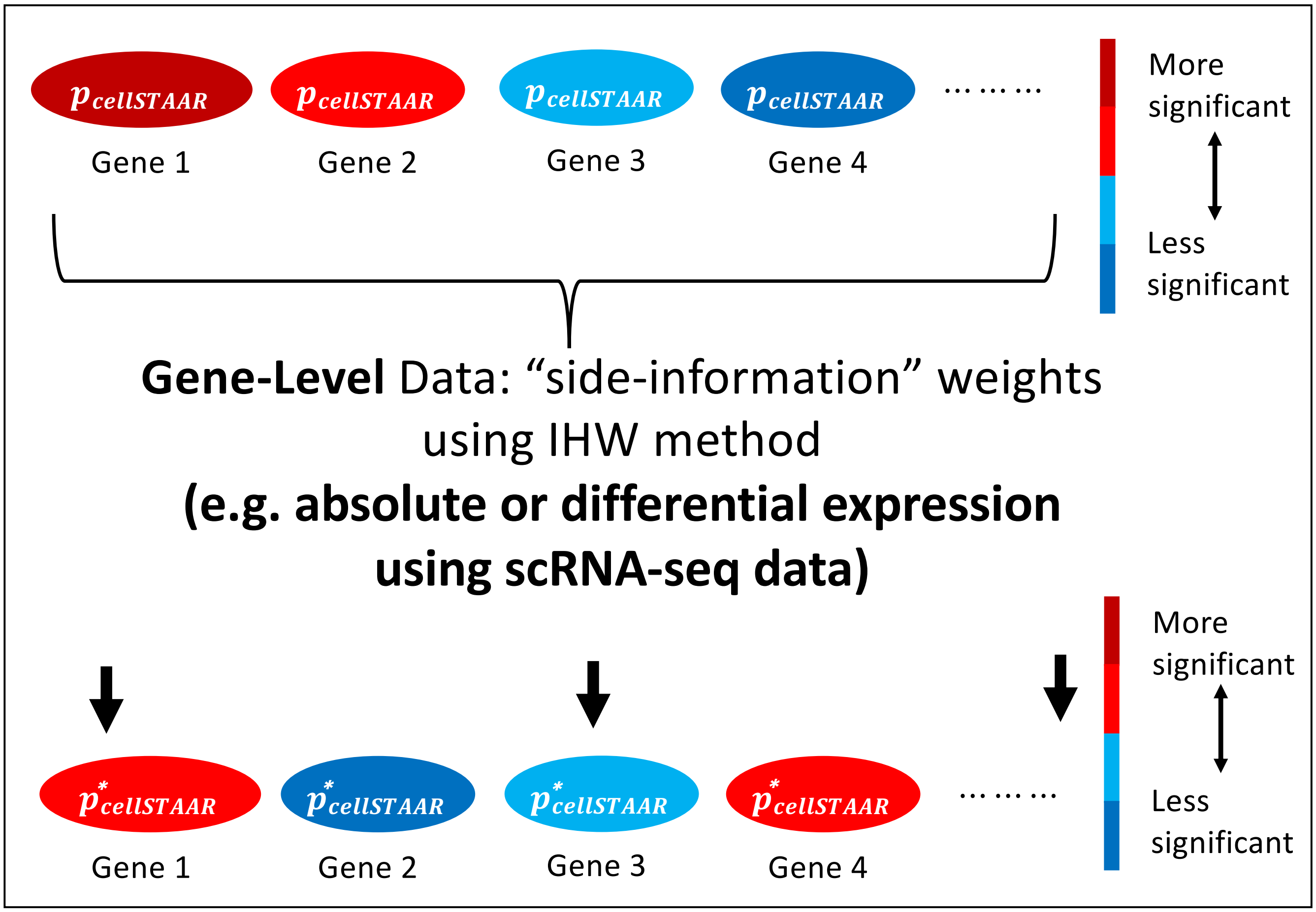

**Supplementary Figure 14**: Discoveries (cellSTAAR p-value < $1x10^{-3}$) by cell type (a) using cellSTAAR with no scRNA-seq data, (b) using cellSTAAR p-values adjusted using logFC values as "side-information" covariates as input to the IHW procedure, (c) using cellSTAAR with no scRNA-seq data, (d) using cellSTAAR p-values adjusted using the percentage of cells within a given cell type as "side-information" covariates as input to the IHW method. The number of discoveries is slightly reduced when using the IHW method with either relative or differential expression as side-information covariates.

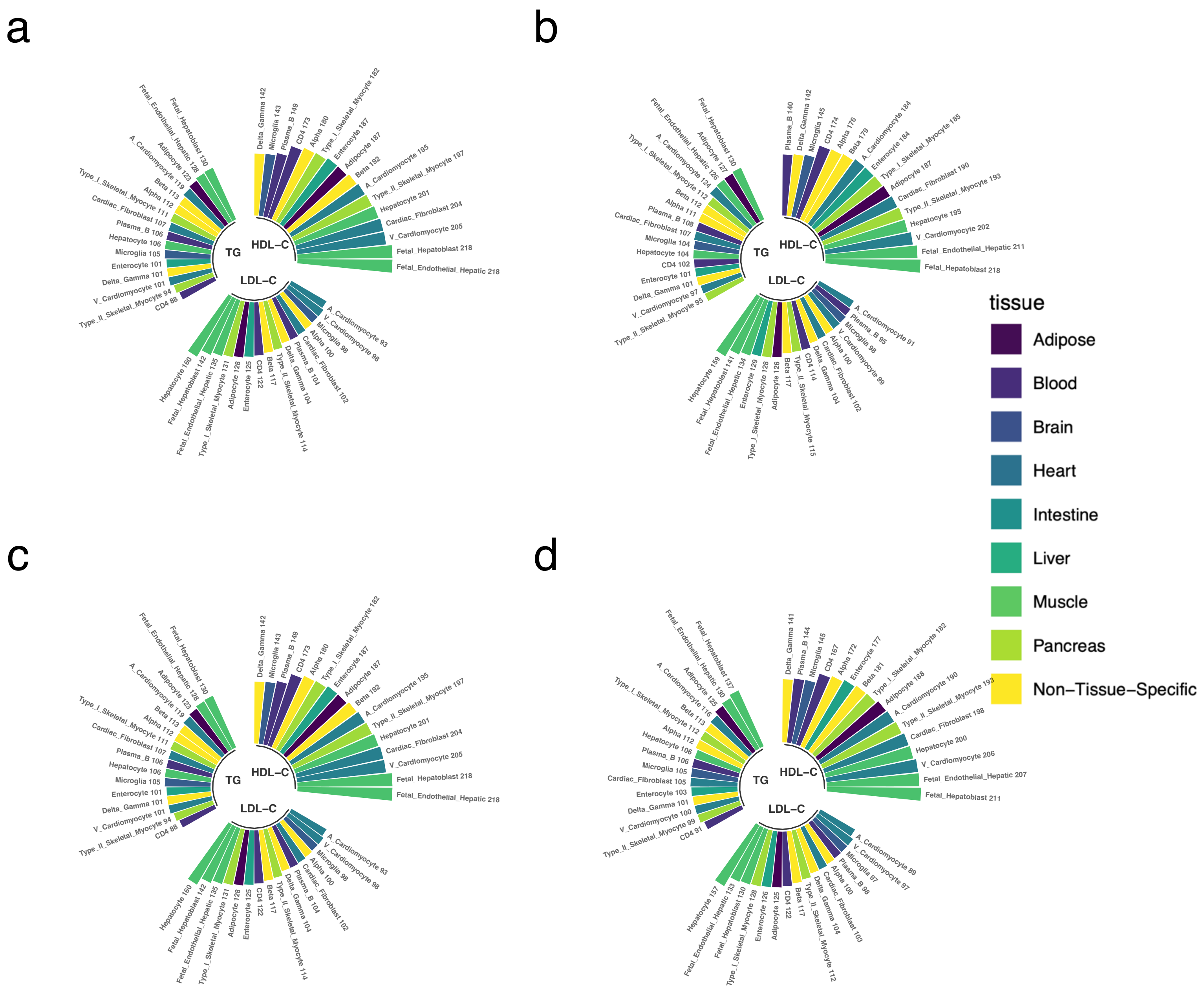

**Supplementary Figure 15**: Relationship between absolute expression (percentage of cells expressing each gene) and the -log10(cellSTAAR association p-value) using (a) Hepatocyte single-cell data to weight LDL-C association p-values, (b) CD4 single-cell data to weight LDL-C association p-values, (c) Hepatocyte single-cell data to weight HDL-C association p-values, (d) CD4 single-cell data to weight HDL-C association p-values, (e) Hepatocyte single-cell data to weight TG association p-values, (f) CD4 single-cell data to weight TG association p-values.

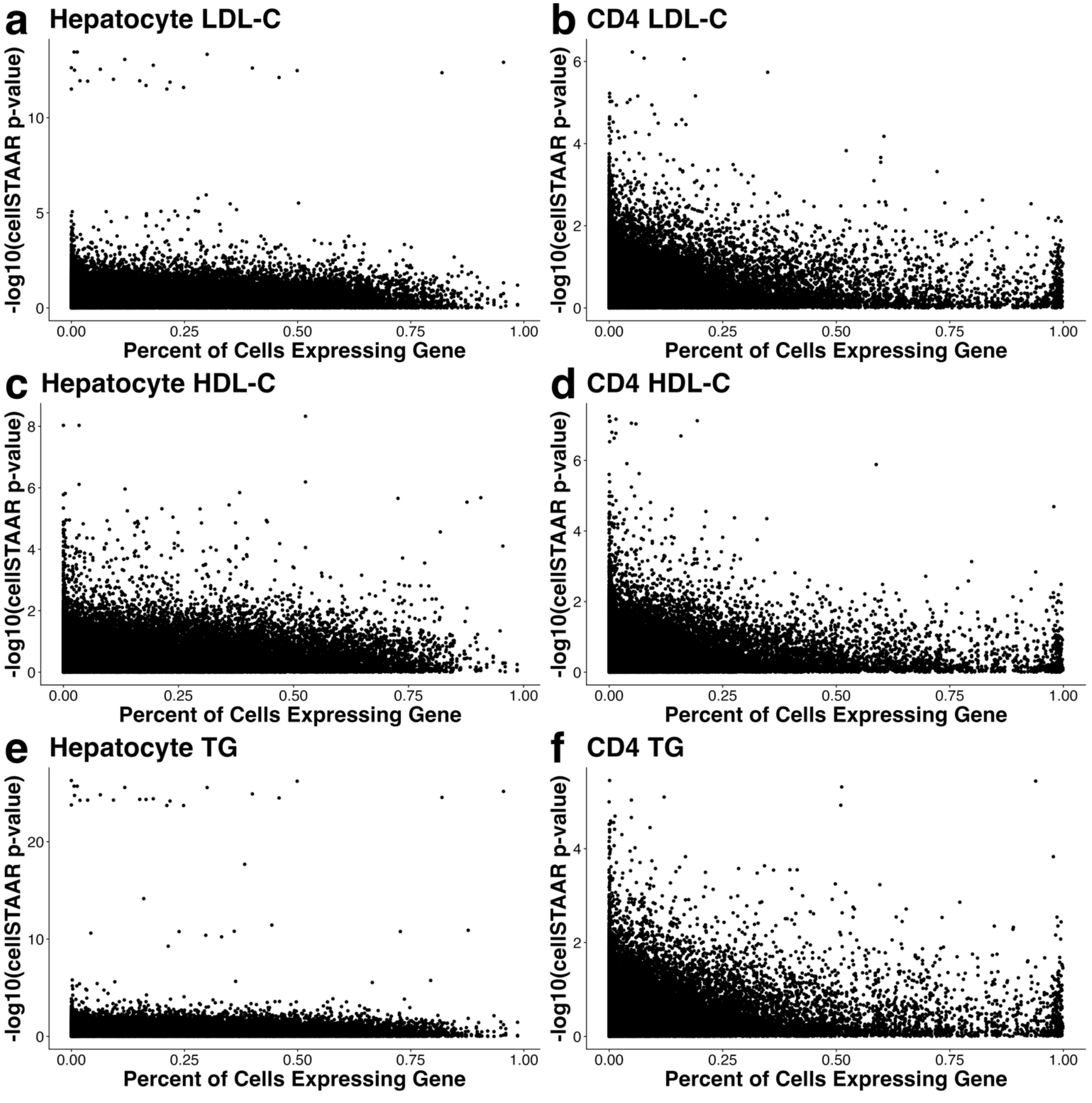

**Supplementary Figure 16**: Relationship between differential expression (log fold-change (logFC) comparing a cell type to other cell types) and the -log10(cellSTAAR association p-value) using (a) Hepatocyte single-cell data to weight LDL-C association p-values, (b) CD4 single-cell data to weight LDL-C association p-values, (c) Hepatocyte single-cell data to weight HDL-C association p-values, (d) CD4 single-cell data to weight HDL-C association p-values, (e) Hepatocyte single-cell data to weight TG association p-values, (f) CD4 single-cell data to weight TG association p-values.

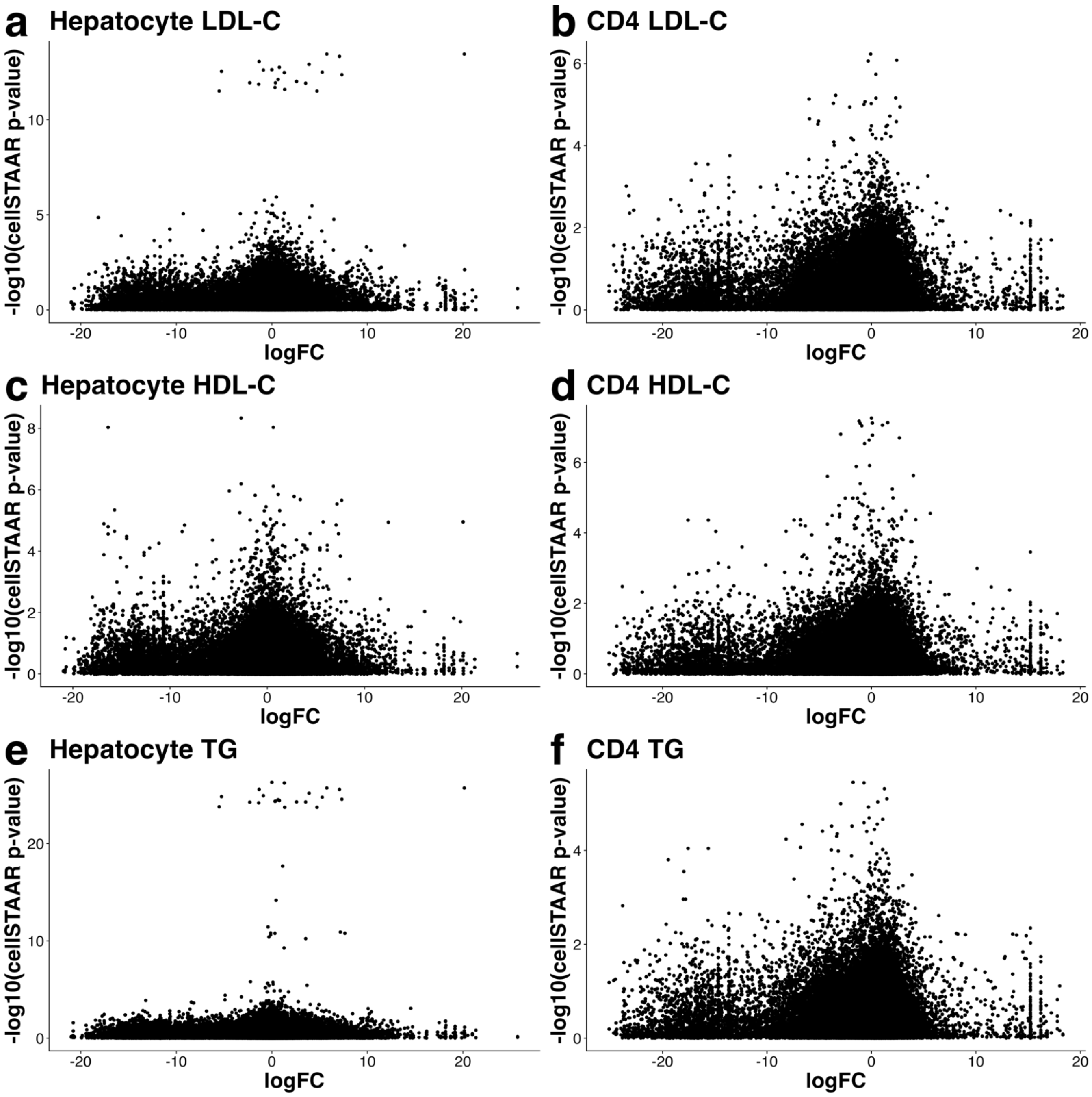

**Supplementary Figure 17:** Expression of (a) *APOE,* (b) *APOC2,* and (c) *APOC4* in the Tabula Sapiens dataset for 14 cell types which correspond to cell types with snATAC-seq data used in lipids traits analyses using cellSTAAR. Log fold-changes ("logFC") shown are compared against the other 13 cell types, the Differential Expression (DE) p-value was calculated using the FindMarkers function in the Seurat R package, and the percentage of cells in each cell type expressing the gene is included in the legend. All three genes shown have been previously associated with LDL cholesterol levels [3, 4], yet transcriptomic evidence provides mixed results regarding which gene is potentially causal. In hepatocytes, for example, gene *APOE* has the highest percentage of cells expressing. Although the percentage of hepatocytes expressing both *APOC2* and *APOC4* is lower, both of these genes exhibit a distinct hepatocyte-specific expression profile. By contrast, *APOE* is broadly expressed across a number of tissues that are not as relevant to lipids levels such as cardiac fibroblast cells and microglial cells.

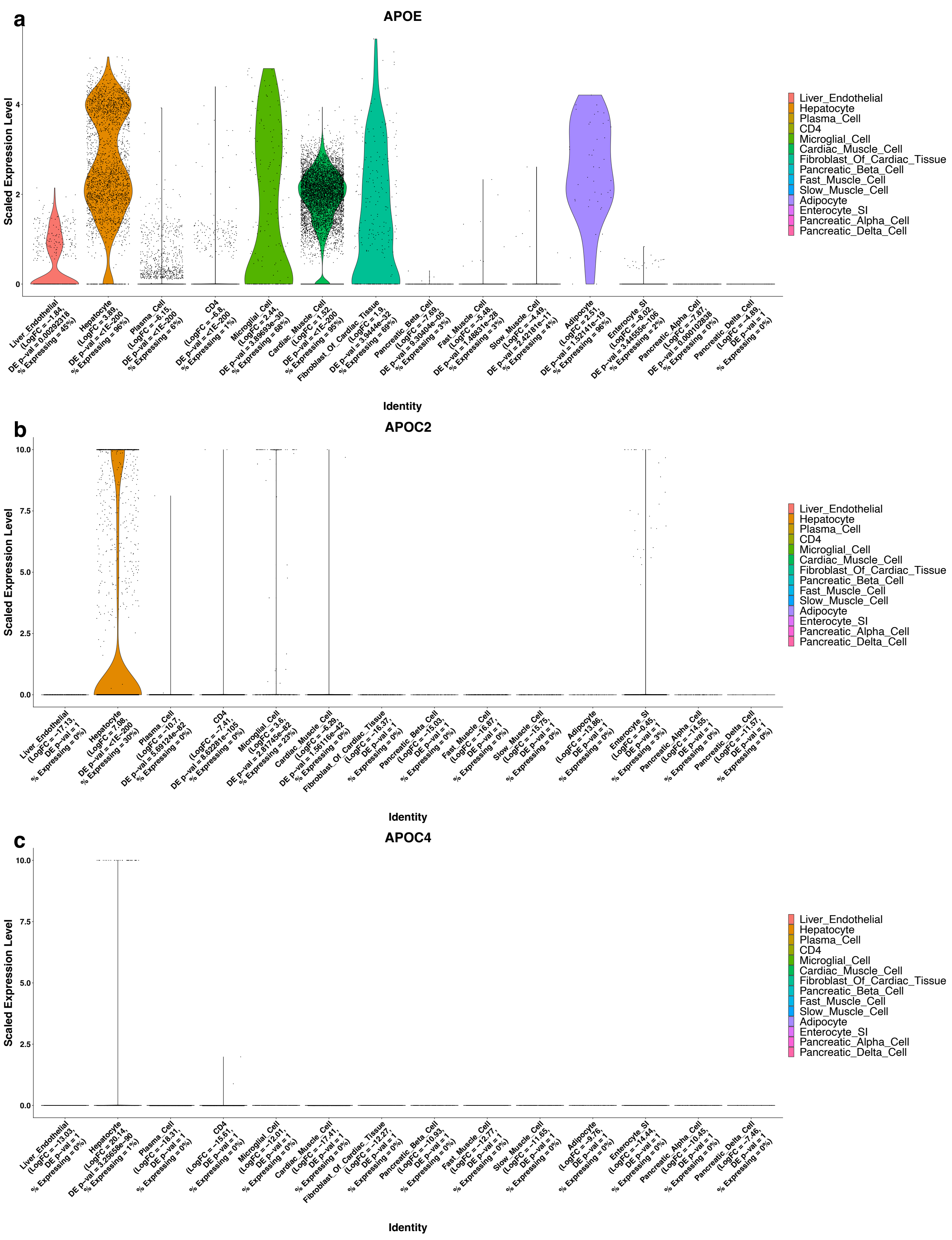
a
APOE
Scaled Expression Level
0
2
4
Identity
Liver_Endothelial
Hepatocyte
Plasma_Cell
CD4
Microglial_Cell
Cardiac_Muscle_Cell
Fibroblast_Of_Cardiac_Tissue
Pancreatic_Beta_Cell
Fast_Muscle_Cell
Slow_Muscle_Cell
Adipocyte
Enterocyte_SI
Pancreatic_Alpha_Cell
Pancreatic_Delta_Cell
b
APOC2
Scaled Expression Level
0.0
2.5
5.0
7.5
10.0
Identity
Liver_Endothelial
Hepatocyte
Plasma_Cell
CD4
Microglial_Cell
Cardiac_Muscle_Cell
Fibroblast_Of_Cardiac_Tissue
Pancreatic_Beta_Cell
Fast_Muscle_Cell
Slow_Muscle_Cell
Adipocyte
Enterocyte_SI
Pancreatic_Alpha_Cell
Pancreatic_Delta_Cell
c
APOC4
Scaled Expression Level
0.0
2.5
5.0
7.5
10.0
Identity
Liver_Endothelial
Hepatocyte
Plasma_Cell
CD4
Microglial_Cell
Cardiac_Muscle_Cell
Fibroblast_Of_Cardiac_Tissue
Pancreatic_Beta_Cell
Fast_Muscle_Cell
Slow_Muscle_Cell
Adipocyte
Enterocyte_SI
Pancreatic_Alpha_Cell
Pancreatic_Delta_Cell

**Supplementary Figure 18**: Discoveries (cellSTAAR p-value < $1x10^{-3}$) in enhancer regions (pELS and dELS regions as defined by ENCODE V3) visualized based on overlap across different linking approaches. All three continuous phenotypes (LDL-C, HDL-C, TG) and all 19 cell types are jointly shown. Shows that most discoveries are found using only a single linking approach.

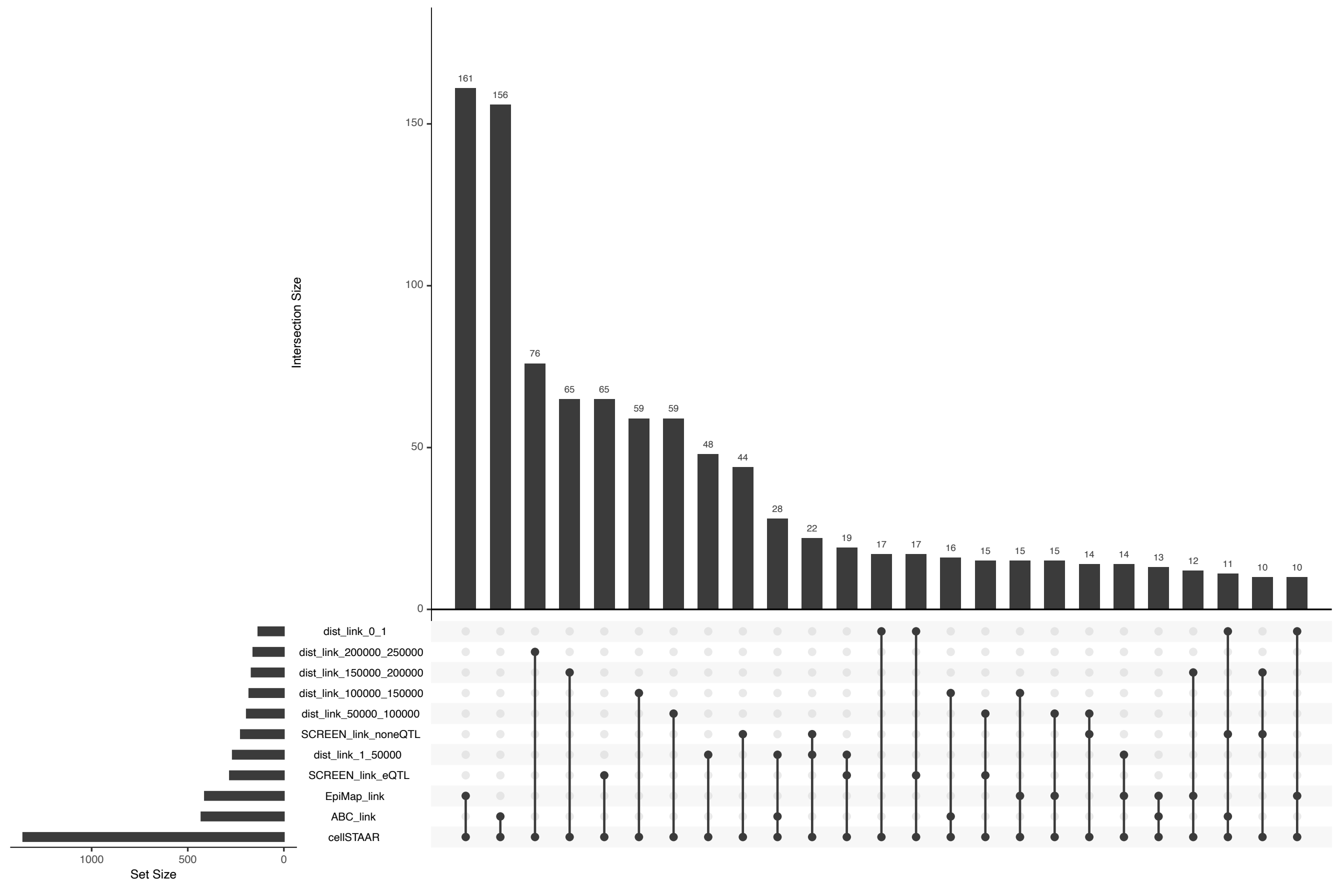

### Supplementary Note

##### TOPMed study participants and acknowledgements

Old Order Amish (OOA)

The Old Order Amish individuals included in this study were participants of several ongoing studies of cardiovascular health carried out at the University of Maryland among relatively healthy volunteers from the Old Order Amish community of Lancaster County, PA, and their family members[5, 6].

Whole genome sequencing (WGS) for the Trans-Omics in Precision Medicine (TOPMed) program was supported by the National Heart, Lung, and Blood Institute (NHLBI). WGS for “NHLBI TOPMed: Genetics of Cardiometabolic Health in the Amish” (phs000956.v1.p1) was performed at the Broad Institute of MIT and Harvard (3R01HL121007-01S1). The TOPMed component of the Amish Research Program was supported by NIH grants R01 HL121007, U01 HL072515, and R01 AG18728. Email Braxton Mitchell for additional input.

Atherosclerosis Risk in Communities Study (ARIC)

The ARIC study is a population-based prospective cohort study of cardiovascular disease sponsored by the National Heart, Lung, and Blood Institute (NHLBI). ARIC included 15,792 individuals, predominantly European American and African American, aged 45-64 years at baseline (1987-89), chosen by probability sampling from four US communities. Cohort members completed three additional triennial follow-up examinations, a fifth exam in 2011-2013, a sixth exam in 2016-2017, a seventh exam in 2018-2019, and eighth exam in 2020, a ninth exam in 2021-2022, and a tenth exam in 2023. The ARIC study has been described in detail previously [7].

Whole genome sequencing (WGS) for the Trans-Omics in Precision Medicine (TOPMed) program was supported by the National Heart, Lung, and Blood Institute (NHLBI). WGS for “NHLBI TOPMed: Atherosclerosis Risk in Communities (ARIC)” (phs001211) was performed at the Baylor College of Medicine Human Genome Sequencing Center (HHSN268201500015C and 3U54HG003273-12S2) and the Broad Institute of MIT and Harvard (3R01HL092577-

06S1). Centralized read mapping and genotype calling, along with variant quality metrics and filtering were provided by the TOPMed Informatics Research Center (3R01HL-117626-02S1; contract HHSN268201800002I). Phenotype harmonization, data management, sample-identity QC, and general study coordination, were provided by the TOPMed Data Coordinating Center (3R01HL-120393-02S1; contract HHSN268201800001I). We gratefully acknowledge the studies and participants who provided biological samples and data for TOPMed.

The Atherosclerosis Risk in Communities study has been funded in whole or in part with Federal funds from the National Heart, Lung, and Blood Institute, National Institutes of Health, Department of Health and Human Services, under Contract nos. (75N92022D00001, 75N92022D00002, 75N92022D00003, 75N92022D00004, 75N92022D00005). The authors thank the staff and participants of the ARIC study for their important contributions.

Mt Sinai BioMe Biobank (BioMe):

The Institute for Personalized Medicine at the Icahn School of Medicine at Mount Sinai is leading the movement toward diagnosis and classification of disease according to the patient's molecular profile. BioMe is the major effort towards this goal, BioMe, an electronic medical record-linked biobank that enables researchers to conduct genetic, epidemiologic, molecular, and genomic studies rapidly and efficiently on large collections of research specimens linked with medical information. BioMed cohort is composed of White, Black, Hispanic, and Asian populations. The Mount Sinai Medical Center services diverse local communities of upper Manhattan, including Central Harlem (86% African American), East Harlem (88% Hispanic Latino), and Upper East Side (88% Caucasian/white) with broad health disparities[8]. Whole genome sequencing (WGS) for the Trans-Omics in Precision Medicine (TOPMed) program was supported by the National Heart, Lung, and Blood Institute (NHLBI). The Mount Sinai BioMe Biobank has been supported by The Andrea and Charles Bronfman Philanthropies and in part by Federal funds from the NHLBI and NHGRI (U01HG00638001; U01HG007417; X01HL134588).

Coronary Artery Risk Development in Young Adults (CARDIA):

The Coronary Artery Risk Development in Young Adults (CARDIA) Study is a study examining the development and determinants of clinical and subclinical cardiovascular disease and their risk factors. It began in 1985-6 with a group of 5115 black and white men and women aged 18-30 years[9]. The participants were selected so that there would be approximately the same number of people in subgroups of race, gender, education (high school or less and more than high school) and age (18-24 and 25-30) in each of 4 centers: Birmingham, AL; Chicago, IL; Minneapolis, MN; and Oakland, CA. These same participants were asked to participate in follow-up examinations during 1987-1988 (Year 2), 1990-1991 (Year 5), 1992-1993 (Year 7), 1995-1996 (Year 10), 2000-2001 (Year 15), 2005-2006 (Year 20), 2010-2011 (Year 25), and 2015-2016 (Year 30).

The Coronary Artery Risk Development in Young Adults Study (CARDIA) is conducted and supported by the National Heart, Lung, and Blood Institute (NHLBI) in collaboration with the University of Alabama at Birmingham (75N92023D00002 & 75N92023D00005), Northwestern University (75N92023D00004), University of Minnesota (75N92023D00006), and Kaiser Foundation Research Institute (75N92023D00003).

Cleveland Family Study (CFS)

The CFS is a family-based longitudinal study that includes participants with laboratory diagnosed sleep apnea, their family members and neighborhood control families followed between 1990 and 2006. Four examinations over 16 years provided measurements of sleep apnea with overnight polysomnography, anthropometry, and other related phenotypes, as detailed previously[10, 11]. After an overnight fast, blood was collected which was assayed for lipid levels at the University of Vermont Laboratory for Clinical Biochemistry Research. Lipids (triglycerides, HDL cholesterol) from fasted blood serum were measured by enzymatic methods using Centers for Disease Control and Prevention guidelines[12].

Whole genome sequencing (WGS) for the Trans-Omics in Precision Medicine (TOPMed) program was supported by the National Heart, Lung, and Blood Institute (NHLBI). WGS for "NHLBI TOPMed: Cleveland Family Study" (phs000954) was performed at the University of Washington Northwest Genomics Center (3R01HL098433-05S1).

This research was supported by grants HL 046389; HL113338; 1R35HL135818 **from the National Heart, Lung, and Blood Institute (NHLBI)**.

Cardiovascular Health Study (CHS)

The Cardiovascular Health Study is a prospective population-based cohort study of risk factors for CHD and stroke in adults 65 years and older[13]. The main objective is to identify factors related to the onset and course of heart disease and stroke. The four Field Centers are located in Forsyth County, NC; Sacramento County, CA; Washington County, MD; and Pittsburgh, PA. The original cohort of 5201 elderly were recruited in 1989-1990; and in 1992-1993, 687 additional minority participants were recruited and examined. Each community sample was obtained from random samples of the Medicare eligibility lists of the Health Care Financing Administration (HCFA). Eligible to participate were persons living in the household of each sampled individual who were: 1) 65 yr or older; 2) non-institutionalized; 3) expected to remain in the area for 3 yr; and 4) able to give informed consent. Excluded were those wheelchair-bound, receiving hospice care or cancer treatment. The minority cohort was recruited using similar methods. Participants were eligible whether or not they had clinically apparent cardiovascular disease. Subjects were followed with semi-annual contacts, alternating between telephone calls and surveillance clinic visits.

Whole genome sequencing (WGS) for the Trans-Omics in Precision Medicine (TOPMed) program was supported by the National Heart, Lung, and Blood Institute (NHLBI). WGS for “NHLBI TOPMed: Cardiovascular Health Study” (phs001368) was performed at the Baylor College of Medicine Human Genome Sequencing Center (HHSN268201500015C). Analyses were limited to those with available DNA who consented to genetic studies.

**The Cardiovascular Health Study research was supported by NHLBI contracts HHSN268201200036C, HHSN268200800007C, HHSN268201800001C, N01HC55222, N01HC85079, N01HC85080, N01HC85081, N01HC85082, N01HC85083, N01HC85086, 75N92021D00006; and NHLBI grants U01HL080295, R01HL087652, R01HL105756, R01HL103612, R01HL120393, and U01HL130114 with additional contribution from the National Institute of Neurological Disorders and Stroke (NINDS). Additional support was provided through R01AG023629 from the National Institute on Aging (NIA). A full list of principal CHS investigators and institutions can be found at CHS-NHLBI.**

Diabetes Heart Study (DHS)

The Diabetes Heart Study (DHS) began as a family-based study enriched for type 2 diabetes (T2D). The initial cohort included 1443 European American and African American participants from 564 families with multiple cases of type 2 diabetes recruited between 1998 and 2006[14]. As an ancillary study, the African American Diabetes Heart Study (AA-DHS) expanded the total number of African Americans to 691 by recruiting additional unrelated participants with type 2 diabetes from 2007 and 2010[15]. All participants were extensively phenotyped for measures of subclinical CVD and other known CVD risk factors. Primary outcomes were quantified burden of vascular calcified plaque in the coronary artery, carotid artery, and abdominal aorta all determined from non-contrast computed tomography scans. For TOPMed, DHS and AA-DHS African American participants with CAC were selected for WGS, prioritizing the inclusion of families.

Whole genome sequencing (WGS) for the Trans-Omics in Precision Medicine (TOPMed) program was supported by the National Heart, Lung, and Blood Institute (NHLBI). WGS for “NHLBI TOPMed: Diabetes Heart Study” (phs001412) was performed at the Broad Institute of MIT and Harvard (HHSN268201500014C).

This work was supported by R01 HL92301, R01 HL67348, R01 NS058700, R01 AR48797, R01 DK071891, R01 AG058921, the General Clinical Research Center of the Wake Forest University School of Medicine (M01 RR07122, F32 HL085989), the American Diabetes Association, and a pilot grant from the Claude Pepper Older Americans Independence Center of Wake Forest University Health Sciences (P60 AG10484).

Framingham Heart Study (FHS)

The FHS is a three generational prospective cohort that has been described in detail previously[16]. Individuals were initially recruited in 1948 in Framingham, USA to evaluate cardiovascular disease risk factors. The second-generation cohort (5,124 offspring of the original cohort) was recruited between 1971 and 1975[17, 18]. The third-generation cohort (4,095 grandchildren of the original cohort) was collected between 2002 and 2005. Fasting lipid levels were measured at exam 1 of the Offspring (1971-1975) and third generation (2002-2005) cohorts, using standard LRC protocols.

Whole genome sequencing (WGS) for the Trans-Omics in Precision Medicine (TOPMed) program was supported by the National Heart, Lung, and Blood Institute (NHLBI). WGS for “NHLBI TOPMed: Whole Genome Sequencing and Related Phenotypes in the Framingham Heart Study” (phs000974.v1.p1) was performed at the Broad Institute of MIT and Harvard (HHSN268201500014C).

The Framingham Heart Study (FHS) acknowledges the support of contracts NO1-HC-25195, HHSN268201500001I and 75N92019D00031 from the National Heart, Lung and Blood Institute and grant supplement R01 HL092577-06S1 for this research. We also acknowledge the dedication of the FHS study participants without whom this research would not be possible. Dr. Vasan is supported in part by the Evans Medical Foundation and the Jay and Louis Coffman Endowment from the Department of Medicine, Boston University School of Medicine.

Genetic Study of Atherosclerosis Risk (GeneSTAR)

GeneSTAR is an ongoing family-based prospective study designed to determine environmental, phenotypic, and genetic causes of premature cardiovascular disease. GeneSTAR was originally conducted in healthy adult European- and African-American siblings of probands with documented early onset coronary disease under 60 years of age at the time of hospitalization in any of 10 Baltimore area hospitals from 1982-2006. Participants were screened for traditional coronary disease and stroke risk factors and have been followed regularly to ascertain incident cardiovascular disease[19]. Commencing in 2003, the siblings, their offspring, and the coparent of the offspring who were free of cardiovascular disease participated in a 2 week trial of aspirin 81 mg/day with pre and post ex vivo platelet function assessed using multiple agonists and were screened for traditional coronary disease and stroke risk factors[20]. Of the total 3949 participants, 1786 were selected for TOPMed prioritized on complete platelet function measures and largest family size.

Whole genome sequencing (WGS) for the Trans-Omics in Precision Medicine (TOPMed) program was supported by the National Heart, Lung, and Blood Institute (NHLBI). WGS for “NHLBI TOPMed: Genetic Study of Atherosclerosis Risk” (phs001218) was performed at Psomagen (formerly Macrogen; 3R01HL112064-04S1), Illumina (R01HL112064), and the Broad Institute of MIT and Harvard (HHSN268201500014C).

GeneSTAR was supported by grants from the National Institutes of Health/National Heart, Lung, and Blood Institute (U01 HL72518, HL087698, HL49762, HL59684, HL58625, HL071025, HL112064), by a grant from the National Institutes of Health/National Institute of Nursing Research (NR0224103), and by a grant from the National Institutes of Health/National Center for Research Resources (M01-RR000052) to the Johns Hopkins General Clinical Research Center.

Genetic Epidemiology Network of Arteriopathy (GENOA)

The Genetic Epidemiology Network of Arteriopathy (GENOA) is one of four networks in the NHLBI Family-Blood Pressure Program (FBPP)[21]. GENOA's long-term objective is to elucidate the genetics of target organ complications of hypertension, including both atherosclerotic and arteriosclerotic complications involving the heart, brain, kidneys, and peripheral arteries[22]. The longitudinal GENOA Study recruited European-American and African-American sibships with at least 2 individuals with clinically diagnosed essential hypertension before age 60 years. All other members of the sibship were invited to participate regardless of their hypertension status. Participants were diagnosed with hypertension if they had either 1) a previous clinical diagnosis of hypertension by a physician with current anti-hypertensive treatment, or 2) an average systolic blood pressure ≥ 140 mm Hg or diastolic blood pressure ≥ 90 mm Hg based on the second and third readings at the time of their clinic visit. Only participants of the African-American Cohort were sequenced through TOPMed.

During the first exam (Phase 1; 1996-2000), 1,583 European-Americans from Rochester, MN and 1,854 African-Americans from Jackson, MS were examined. Between 2000 and 2004 (Phase 2), 1,241 participants of the European-American Cohort and 1,482 participants of the African-American cohort returned for a second examination. The second examination of the European-American cohort included computed tomography scans for coronary artery calcification while the second examination of the African-American cohort included an echocardiogram. Between 2009 and 2011, an examination that included computed tomography scans for coronary artery calcification (CAC Study) was conducted on 752 participants of the African-American Cohort.

Every participant with an echocardiogram was selected for whole genome sequencing (WGS) through TOPMed. We then selected 106 African-American participants who had a computed tomography scan for coronary artery calcification but not an echocardiogram or were a sibling of someone already selected for WGS. Finally, we excluded individuals whom we knew were already being whole genome sequenced through TOPMed or another sequencing effort (GENOA participants who overlap with ARIC or JHS participants).

Support for GENOA was provided by the National Heart, Lung, and Blood Institute (U10HL054457, U01HL054464, U01HL054481, R01HL119443, R01HL085571, and R01HL087660) of the National Institutes of Health. DNA extraction for “NHLBI TOPMed: Genetic Epidemiology Network of Arteriopathy” (phs001345) was performed at the Mayo Clinic Genotyping Core, and WGS was performed at the DNA Sequencing and Gene Analysis Center at the University of Washington (3R01HL055673-18S1) and the Broad Institute (HHSN268201500014C). We would like to thank the GENOA participants.

Genetic Epidemiology Network of Salt Sensitivity (GenSalt)

GenSalt utilizes a family feeding-study design. Each family is ascertained through a proband with untreated prehypertension or stage-1 hypertension in rural China. Medical history, lifestyle risk factors, and cold pressor tests are obtained at baseline visits while BP, weight, blood and urine specimens are collected at baseline and follow-up visits[23]. The dietary intervention includes a 7-day low sodium-feeding (51.3 mmol/day), a 7-day high sodium-feeding (307.8 mmol/day), and a 7-day high sodium-feeding with an oral potassium supplementation (60 mmol/day). Whole genome sequencing (WGS) for the Trans-Omics in Precision Medicine (TOPMed) program was supported by the National Heart, Lung, and Blood Institute (NHLBI). GenSalt was supported by research grants (U01HL072507, R01HL087263, and R01HL090682) from the National Heart, Lung and Blood Institute, National Institutes of Health, Bethesda, MD.

Genetics of Lipid Lowering Drugs and Diet Network (GOLDN)

GOLDN is a family-based study of European descent individuals recruited in Minneapolis and Salt Lake City (two of the NHLBI Family Heart Study sites). It aims to uncover genetic predictors of variability in lipid phenotypes, which include both fasting and postprandial lipids quantified using traditional methods, NMR, and high-throughput lipidomics. During the initial screening of ~1,350 individuals, the following criteria were used for exclusion: age < 18 years; fasting triglycerides ≥1500 mg/dL; recent history of myocardial infarction, coronary bypass surgery, or coronary angioplasty; self-report of a positive history of liver, kidney, pancreas, or gallbladder disease, or a history of nutrient malabsorption; current use of insulin; abnormal liver or kidney function; in women of childbearing potential, pregnancy, breastfeeding, not using an acceptable form of contraception. Of those who enrolled, 1,048 individuals consented to the use of their DNA in research; 893 participants with data on all exposures, outcomes, and covariates were included in the current study.

GOLDN biospecimens, baseline phenotype data, and intervention phenotype data were collected with funding from National Heart, Lung, and Blood Institute (NHLBI) grant U01 HL072524. Whole-genome sequencing in GOLDN was funded by NHLBI grant R01 HL104135-04S1.

Whole genome sequencing (WGS) for the Trans-Omics in Precision Medicine (TOPMed) program was supported by the National Heart, Lung, and Blood Institute (NHLBI). WGS for “NHLBI TOPMed: Genetics of Lipid Lowering Drugs and Diet Network” (phs001359) was performed at the University of Washington Northwest Genomics Center (3R01HL104135-04S1).

<u>Hispanic Community Health Study - Study of Latinos (HCHS/SOL)</u>

The Hispanic Community Health Study (HCHS)/Study of Latinos (SOL) is a multicenter, community-based cohort study of Hispanic/Latino adults in the United States. The main goal of the study is to identify risk factors which could either be protective or harmful to the Hispanic community[24, 25]. A total of 16,415 Hispanic/Latino individuals of age 18-74 were recruited. Participants are recruited in community areas surrounding four field centers in the Bronx, Chicago, Miami, and San Diego. Whole genome sequencing (WGS) for the Trans-Omics in Precision Medicine (TOPMed) program was supported by the National Heart, Lung, and Blood Institute (NHLBI). The Hispanic Community Health Study/Study of Latinos was carried out as a collaborative study supported by contracts from the National Heart, Lung, and Blood Institute (NHLBI) to the University of North Carolina (N01-HC65233), University of Miami (N01-HC65234), Albert Einstein College of Medicine (N01-HC65235), Northwestern University (N01-HC65236), and San Diego State University (N01-HC65237).

Hypertension Genetic Epidemiology Network and Genetic Epidemiology Network of Arteriopathy (HyperGEN)

The Hypertension Genetic Epidemiology Network Study (HyperGEN) - Genetics of Left Ventricular (LV) Hypertrophy is a familial study aimed to understand genetic risk factors for LV hypertrophy by conducting genetic studies of continuous traits from echocardiography exams[26, 27]. As part of HyperGEN study, four field centers recruited African American and white hypertensive siblings aged 23 to 87 years. Data from detailed clinical exams as well as genotyping data for linkage studies, candidate gene studies and GWAS have been collected and is shared between HyperGEN and the ancillary HyperGEN - Genetics of LV Hypertrophy study. Whole genome sequencing (WGS) for the Trans-Omics in Precision Medicine (TOPMed) program was supported by the National Heart, Lung, and Blood Institute (NHLBI). The HyperGEN Study is part of the National Heart, Lung, and Blood Institute (NHLBI) Family Blood Pressure Program; collection of the data represented here was supported by grants U01 HL054472 (MN Lab), U01 HL054473 (DCC), U01 HL054495 (AL FC), and U01 HL054509 (NC FC). The HyperGEN: Genetics of Left Ventricular Hypertrophy Study was supported by NHLBI grant R01 HL055673 with whole-genome sequencing made possible by supplement - 18S1.

Jackson Heart Study (JHS)

The JHS is a large, population-based observational study evaluating the etiology of cardiovascular, renal, and respiratory diseases among African Americans residing in the three counties (Hinds, Madison, and Rankin) that make up the Jackson, Mississippi metropolitan area[28, 29]. Data and biologic materials have been collected from 5,306 participants, including a nested family cohort of 1,498 members of 264 families. The age at enrollment for the unrelated cohort was 35-84 years; the family cohort included related individuals >21 years old. Participants provided an extensive medical and social history and had an array of physical and biochemical measurements and diagnostic procedures, and a subset of participants provided genomic DNA during a baseline examination (2000-2004) and two follow-up examinations (2005-2008 and 2009-2012), with a fourth examination ongoing. Annual follow-up interviews and cohort surveillance are ongoing.

Whole genome sequencing (WGS) for the Trans-Omics in Precision Medicine (TOPMed) program was supported by the National Heart, Lung, and Blood Institute (NHLBI). WGS for "NHLBI TOPMed: The Jackson Heart Study" (phs000964.v1.p1) was performed at the University of Washington Northwest Genomics Center (HHSN268201100037C).

The Jackson Heart Study (JHS) is supported and conducted in collaboration with Jackson State University (HHSN268201800013I), Tougaloo College (HHSN268201800014I), the Mississippi State Department of Health (HHSN268201800015I/HHSN26800001) and the University of Mississippi Medical Center (HHSN268201800010I, HHSN268201800011I and HHSN268201800012I) contracts from the National Heart, Lung, and Blood Institute (NHLBI) and the National Institute for Minority Health and Health Disparities (NIMHD). The authors also wish to thank the staffs and participants of the JHS.

Multi-Ethnic Study of Atherosclerosis (MESA)

The Multi-Ethnic Study of Atherosclerosis is a National Heart, Lung and Blood Institute-sponsored, population-based investigation of subclinical cardiovascular disease and its progression[30]. A total of 6,814 individuals, aged 45 to 84 years, were recruited from six US communities (Baltimore City and County, MD; Chicago, IL; Forsyth County, NC; Los Angeles County, CA; New York, NY; and St. Paul, MN) between July 2000 and August 2002. Participants were excluded if they had physician-diagnosed cardiovascular disease prior to enrollment, including angina, myocardial infarction, heart failure, stroke, or TIA, resuscitated cardiac arrest or a cardiovascular intervention (e.g., CABG, angioplasty, valve replacement, or pacemaker/defibrillator placement). Pre-specified recruitment plans identified four racial/ethnic groups (White European-American, African-American, Hispanic-American, and Chinese-American) for enrollment, with targeted oversampling of minority groups to enhance statistical power.

Whole genome sequencing (WGS) for the Trans-Omics in Precision Medicine (TOPMed) program was supported by the National Heart, Lung, and Blood Institute (NHLBI). WGS for "NHLBI TOPMed: Multi-Ethnic Study of Atherosclerosis (MESA)" (phs001416.v3.p1) was performed at the Broad Institute of MIT and Harvard (3U54HG003067-13S1). Centralized read mapping and genotype calling, along with variant quality metrics and filtering were provided by the TOPMed Informatics Research Center (3R01HL-117626-02S1). Phenotype harmonization, data management, sample-identity QC, and general study coordination, were provided by the TOPMed Data Coordinating Center (3R01HL-120393-02S1), and TOPMed MESA Multi-Omics (HHSN2682015000031/HSN26800004). The MESA projects are conducted and supported by the National Heart, Lung, and Blood Institute (NHLBI) in collaboration with MESA investigators. Support for the Multi-Ethnic Study of Atherosclerosis (MESA) projects are conducted and supported by the National Heart, Lung, and Blood Institute (NHLBI) in collaboration with MESA investigators. Support for MESA is provided by contracts 75N92020D00001, HHSN268201500003I, N01-HC-95159, 75N92020D00005, N01-HC-95160, 75N92020D00002, N01-HC-95161, 75N92020D00003, N01-HC-95162, 75N92020D00006, N01-HC-95163, 75N92020D00004, N01-HC-95164, 75N92020D00007, N01-HC-95165, N01-HC-95166, N01-HC-95167, N01-HC-95168, N01-HC-95169, UL1-TR-000040, UL1-TR-001079, UL1-TR-001420, UL1TR001881, DK063491, and R01HL105756. The authors thank the other investigators, the staff, and the participants of the MESA study for their valuable contributions. A full list of participating MESA investigators and institutes can be found at http://www.mesa-nhlbi.org.

San Antonio Family Heart Study (SAFS)

The SAFHS began in 1991 and included 1,431 individuals in 42 extended families at baseline. Probands were 40- to 60-year-old low-income Mexican Americans selected at random without regard to presence or absence of disease, almost exclusively from Mexican American census tracts in San Antonio, Texas. All first-, second-, and third-degree relatives of the proband and of the proband's spouse, aged 16 years or above, were eligible to participate in the study. As part of our ongoing studies, we have recruited new family members from the original families, expanding the cohort to almost 3,099 individuals primarily from 73 families. Our study is a mixed longitudinal design. Subjects have been seen between 1 and 4 times with an average of 1.95 examinations.

Whole genome sequencing (WGS) for the Trans-Omics in Precision Medicine (TOPMed) program was supported by the National Heart, Lung, and Blood Institute (NHLBI). WGS for "NHLBI TOPMed: San Antonio Family Heart Study" (phs001215) was performed at the Illumina Genomic Services (3R01HL113323-03S1).

Collection of the San Antonio Family Study data was supported in part by National Institutes of Health (NIH) grants R01 HL045522, MH078143, MH078111 and MH083824; and whole genome sequencing of SAFS subjects was supported by U01 DK085524 and R01 HL113323. We are very grateful to the participants of the San Antonio Family Study for their continued involvement in our research programs.

<u>Genome-wide Association Study of Adiposity in Samoans (Samoan)</u>

The parent Samoan study is a population-based genome-wide association study (GWAS) of adiposity and cardiometabolic phenotypes among adults, 25-65 years of age, from the independent nation of Samoa in the South Pacific. The research goal of this study is to identify genetic variation that increases susceptibility to obesity and cardiometabolic phenotypes. Biomarker and questionnaire data were collected to assess cardiometabolic phenotypes. DNA was collected and the Affymetrix 6.0 chip used for SNP genotyping. After quality control checks on genotyping and excluding individuals with key missing data we have a final sample of 3,122 adults with high-quality genome-wide marker data[31]. Participation in TOPMed provided whole genome sequence data for 1,285 individuals from the GWAS sample chosen for maximal informativity for our Samoan-specific imputation panel.

The Samoan study was supported by the National Institutes of Health grants R01 HL093093 (S.T. McGarvey) and R01 HL133040 (R.L. Minster). Molecular data for the Trans-Omics in Precision Medicine (TOPMed) program were provided by NHLBI. Genome sequencing for the Soifua Manuia study, labeled “NHLBI TOPMed: Genome-wide Association Study of Adiposity in Samoans” (phs000972.v5.p1) in dbGaP, was performed at the Northwest Genomics Center (HHSN268201100037C) and the New York Genome Center (HHSN268201500016C). Core support, including centralized genomic read mapping and genotype calling, along with variant quality metrics and filtering, was provided by the TOPMed Informatics Research Center (R01 HL117626 02S1; contract HHSN268201800002I). Core support, including phenotype harmonization, data management, sample-identity QC, and general program coordination, was also provided by the TOPMed Data Coordinating Center (R01 HL120393; U01 HL120393; contract HHSN268201800001I). We gratefully acknowledge the studies and participants who provided biological samples and data for TOPMed.

We thank the study participants, local field workers, local village authorities, and the Samoan government—particularly the Ministry of Health; the Ministry of Women, Community, and Social Development; the Office of the Prime Minister; and the Samoa Bureau of Statistics—for their continued support of The Samoan Obesity, Lifestyle and Genetic Adaptations (OLaGA) Study Group. The members of the OLaGA Study Group are: Ranjan Deka, Nicola L. Hawley, Ryan L. Minster, Take Naseri, Satupaʻitea Viali, Daniel E. Weeks, and Stephen T. McGarvey.

Taiwan Study of Hypertension using Rare Variants (THRV)

The THRV-TOPMed study consists of three cohorts: The SAPPHIRe Family cohort, TSGH (Tri-Service General Hospital, a hospital-based cohort), and TCVGH (Taichung Veterans General Hospital, another hospital-based cohort), all based in Taiwan[32, 33]. 1,271 subjects were previously recruited as part of the NHLBI-sponsored SAPPHIRe Network (which is part of the Family Blood Pressure Program, FBPP). THRV is a collaborative study between Washington University in St. Louis, LA BioMed at Harbor UCLA, University of Texas in Houston, Taichung Veterans General Hospital, Taipei Veterans General Hospital, Tri-Service General Hospital, National Health Research Institutes, National Taiwan University, and Baylor University. THRV is based (substantially) on the parent SAPPHIRe study, along with additional population-based and hospital-based cohorts. Whole genome sequencing (WGS) for the Trans-Omics in Precision Medicine (TOPMed) program was supported by the National Heart, Lung and Blood Institute (NHLBI). The Rare Variants for Hypertension in Taiwan Chinese (THRV) is supported by the National Heart, Lung, and Blood Institute (NHLBI) grant (R01HL111249) and its participation in TOPMed is supported by an NHLBI supplement (R01HL111249-04S1). SAPPHIRe was supported by NHLBI grants (U01HL54527, U01HL54498) and Taiwan funds, and the other cohorts were supported by Taiwan funds.

Women's Health Initiative (WHI)

The Women's Health Initiative (WHI) is a large study of postmenopausal women's health investigating risk factors for cancer, CVD, age-related fractures and chronic disease[34]. It began in 1993 as a set of randomized controlled clinical trials (CT) and an observational study (OS). Specifically, the CT (n=68,132) included three overlapping components: The Hormone Therapy (HT) Trials (n=27,347), Dietary Modification (DM) Trial (n=48,835), and Calcium and

Vitamin D (CaD) Trial (n=36,282). Eligible women could be randomized into as many as all three CTs components. Women who were ineligible or unwilling to join the CT were then invited to join the OS (n=93,676).

Whole genome sequencing (WGS) for the Trans-Omics in Precision Medicine (TOPMed) program was supported by the National Heart, Lung, and Blood Institute (NHLBI). WGS for "NHLBI TOPMed: Women's Health Initiative" (phs001237) was performed at the Broad Institute of MIT and Harvard (HHSN268201500014C).

The WHI program is funded by the National Heart, Lung, and Blood Institute, National Institutes of Health, U.S. Department of Health and Human Services through contracts 75N92021D00001, 75N92021D00002, 75N92021D00003, 75N92021D00004, 75N92021D00005. The authors thank the WHI investigators and staff for their dedication, and the study participants for making the program possible. A full listing of WHI investigators can be found at: http://www.whi.org/researchers/Documents%20%20Write%20a%20Paper/WHI%20Investigator%20Long%20List.pdf.

SHORT LIST OF WHI INVESTIGATORS **Program Office:** (National Heart, Lung, and Blood Institute, Bethesda, Maryland) Jacques Rossouw, Shari Ludlam, Joan McGowan, Leslie Ford, and Nancy Geller **Clinical Coordinating Center:** (Fred Hutchinson Cancer Research Center, Seattle, WA) Garnet Anderson, Ross Prentice, Andrea LaCroix, and Charles Kooperberg **Investigators and Academic Centers:** (Brigham and Women's Hospital, Harvard Medical School, Boston, MA) JoAnn E. Manson; (MedStar Health Research Institute/Howard University, Washington, DC) Barbara V. Howard; (Stanford Prevention Research Center, Stanford, CA) Marcia L. Stefanick; (The Ohio State University, Columbus, OH) Rebecca Jackson; (University of Arizona, Tucson/Phoenix, AZ) Cynthia A. Thomson; (University at Buffalo, Buffalo, NY) Jean Wactawski-Wende; (University of Florida, Gainesville/Jacksonville, FL) Marian Limacher; (University of Iowa, Iowa City/Davenport, IA) Jennifer Robinson; (University of Pittsburgh, Pittsburgh, PA) Lewis Kuller; (Wake Forest University School of Medicine, Winston-Salem, NC) Sally Shumaker; (University of Nevada, Reno, NV) Robert Brunner Women's Health Initiative Memory Study: (Wake Forest University School of Medicine, Winston-Salem, NC) Mark Espeland

The content is solely the responsibility of the authors and does not necessarily represent the official views of the National Institutes of Health.
